# Intracellular complement Factor B couples inflammatory sensing to metabolism, chromatin regulation and maladaptive tissue repair

**DOI:** 10.64898/2026.10.04.756581

**Authors:** Tilo Freiwald, Daniel Chauss, Md Tajmul, Jorge Luis Trujillo-Ochoa, Dhaneshwar Kumar, Jianping Ma, Madeline A Snyder, Nardos T. Cheru, Julius FE Jaeger, Yuxin Zhuang, Justin B. Gregor, Akshatha Ganne, Luopin Wang, Yubing Guo, Maja Gunkel, Tobias B. Huber, Thorsten Wiech, Erin E West, Srinivas Ayyadevara, Martin Kolev, Scott D. Briggs, Claudia Kemper, Majid Kazemian, Didier Portilla, Behdad Afzali

**Author notes:** co-equal contribution (Daniel Chauss, Md Tajmul). Joint last authors (Majid Kazemian, Didier Portilla, Behdad Afzali).

## Abstract

Inflammatory signals are broadly conserved, yet how tissue context converts them into programs of recovery or maladaptive repair remains unclear. Using kidney injury as a model of inflammation-to-repair transitions, we identify an injury-induced proximal-tubule complement program in which cell-intrinsic complement proteins are repurposed across subcellular compartments to couple inflammation to metabolic and chromatin regulation and promote maladaptive fibrotic remodeling. Mechanistically, myeloid-derived IL-1β signals through IL-1R1 on injured tubular cells to recruit NF-κB and AP-1 to *C3* and *Cfb*, coordinately inducing complement component 3 (C3) and Factor B (FB). Cytoplasmic FB drives cleavage-activation of cell-intrinsic C3, which supports tubular cell mitochondrial metabolic adaptation to stress and accumulation of C3aR-expressing macrophages around injured tubules. Nuclear FB, on the other hand, binds chromatin through a previously unrecognized DNA-binding interface and mediates histone H3 proteolysis and the preferential remodeling of chromatin promotors enriched in transcriptional programs relevant to maladaptive repair. Key features of this injury-associated C3-FB response, including intranuclear FB, are conserved in human kidney disease, associate with kidney dysfunction, and distinguish diseased from healthy kidneys. Together, these findings suggest that extracellular complement-targeting therapies may miss disease-relevant intracellular functions and suggest nuclear FB as a distinct target whose chromatin interaction and processing capacity could selectively limit maladaptive tissue responses beyond conventional inhibition.

**Graphical abstract:** 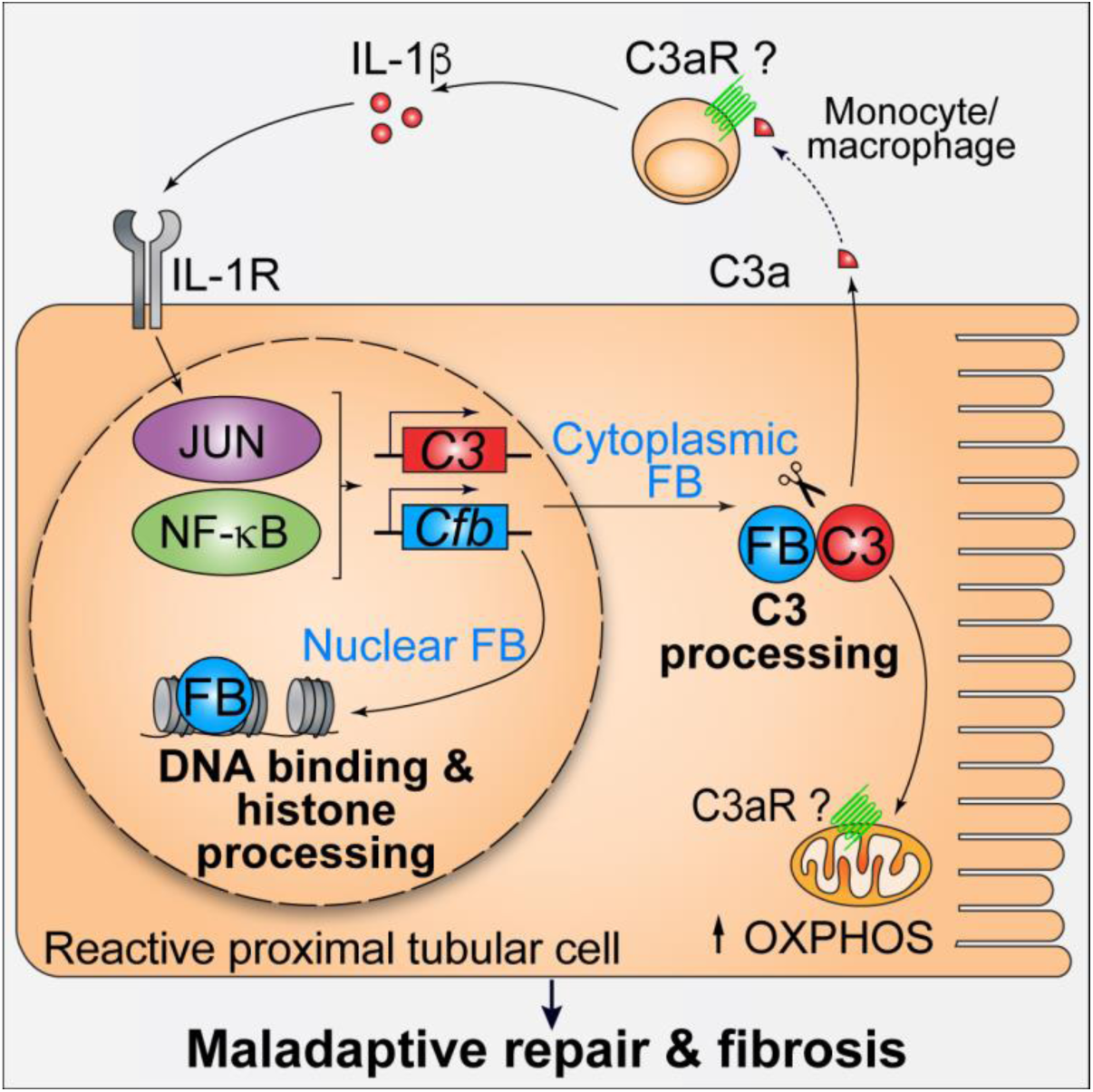

- Tissue injury induces coordinated C3-Factor B expression through myeloid IL-1β, IL-1R1, NF-κB and AP-1
- Cytoplasmic FB regulates intracellular C3 processing, supporting mitochondrial respiratory adaptation to stress
- Nuclear FB binds chromatin, promotes histone H3 processing and remodels promoter chromatin
- The tubular C3-Factor B program promotes maladaptive repair and fibrosis, with key features conserved in human kidney disease

## Introduction

Tissue-resident structural and immune cells cooperate to coordinate local inflammatory and repair responses that eliminate damage while preserving specialized functions of the affected organ^1,2^. Although many inflammatory pathways are broadly conserved across tissues, their effects are profoundly shaped by the identity and physiological state of the responding cells. Local inflammatory signals can engage tissue-intrinsic programs that promote recovery or, when sustained or dysregulated, result in maladaptive remodeling and fibrosis^3,4^. However, how these inflammatory pathways are translated into tissue-and cell-specific outcomes remains poorly understood.

The complement system is an evolutionarily ancient arm of innate immunity that detects pathogens and altered self and coordinates inflammatory responses^5^. In its canonical extracellular form, liver-derived complement proteins circulate in plasma, where activation through the classical, lectin or alternative pathways converges on proteolytic processing of complement component 3 (C3). Complement Factor B (FB), the catalytic component of the alternative pathway C3 convertase, associates with C3b and, following cleavage by Factor D, forms the proteolytically active C3bBb convertase, thereby amplifying C3 activation and generating inflammatory and effector products including C3a, C3b and, downstream, C5a and the membrane attack complex^6,7^. C3a and C5a act through their cognate receptors, C3aR and C5aR1, to promote leukocyte recruitment and inflammatory activation^6^.

Beyond this canonical circulating system, complement produced locally by immune cells and tissue-resident structural cells, including epithelial cells and fibroblasts, has emerged as an integral regulator of tissue immunity and inflammation. Locally synthesized complement can act through paracrine or autocrine pathways or function intracellularly, where it regulates various aspects of cellular physiology including survival, differentiation and metabolism^8,9^. In immune cells, we have previously shown that T cells synthesize and process C3 intracellularly, with cell-intrinsic complement signaling being essential for T cells to assume full inflammatory function in tissues^10,11^. In structural cells, synovial fibroblasts and respiratory epithelial cells similarly synthesize and process C3 intracellularly to sustain local inflammatory responses and cell survival^12–15^. These findings show that cell-intrinsic C3 functions across both immune and non-immune compartments, with its effects shaped by the identity and physiological context of the responding cell^16–18^.

A key question is whether this intracellular repurposing extends beyond C3 to other complement proteins that normally act with C3 in the extracellular cascade. FB is particularly compelling because its canonical catalytic function is closely linked to C3 activation in the alternative pathway, yet its intracellular biology remains largely unexplored. Whether FB simply supports intracellular C3 processing or acquires autonomous cell-intrinsic functions is unknown. This distinction has become increasingly relevant with the clinical development of proximal complement inhibitors, including therapies targeting C3 and FB^19,20^, because disease-relevant intracellular functions may require distinct target engagement and could expose opportunities for function-selective therapeutic intervention.

Here, using an integrated functional-genomic approach spanning single-cell analysis, cell-specific genetics, structural modelling, biochemistry and chromatin profiling, we identify locally produced complement as an important regulator of maladaptive tissue repair in the kidney and uncover an unexpected non-canonical role for FB in direct chromatin regulation. We define an injury-induced proximal-tubule complement response in which inflammatory IL-1β signaling recruits NF-κB and AP-1 to regulatory elements at the *C3* and *Cfb* loci, inducing both genes. Once expressed, however, C3 and FB assume distinct functions across subcellular compartments. In the cytoplasm, FB promotes processing of intracellularly synthesized C3, which supports mitochondrial metabolic adaptation. In the nucleus, FB binds chromatin through an unexpected DNA-binding interface and supports histone H3 processing, promoter chromatin remodeling and expression of associated transcriptional programs. *In vivo*, injury-triggered tubular C3 promotes the transition from acute injury to maladaptive fibrotic repair, shaping macrophage accumulation and state, with conditional deletion attenuating subsequent fibrosis without altering the severity of the initial injury. In human kidney disease, coordinated C3 and FB expression is conserved in injury-associated proximal tubular cells, together with the associated IL-1R-NF-κB/AP-1 transcriptional signature. Higher renal C3 and FB expression is linked to worse kidney function, distinguishes diseased from healthy renal tissue, and is particularly prominent in ANCA-associated vasculitis. Together, these findings reveal how canonical complement proteins can be locally induced within tissue-resident structural cells and repurposed across subcellular compartments to couple inflammatory signals to metabolic adaptation, chromatin regulation and tissue repair.

## Results

### Kidney injury induces a C3-FB complement program in reactive proximal tubular cells

To identify transcriptional programs governing specialized tissue biology, we first analyzed an RNA-seq dataset encompassing 13 tissues from perfused healthy mice (**Figure 1A**; E-MTAB-6081)^21^. We used principal-component analysis (PCA) to identify the major axes of transcriptional variation between these tissues and asked which biological processes contributed most strongly to the top three principal components (PCs) by performing pathway enrichment analysis on genes associated with each PC (**Figure 1B**). Together, PC1-PC3 accounted for 92.9% of total transcriptional variance. PC1 was dominated by metabolic pathways, whereas cell-surface and ligand-interaction pathways contributed prominently to PC2. Strikingly, complement-and coagulation-related pathways were among the strongest contributors to PC3. We next asked to what extent complement-gene expression alone could distinguish tissues. PCA restricted to complement components and regulators revealed a distinct tissue distribution, with kidney and liver separating prominently from the other tissues **(Figure 1C**). Consistent with this, expression of individual complement components and regulators showed a distinct pattern in kidney compared with other tissues (**Figure S1A**), and pairwise correlation across complement-pathway genes grouped kidney samples separately from the other tissues (**Figure S1B)**. These analyses suggest that the healthy kidney has a distinctive complement transcriptional program relative to other tissues under baseline conditions.

**Figure 1.**
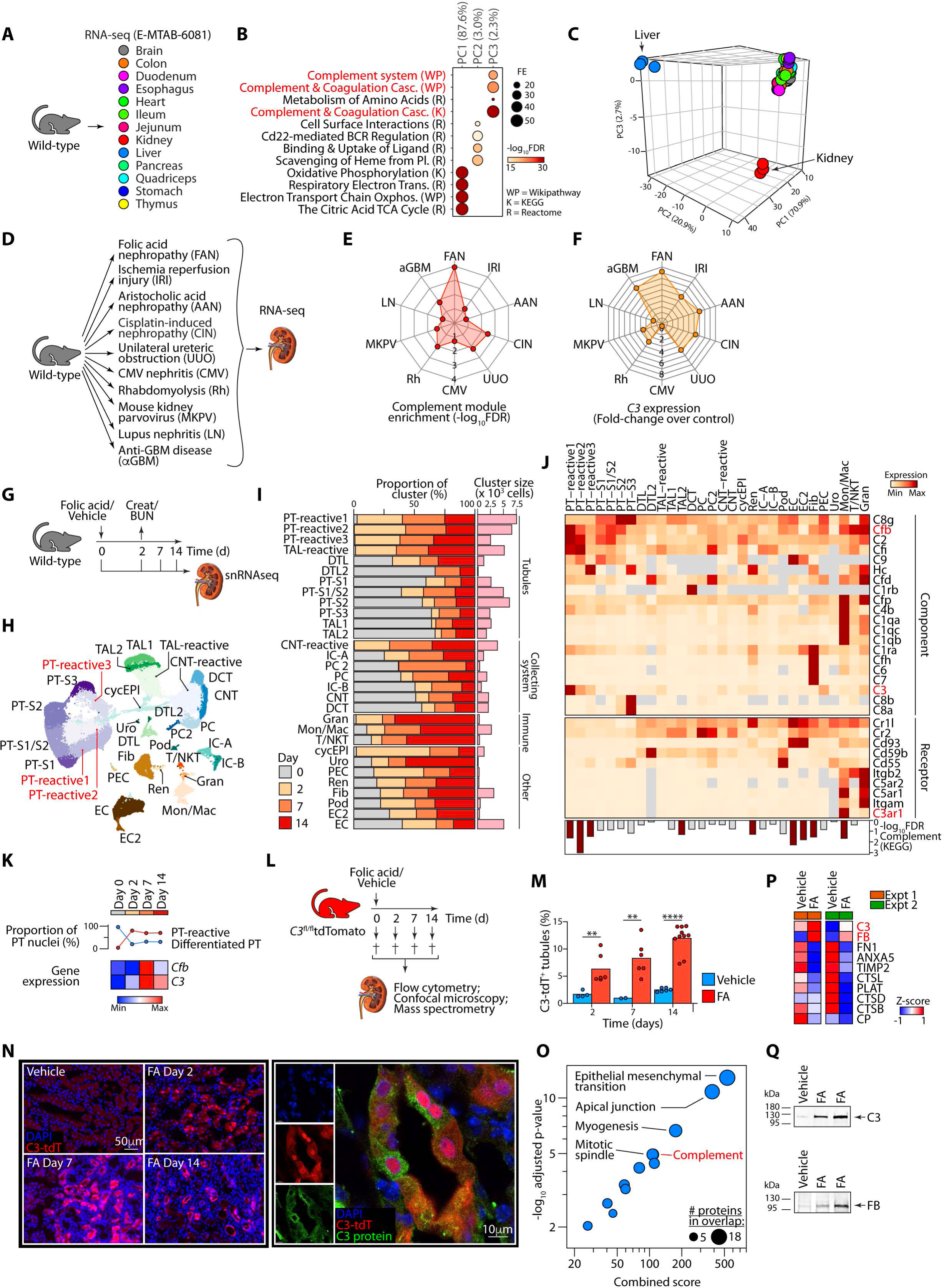
Kidney injury induces a C3-CFB complement program in reactive proximal tubular epithelial cells. A, Schematic of RNA-seq analysis of 13 perfused tissues from wild-type mice using the E-MTAB-6081 dataset. **B**, Pathway enrichment analysis of genes contributing to the first three principal components (PCs) of transcriptional variation across tissues. Bubble size indicates fold enrichment (FE), and color indicates statistical significance. Percentages indicate the proportion of total transcriptional variance explained by each PC; PC1-PC3 together accounted for 92.9% of the variance. **C**, PCA of the 13 mouse tissues based specifically on expression of complement-pathway genes. Kidney and liver samples are highlighted. **D** Schematic of publicly available mouse kidney injury transcriptomic datasets used to examine complement responses across experimental renal diseases. FAN, folic acid nephropathy; IRI, ischemia-reperfusion injury; AAN, aristolochic acid nephropathy; CIN, cisplatin-induced nephropathy; UUO, unilateral ureteric obstruction; CMV, cytomegalovirus nephritis; Rh, rhabdomyolysis; MKPV, mouse kidney parvovirus nephritis; LN, lupus nephritis; aGBM, anti-glomerular basement membrane disease. **E-F**, Radar plots showing KEGG complement pathway module enrichment (**E**) and *C3* expression relative to control (**F**) across the indicated kidney injury models. Where multiple time points were available, the highest enrichment score or *C3* expression value for each model is shown; complete time-course and enrichment data are shown in **Figure S2**. **G**, Experimental design for single-nucleus RNA-sequencing (snRNA-seq) of kidneys during FAN. **H**, UMAP visualization of kidney cell populations identified by snRNA-seq across the indicated time points. Three injury-associated proximal tubular (PT) populations are denoted PT-reactive1, PT-reactive2 and PT-reactive3. PT-S1, PT-S2 and PT-S3, proximal tubule segments 1, 2 and 3; TAL, thick ascending limb; DTL, descending thin limb; DCT, distal convoluted tubule; CNT, connecting tubule; PC, principal cell; IC-A, type A intercalated cell; IC-B, type B intercalated cell; Pod, podocyte; PEC, parietal epithelial cell; Ren, juxtaglomerular renin-producing granular cell; Fib, fibroblast; EC, endothelial cell; cycEPI, cycling epithelial cell; Gran, granulocyte; Mon/Mac, monocyte/macrophage; T/NKT, T/natural killer T cell; and Uro, urothelial cell. **I**, Composition of each kidney cell population according to sampling time point (left), with the total number of nuclei assigned to each population shown on the right. **J**, Heatmap showing expression of complement components and receptors across kidney cell populations. *C3*, *Cfb* and *C3ar1* are highlighted in red. Bars below indicate enrichment of the KEGG complement pathway. **K**, Proportion of proximal tubular nuclei assigned to differentiated PT or PT-reactive states over time (top) and expression of C3 and Cfb within proximal tubular populations during the injury-repair response (bottom). **L**, Experimental design for analysis of C3 expression using *C3*-tdTomato reporter mice expressing following vehicle or folic acid administration. **M**, Percentage of renal tubules containing C3-tdTomato^+^ cells at the indicated time points after vehicle or folic acid administration. Each point represents an individual mouse. **N**, Representative kidney sections showing C3-tdTomato expression after vehicle or folic acid administration (left) and co-localization of C3-tdTomato with C3 protein following injury (right). Scale bars, 50μm (left) and 10 μm (right). **O**, Pathway enrichment analysis of proteins differentially abundant in cell-culture supernatants from primary tubular epithelial cells following FA treatment. Bubble size indicates the number of proteins overlapping each pathway; the x axis shows the combined enrichment score and the y axis shows statistical significance. Complement and selected epithelial injury-associated pathways are indicated. **P**, Heatmap showing selected differentially abundant proteins in cell-culture supernatants following FA treatment in two independent primary tubular-cell preparations. C3 and FB are highlighted. **Q**, Representative immunoblots of cell-culture supernatants from primary tubular epithelial-cell preparations following vehicle or FA treatment, showing increased C3 and FB abundance. Bars show mean ± s.e.m. where indicated. **p<0.01, ****p<0.0001. snRNAseq data in Figure 1 are from n=1 vehicle and n=2 folic acid treated mice per timepoint.

Since the cellular sources and molecular functions of locally produced complement in kidneys remain incompletely understood^17,22,23^, we next asked whether the kidney complement program identified in healthy tissue was altered during distinct kidney injury models. We analyzed publicly available renal transcriptomes spanning multiple experimental models of kidney disease, including folic acid nephropathy (FAN), ischemia-reperfusion injury (IRI), unilateral ureteric obstruction (UUO), cisplatin nephrotoxicity (CIN), aristolochic acid nephropathy (AAN), rhabdomyolysis (Rh), viral nephritis, and immune-mediated kidney injury (**Figures 1D**). Complement-pathway enrichment was observed across multiple forms of renal injury but varied substantially between models and over time, with FAN showing the highest enrichment (**Figure 1E** and **S2**). FAN was also characterized by marked induction of *C3* (**Figure 1F** and **S2**), encoding the central substrate of the complement pathway. We therefore selected FAN as an experimental model to define the cellular origin and function of the injury-associated complement response.

To resolve this response at cellular resolution, we performed single-nucleus RNA sequencing of kidneys at days 0, 2, 7 and 14 after folic acid (FA) administration (**Figure 1G**). Unsupervised clustering and annotation using established lineage markers identified the major tubular, endothelial, stromal and immune populations of the kidney, together with several injury-associated epithelial states (**Figures 1H** and **S3A**). Among these were three proximal tubular populations that emerged following injury, which we termed PT-reactive1, PT-reactive2 and PT-reactive3. These populations expanded rapidly after FA injury and remained prominent during the subsequent injury-repair response, whereas differentiated proximal tubular populations (PT-S1, PT-S2 and PT-S3) contracted (**Figure 1I** and **S3B**).

The complement program showed a degree of cell-type specificity within the kidney. *C3* expression was most prominent within PT-reactive populations, whereas receptors for C3-derived activation fragments showed a distinct distribution: *C3ar1* and the complement receptor 3 (CR3) components *Itgam* and *Itgb2*, were concentrated predominantly in myeloid cells (**Figure 1J** and **S3C**). We noted that *Cfb*, encoding the catalytic component of the alternative pathway C3 convertase showed a broader and distinctive pattern. *Cfb* was also expressed most highly by PT-reactive cells and selected myeloid populations, but lower-level expression was also evident in differentiated healthy proximal tubules cells (**Figures 1J** and **S3C**). The prominent co-expression of *C3* and *Cfb* within PT-reactive populations was notable. Across kidney cell populations, expression of the two genes frequently co-varied, with the three PT-reactive populations showing the highest expression levels (**Figure S3D**). Temporally, both *C3* and *Cfb* increased within proximal tubular cells after injury, peaked at day 7, and remained elevated above baseline at day 14 (**Figure 1K**), consistent with coordinated induction during the injury-repair response.

We next examined tubular *C3* induction independently using a *C3*-tdTomato reporter mouse^10^ subjected to the same injury time course (**Figure 1L**). C3 reporter-positive tubules increased by day 2 following FA administration and became progressively more abundant at days 7 and 14 (**Figure 1M**). Confocal microscopy localized reporter-positive cells to the tubular epithelium and demonstrated intracellular C3 protein within tdTomato^+^ tubular cells (**Figure 1N**). These data suggest that injury-induced tubular *C3* transcription was accompanied by accumulation of C3 protein within renal epithelial cells.

Finally, to determine whether the injury-associated complement transcriptional response leads to release of complement proteins from tubular cells, we performed LC-MS/MS proteomic analysis of concentrated cell-culture supernatants from primary mouse kidney tubular cells treated *in vivo* with vehicle or folic acid (FA). FA broadly altered the composition of proteins released into the culture medium, with pathway enrichment identifying complement among the significantly enriched biological programs, alongside pathways associated with epithelial injury and remodeling (**Figure 1O**). C3 and FB were among the proteins significantly increased following FA treatment (**Figures 1P and S4**). Immunoblotting of cell-culture supernatants independently confirmed increased abundance of C3 and FB following FA exposure (**Figure 1Q**).

Together, these complementary transcriptomic, reporter, imaging and proteomic analyses identify reactive proximal tubular epithelial cells as a major source of an injury-inducible, coordinated C3-FB complement program in the kidney.

### C3 marks heterogeneous reactive proximal tubular states within an inflammatory signaling network

We next sought to define the cellular state associated with tubular C3 expression and its relationship to the surrounding microenvironment in FAN. Reclustering of proximal tubular (PT) nuclei resolved differentiated PT-S1, PT-S2 and PT-S3 populations together with the three injury-associated states, termed PT-reactive1, PT-reactive2 and PT-reactive3 (**Figure 2A**). *C3* expression was concentrated predominantly within these reactive populations (**Figures 2A** and **S5A-B**), which showed increased expression of injury-associated genes including *Havcr1*, *Vcam1*, *Cd44*, *Cxcl1*, *Cxcl2*, *Ccl2*, loss of segment-specific differentiated markers (**Figures 2B** and **S5A-B**), and transcriptional features characteristic of previously described maladaptive and failed-repair proximal tubular^24–26^. Although *C3* and *Vcam1*, one of the most widely used markers of maladaptive and failed-repair proximal tubular states^24,27^, were both enriched within reactive PT states, they identified only partially overlapping cells. Among PT-reactive nuclei with detectable *C3* expression, 64% also expressed *Vcam1*, whereas 36% did not; conversely, only 39% of *Vcam1*-positive PT-reactive nuclei showed detectable *C3* (**Figure 2B**). Consistent with this, confocal microscopy identified both VCAM1-positive and VCAM1-negative C3-reporter-positive tubules following injury (**Figure 2C**; vehicle control in **Figure S5C**). These data show that *C3* is expressed in a heterogeneous reactive proximal tubular population that overlaps with, but is not fully captured by, *Vcam1* expression alone.

**Figure 2.**
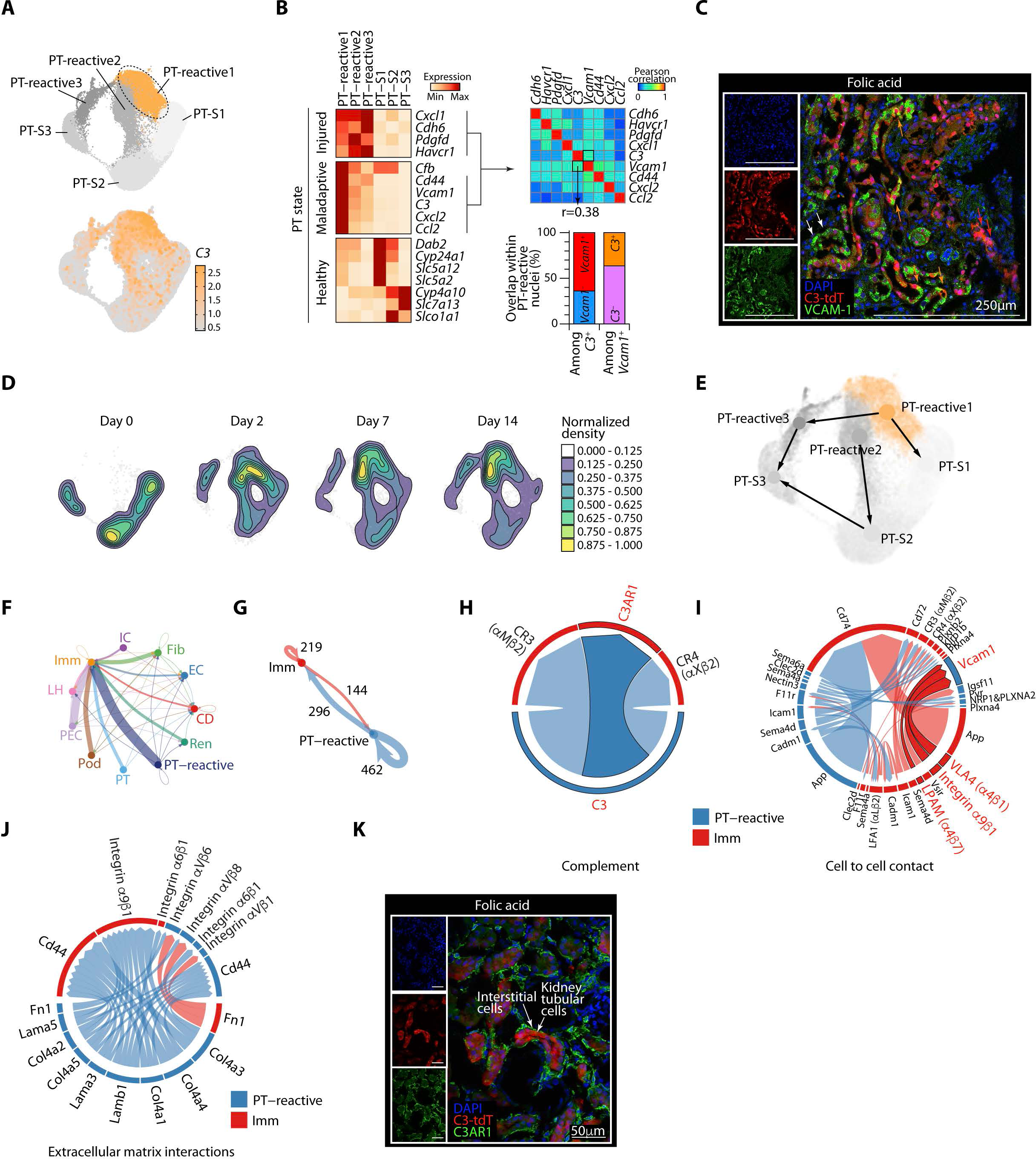
C3 marks heterogeneous reactive proximal tubular states within an inflammatory signaling network. A, UMAP visualization of reclustered proximal tubular populations from FAN kidneys (top) and *C3* expression across these populations (bottom). **B**, Heatmap showing expression of selected markers of injured, maladaptive and differentiated proximal tubular states (left); Pearson correlations among selected reactive-state genes (upper right); and overlap of detectable *C3* and *Vcam1* expression among PT-reactive (lower right). **C**, Representative kidney section following folic acid injury showing *C3*-tdTomato reporter signal and VCAM1 expression. Orange arrows indicate tubules positive for both C3-tdTomato and VCAM1; red and white arrows indicate tubules showing only *C3*-tdTomato^+^VCAM1^−^ and *C3*-tdTomato^−^VCAM1^+^ tubules, respectively. Scale bar, 250μm. Data are representative images from n=3 independent experiments. **D**, Density maps showing temporal redistribution of proximal tubular nuclei within the shared UMAP embedding at days 0, 2, 7 and 14 after folic acid administration. Color indicates normalized density. **E**, Velocity-informed partition-based graph abstraction (PAGA) summarizing inferred directional relationships among differentiated and reactive proximal tubular states across the integrated dataset. Arrows indicate the dominant inferred direction between connected cell states. **F-G**, CellChat analysis of inferred cell-cell communication among the indicated kidney cell populations. Line width denotes inferred interaction number (**F**); and inferred bidirectional communication between PT-reactive and immune-cell populations (**G**). Numbers in **G** indicate the number of inferred ligand-receptor interactions in each direction. Imm, immune cells; IC, intercalated cells; Fib, fibroblasts; EC, endothelial cells; CD, collecting duct cells; Ren, juxtaglomerular renin-producing granular cells; PT, proximal tubular cells; PT-reactive, reactive proximal tubular cells; Pod, podocytes; PEC, parietal epithelial cells; LH, loop of Henle cells. **H**, Chord diagram showing selected inferred complement-mediated interactions between PT-reactive and immune cells, including C3-C3aR1 and interactions involving the β2-integrin complement receptors Mac-1 (αMβ2; CD11b/CD18) and αXβ2 (CD11c/CD18). **I**, Chord diagram showing selected inferred cell-contact interactions between PT-reactive and immune cells. **J**, Chord diagram showing selected inferred extracellular-matrix interactions involving PT-reactive and immune cells. **K**, Representative kidney section after folic acid injury showing *C3*-tdTomato reporter signal and C3aR1**-**positive interstitial cells. Images are representative examples from n=3 independent experiments. Scale bar, 50μm. snRNAseq data in Figures 2 are from n=1 vehicle and n=2 folic acid treated mice per timepoint.

The abundance and organization of these proximal tubular cells changed substantially over time. FAN injury shifted the distribution of proximal tubular cell populations, with expansion of reactive populations and reciprocal contraction of differentiated populations during the acute injury and repair phases (**Figures 2D** and **S5D**). To investigate the relationships between these changes, we inferred RNA velocity from spliced and unspliced transcripts and summarized cluster-level directional relationships using velocity-informed graph abstraction (PAGA)^28,29^. Across the integrated dataset, the dominant inferred trajectories included transitions from each reactive PT state toward differentiated segmental states, with directionality from PT-reactive1 to PT-S1, PT-reactive2 to PT-S2 and PT-reactive3 to PT-S3 (**Figure 2E**). Time-resolved RNA-velocity fields at days 2, 7 and 14 were consistent with dynamic transitions among these states during the injury-repair response (**Figure S5F**). These inferred trajectories suggest that C3-high reactive proximal tubular cells can transition back toward more differentiated transcriptional segmental states during repair.

We next asked how these reactive proximal tubular cells were positioned within the multicellular signaling network of the injured kidney using CellChat^30^. For this analysis, related transcriptional states were grouped into broader cell populations, with PT-reactive comprising the combined PT-R1, PT-R2 and PT-R3 populations. CellChat ligand-receptor inference predicted extensive communication between PT-reactive cells and epithelial, immune, endothelial and stromal populations (**Figure 2F**). Among these interactions, communication with immune cells was particularly prominent and was predicted in both directions (**Figures 2G** and **S6A**). Analysis restricted to complement-mediated interactions identified PT-reactive cells as the predominant source of outgoing complement signals and immune cells as the principal recipient population (**Figure S6B**). Predicted complement interactions linked tubular C3 with C3a-binding C3aR1 and with inactive C3b (iC3b)-binding β2-integrin complement receptors CR3 (CD11b/CD18) and CR4 (CD11c/CD18) on immune cells (**Figure 2H**), consistent with the predominant expression of *C3* in reactive tubules and of *C3ar1*, *Itgam* and *Itgb2* in myeloid cells (**Figures 1J and S3C**). The predicted interaction network extended beyond complement. Reactive proximal tubular and immune cells were connected by multiple cell-contact pathways, including VCAM1-integrin interactions (**Figure 2I**), while reactive proximal tubular cells also participated in extracellular-matrix interactions involving fibronectin, laminins, collagens, CD44 and integrins (**Figure 2J**). Thus, C3-high reactive proximal tubular cells occupy an injury-associated signaling niche characterized by complement, adhesive and matrix interactions with surrounding cell populations.

To determine whether the predicted complement interactions were reflected spatially in injured tissue, we examined C3-expressing tubules and C3aR1-expressing cells *in situ*. Following FAN, *C3*-tdTomato^+^ tubules were frequently juxtaposed to C3aR1-positive interstitial cells (**Figure 2K**), consistent with the predominantly myeloid expression of *C3ar1* observed by single-nucleus transcriptomics. Vehicle-treated kidneys showed little C3 reporter or C3aR1 staining (**Figure S6C**).

Together, these analyses identify C3-high reactive proximal tubular cells as a heterogeneous and dynamically evolving injury-associated population embedded within an extensive multicellular signaling network.

### IL-1β directly induces the tubular C3-FB program through AP-1 and NF-κB

The prominent communication between C3-expressing reactive proximal tubular cells and immune cells suggested that inflammatory signals within the local injury niche might drive the tubular C3-FB response. Direct comparison of reactive and differentiated PT populations revealed enrichment of cytokine-signaling pathways in PT-reactive cells (**Figure S7A**). To identify candidate cytokine signals, we performed Ingenuity Pathway Analysis of genes differentially expressed in proximal tubular cells isolated from folic acid-compared with vehicle-treated mice. This analysis predicted activation of several inflammatory cytokines, with IL-1 among the most strongly activated cytokine pathways (**Figure 3A**). To test whether these candidate signals could directly induce tubular *C3* expression, we exposed tubular cells from *C3*-tdTomato reporter mice to a panel of cytokines selected from the upstream-regulator analysis together with additional inflammatory cytokines implicated in epithelial injury. IL-1β produced a significant increase in C3 reporter fluorescence and was the only inducer among the cytokines tested (**Figures 3B**). Single-nucleus analysis localized *Il1b* expression to myeloid, particularly the monocyte/macrophage, populations within the injured kidney, identifying these cells as likely local sources of IL-1β (**Figure 3C**). Consistent with this, an IL-1R pathway signature was enriched in reactive relative to differentiated proximal tubular cells and increased following injury (**Figures S7B-C**).

**Figure 3.**
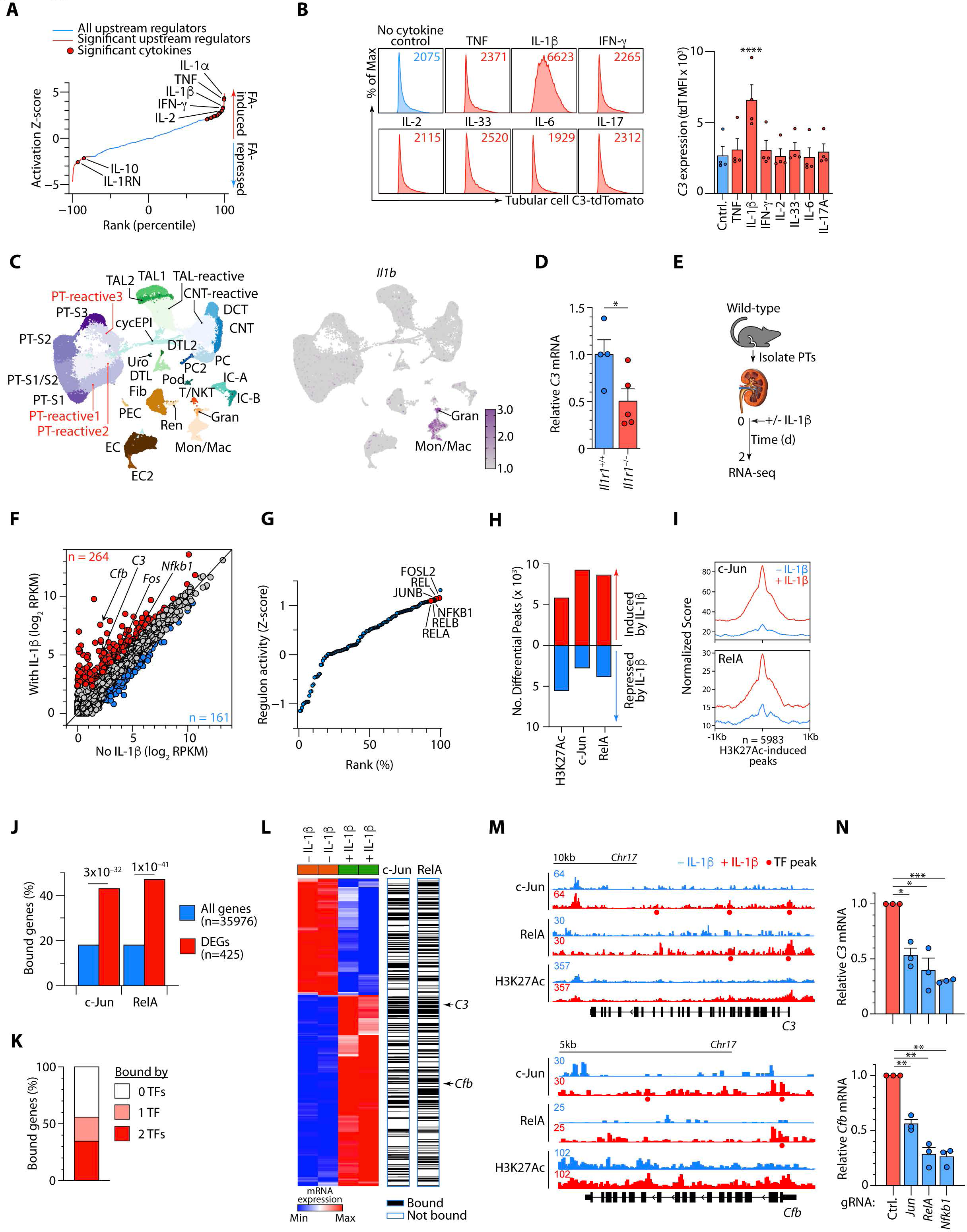
IL-1β induces the tubular C3-FB program through AP-1 and NF-κB. A, Predicted upstream regulators of the proximal tubular transcriptional response to folic acid injury, ranked by activation Z-score. Significant upstream regulators are shown in red; cytokines are indicated by filled circles. **B**, Representative flow-cytometry histograms (left) and quantification (right) from n=2 experiments of *C3*-tdTomato reporter expression in primary renal tubular epithelial cells following treatment with the indicated cytokines. Numbers within histograms indicate median fluorescence intensity (MFI). Dots represent individual mice. **C**, UMAP representation of kidney cell populations (left) and *Il1b* expression across these populations following injury (right). **D**, Relative *C3* mRNA expression in tubular cells isolated from *Il1r1*^+/+^ and *Il1r1*^−/−^ mice following folic acid injury. Each point represents an individual mouse from n=2 independent experiments. **E**, Experimental design for isolation and IL-1β stimulation of primary proximal tubular epithelial cells followed by RNA-sequencing. **F**, Comparison of gene expression in untreated and IL-1β-treated primary proximal tubular epithelial cells (n=2 per group). Genes significantly induced or repressed by IL-1β are shown in red and blue, respectively; selected genes are indicated. **G**, SCENIC-inferred transcription-factor regulon activities ranked across proximal tubular cells. Selected AP-1 and NF-κB family members are indicated. **H**, Numbers of genomic regions showing significantly increased or decreased H3K27Ac, c-JUN or RELA signal following IL-1β stimulation. **I**, Aggregate c-JUN and RelA occupancy centered on the 5,983 genomic regions showing increased H3K27Ac following IL-1β-stimulation. **J**, Proportion of all expressed genes and IL-1β-regulated genes associated with c-JUN or RelA occupancy. **K**, Proportion of IL-1β-regulated genes associated with neither, one or both c-JUN or RELA. **L**, Heatmap showing expression of IL-1β-regulated genes before and after stimulation, together with annotation of c-JUN and RELA occupancy; *C3* and *Cfb* are indicated. **M**, Genome-browser tracks showing c-JUN, RelA and H3K27Ac across the *C3* and *Cfb* loci before and after IL-1β stimulation. Called transcription-factor peaks are indicated by dots. **N**, Relative *C3* (top) and *Cfb* (bottom) mRNA expression 24h after CRISPR-Cas9 targeting of *Jun*, *Rela* or *Nfkb1*, relative to control gRNA, in IL-1β-stimulated tubular epithelial cells. Bars show mean ± s.e.m. *p<0.05; **p<0.01; ***p<0.001; ****p<0.0001. snRNAseq data in Figure 3 are from n=1 vehicle and n=2 folic acid treated mice per timepoint.

We next tested whether IL-1 signaling contributed to tubular *C3* induction *in vivo* in mice lacking IL-1R, the primary receptor for IL-1β. Following FAN, *C3* expression was significantly reduced in tubular cells isolated from *Il1r1*^−/−^ compared with *Il1r1*^+/+^ mice (**Figure 3D**). Consistent with a direct epithelial requirement for IL-1R1 signaling, CRISPR-mediated disruption of *Il1r1* in isolated proximal tubular epithelial cells derived from wild-type mice similarly attenuated IL-1β-induced expression of both *C3* and *Cfb* (**Figure S7D**).

To define candidate transcriptional programs linking IL-1β signaling to complement gene induction, we performed RNA sequencing of proximal tubular cells following IL-1β stimulation (**Figure 3E)**. IL-1β induced 264 and repressed 161 genes, with *C3* and *Cfb* among the induced transcripts (**Figure 3F**). The induced program also included *Fos* and *Nfkb1*, encoding components of the AP-1 and NF-κB transcription-factor families that act downstream of IL-1R1 signaling. In parallel, gene-regulatory network analysis of the kidney snRNA-seq dataset using SCENIC^31^ identified increased AP-1 and NF-κB family regulon activity within reactive proximal tubular populations *in vivo* (**Figures 3G** and **S8A-B**). Together, these observations implicated AP-1 and NF-κB as candidate transcriptional mediators of IL-1β-induced tubular *C3* and *Cfb* expression.

We therefore profiled genome-wide occupancy of JUN and RELA (key components of AP-1 and NF-κB), together with the active chromatin mark histone 3 lysine 27 acetylation (H3K27Ac), by CUT&Tag in tubular epithelial cells before and after IL-1β-stimulation (**Figure S9A**). IL-1β produced widespread changes, with thousands of genomic regions showing increased H3K27Ac or increased c-JUN or RELA occupancy (**Figures 3H** and **S9B-D**). c-JUN and RELA were strongly enriched around the 5,983 regions that gained H3K27Ac following IL-1β treatment (**Figure 3I**), linking recruitment of these transcription factors to IL-1β-induced activation of regulatory chromatin.

We then integrated transcription-factor occupancy with the IL-1β-induced transcriptional response. IL-1β-responsive genes were significantly more likely than genes overall to be associated with c-JUN or RELA binding (**Figure 3J**). More than half of the differentially expressed genes were associated with at least one of these factors, with approximately one-third associated with both c-JUN and RELA (**Figures 3K** and **S9E**). *C3* and *Cfb* among the IL-1β-responsive genes occupied by both c-JUN and RELA (**Figure 3L**). At the individual loci, IL-1β induced c-JUN and RELA occupancy together with increased H3K27Ac across regulatory regions associated with both *C3* and *Cfb* (**Figure 3M**).

Finally, to test whether AP-1 and NF-κB activity was required for *C3* and *Cfb* induction, we disrupted *Jun*, *Rela* or *Nfkb1* in proximal tubular epithelial cells using CRISPR-Cas9 and stimulated the edited cells with IL-1β. Disruption of each factor attenuated IL-1β-induced *C3* and *Cfb* expression at 24h and 48h (**Figures 3N** and **S9F-G**). Together, these findings establish an IL-1β-IL-1R1 signaling axis in which myeloid-derived inflammatory signals engage AP-1 and NF-κB at the *C3* and *Cfb* loci to drive the tubular C3-FB program following kidney injury.

### FB regulates cell-intrinsic C3 and supports metabolic adaptation in proximal tubular cells

We next examined the fate of IL-1β-induced C3 and CFB. Native immunoblotting detected increased intracellular C3 following IL-1β stimulation (**Figure S10A**), while immunoblotting of cell-culture supernatants confirmed increased abundance of both C3 and FB (**Figure S10B**). In a complementary proximal-tubule epithelial model, IL-1β also increased intact C3 together with its cleavage product C3a in cell-culture supernatants, indicating that newly synthesized tubular C3 undergoes proteolytic processing and secretion (**Figure 4A**).

**Figure 4.**
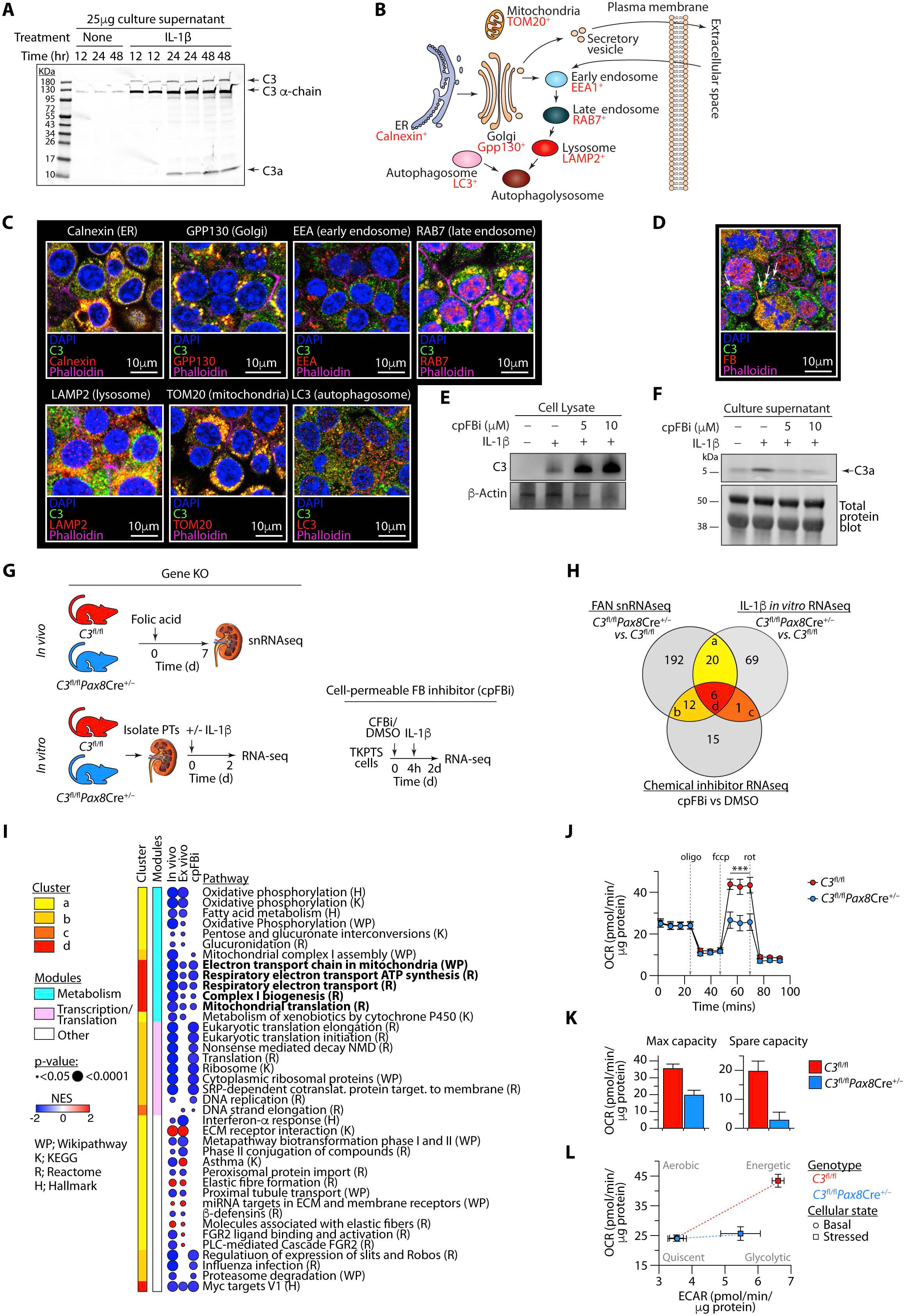
FB regulates intracellular C3 processing and tubular C3 supports metabolic adaptation in proximal tubular cells. **A**, Immunoblot of cell-culture supernatants from TKPTS cells treated with IL-1β for the indicated times, showing intact C3, C3 α-chain and C3a. Shown is a representative example from n=3 experiments. **B**, Schematic illustrating major intracellular protein-trafficking compartments and the canonical secretory pathway. Organelle markers used for subsequent co-localization analyses are indicated in red. **C**, Representative confocal microscopy of IL-1β-treated renal tubular epithelial cells showing C3 together with markers of endoplasmic reticulum (calnexin), Golgi (GPP130), early endosomes (EEA1), late endosomes (RAB7), lysosomes (LAMP2), mitochondria (TOM20) or autophagosomes (LC3). Phalloidin and DAPI mark F-actin and nuclei, respectively. Shown are enlarged regions of the representative images in **Fig. S10C**. Scale bars, 10μm. **D**, Representative confocal image showing intracellular C3 and FB following IL-1β stimulation. White and red arrows indicate C3- and FB-predominant regions, respectively; orange arrows indicate co-localization of the two. Shown are enlarged regions of the representative images in **Fig. S10D**. Scale bar, 10μm. **E**, Representative native immunoblot from n=4 experiments showing intracellular C3 in vehicle- or IL-1β-treated tubular epithelial cells in the presence of the indicated concentrations of cell-permeable FB inhibitor (cpFBi). Because proteins were resolved under native, non-denaturing conditions, standard molecular-weight markers are not applicable and are therefore not shown. β-actin serves as a loading control. **F**, Representative immunoblot from n=4 experiments showing C3a in cell-culture supernatants from tubular epithelial cells treated with IL-1β in the presence of the indicated concentrations of cpFBi. A cropped region of the corresponding total-protein stain is shown below to demonstrate comparable protein loading; the complete total-protein image is shown in **Fig. S10E**. **G**, Experimental designs used to define C3- and FB-dependent transcriptional programs across three complementary perturbations: tubular epithelial *C3* deletion during FAN *in vivo*; *C3* deletion in IL-1β-stimulated primary proximal tubular cells *in vitro*; and pharmacological FB inhibition with cpFBi in IL-1β-stimulated TKPTS cells. **H**, Venn diagram showing significantly enriched pathways identified by fGSEA in FAN kidneys following tubular *C3* deletion, IL-1β-stimulated *C3*-deficient primary proximal tubular cells, and cpFBi-treated TKPTS cells. Letters a-d denote pathway-overlap groups. FAN snRNAseq data are from n=3 WT and n=3 KO samples, IL-1β *in vitro* RNAseq data are from n=2 WT and n=2 KO samples and chemical inhibitor RNAseq data are from n=3 cpFBi and n=2 DMSO samples. **I**, Biological pathways within the four overlap groups (a-d) defined in **H**, grouped by functional module. Dot color indicates normalized enrichment score (NES), with negative and positive values denoting negative and positive enrichment, respectively; dot size indicates statistical significance. Mitochondrial respiratory and translation pathways are highlighted in bold. H, Hallmark; K, KEGG; R, Reactome; WP, WikiPathways. **J**, Oxygen consumption rate (OCR) in IL-1β-stimulated primary renal tubular epithelial cells from *C3*^fl/fl^ and *C3*^fl/fl^*Pax8Cre*^+/−^ mice during sequential addition of oligomycin, FCCP and rotenone. **K**, Maximal respiratory capacity and spare respiratory capacity derived from **J**. **L**, Metabolic phenotype plot showing OCR and extracellular acidification rate (ECAR) under basal and mitochondrial stress-test conditions in IL-1β-stressed control and C3-deficient proximal tubular cells. Data in **J-L** are from n=2 independent experiments. Bars and symbols show mean ± s.e.m. ***p<0.0001.

We therefore asked where newly produced C3 and FB localized within tubular epithelial cells. Confocal microscopy revealed C3 in multiple intracellular compartments, including the endoplasmic reticulum, Golgi apparatus, endosomal and lysosomal compartments, autophagosomes and mitochondria (**Figures 4B-C** and **S10C**). FB also showed broad intracellular localization and substantial overlap with C3, but was additionally present at C3-negative sites, including a prominent nuclear pool (**Figures 4D** and **S10D**). Thus, alongside their release into the extracellular compartment, substantial intracellular pools of C3 and FB were distributed across multiple subcellular compartments.

To determine whether these non-canonical localization patterns were due to tissue-specific alternative transcripts, we examined *C3* and *Cfb* isoform usage in proximal tubular cells. RNA-seq splice-junction and transcript-level analyses showed that tubular cells largely expressed the same canonical *C3* and *Cfb* transcripts observed in hepatocytes, with no major change in isoform usage following IL-1β stimulation (**Figures S11A-E**). These data suggest that the intracellular distribution of C3 and FB may not be due to kidney-specific alternative transcripts.

The extensive intracellular co-localization of FB with C3 raised the possibility that FB might regulate intracellular C3 processing. Consistently, pharmacological inhibition of FB with a cell-permeable small-molecule inhibitor of FB (cpFBi)^14,32^ resulted in dose-dependent accumulation of intracellular C3 protein (**Figure 4E**) without a corresponding increase in *C3* transcription (**Figure S11F; Table 5B**). Consistent with FB-dependent C3 processing, cpFBi also reduced C3a released into cell-culture supernatants (**Figures 4F** and **S10E**). Together, these findings support FB-dependent post-translational processing of tubular-cell-derived C3, with inhibition of cell-intrinsic FB resulting in intracellular C3 accumulation and reduced generation of C3a.

To identify key cellular programs consistently regulated by the C3-CFB pathway, across, we examined transcriptomic responses across three complementary experimental perturbation systems: tubular epithelial C3 deletion following FAN *in vivo*, C3 deletion in primary proximal tubular cells stimulated with IL-1β *in vitro*, and pharmacological FB inhibition in IL-1β-stimulated tubular cells (**Figure 4G**). Tubular C3 deletion was achieved by crossing *C3*^fl/fl^ mice with *Pax8*-*Cre* mice to generate *C3*^fl/fl^*Pax8*^Cre+/–^ mice (**Figure S12A**). For each comparison, genes were ranked by differential expression and analyzed by fast gene set enrichment analysis (fGSEA) to identify pathways altered by loss of C3 or inhibition of FB. Comparison of these pathway-level responses revealed overlapping negatively enriched (depleted) programs across the three perturbations, including a core set shared by all three (**Figure 4H**; **Table 5C**). These convergent pathways were dominated by mitochondrial and metabolic functions, including oxidative phosphorylation, respiratory electron transport, mitochondrial complex I assembly and mitochondrial translation (**Figure 4I**).

This convergence on mitochondrial pathways suggested that the intrinsic C3-FB axis might regulate tubular-cell mitochondrial metabolism. We therefore tested the metabolic function of tubular C3 directly by measuring mitochondrial respiration in primary proximal tubular cells from control and tubular C3-deficient mice using extracellular flux analysis, both at baseline and following IL-1β stimulation. Basal oxygen consumption was similar between genotypes (**Figure S12B**), indicating that C3 was not required for basal mitochondrial respiration. Following IL-1β stimulation, however, C3-deficient cells exhibited an impaired respiratory response during mitochondrial stress testing compared with control cells (**Figure 4J**). This was reflected by reduced maximal respiratory capacity and a particularly pronounced loss of spare respiratory capacity (**Figure 4K**). Extracellular acidification showed comparatively modest differences between genotypes (**Figures S12C-D**). Metabolic phenotype analysis further showed that IL-1β-stimulated control proximal tubular cells transitioned toward a more energetic state, with increases in both oxygen consumption and extracellular acidification, whereas this response was markedly attenuated in C3-deficient cells (**Figure 4L**). Together, these findings indicate that the cell-intrinsic C3-FB axis supports metabolic adaptation of proximal tubular epithelial cells to inflammatory stimulation, with tubular C3 dispensable for basal respiration but required for the full increase in mitochondrial respiratory capacity under metabolic demand.

### Chromatin-bound FB mediates histone proteolysis and promoter remodeling

To determine whether FB expression was necessarily coupled to C3 in renal epithelium, we examined their expression across proximal tubular states. Although *C3* and *Cfb* showed coordinated expression in reactive proximal tubular cells following injury (**Figures S3D and 1K**), our single-nucleus data also showed that *Cfb* was detectable in differentiated proximal tubular populations with little or no *C3* expression (**Figures 1J and S3C**). RNAscope analysis of healthy kidney confirmed this heterogeneity *in situ*, identifying *Cfb*-expressing tubules lacking detectable *C3* (**Figure 5A**), and showing that homeostatic FB expression can occur independently of C3. This was further supported by the presence of a prominent nuclear FB pool that did not overlap with C3 (**Figure 4C**). Together, these observations raised the possibility that FB might have intracellular functions independent of its C3-regulating activity. Consistent with this possibility, FB inhibition also altered additional pathways, including those related to DNA synthesis and SREBF-dependent gene expression (**Table 5C**), distinct from the C3-linked metabolic programs identified above. We therefore investigated the nuclear FB pool more directly.

**Figure 5.**
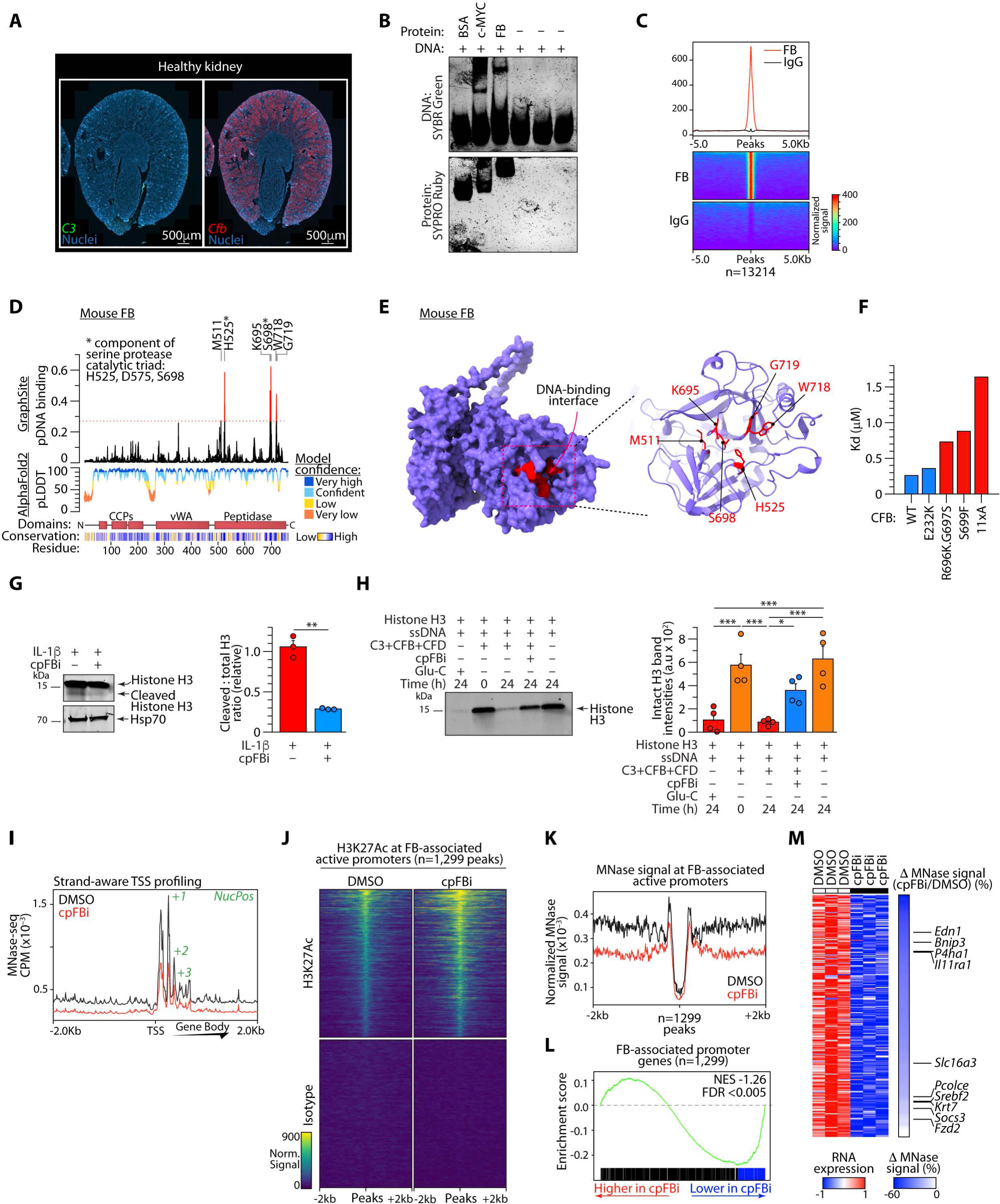
Nuclear FB couples histone proteolysis to promoter chromatin state and transcription. **A**, Representative RNAscope images of healthy mouse kidney showing *C3* (green) and *Cfb* (red) expression. Nuclei are shown in blue. Scale bars, 500μm. **B**, Electrophoretic mobility-shift assay (EMSA) showing binding of recombinant FB to fragmented genomic DNA from renal tubular epithelial cells. BSA and c-MYC serve as negative and positive DNA-binding controls, respectively. DNA and protein are visualized by SYBR Green and SYPRO Ruby staining. Shown is a representative example from n=3 experiments. **C**, Aggregate plot and heatmaps of FB and isotype-control CUT&Tag signal centered on FB-bound genomic regions. **D**, GraphSite-predicted DNA-contacting residues across AlphaFold2-predicted mouse FB sequence, shown together with AlphaFold2 model confidence, protein domains and residue conservation. The dotted line denotes the GraphSite prediction threshold used to classify candidate DNA-contacting residues, which are labeled above the plot. Asterisks denote residues belonging to the serine-protease catalytic triad (as indicated on the left). **E**, AlphaFold2 model of mouse FB showing the predicted C-terminal DNA-binding interface, with candidate DNA-contacting residues highlighted in red (left). An enlarged view of the predicted interface and associated residues is shown at right. **F**, Apparent dissociation constants (*K*_D_) for recombinant wild-type FB and the indicated FB variants measured by quantitative EMSA using a double-stranded oligonucleotide containing a sequence motif enriched within FB-bound genomic regions (**Methods**). *K*_D_ values were estimated from n=2 independent experiments. Please note that GraphSite analysis of FB^11xAlaMut^ additionally identified R175 as a candidate DNA-contacting residue not predicted in wild-type FB (**Figure S14A**), suggesting a potential alternative DNA-contact that could contribute to residual binding following disruption of the principal predicted interface. **G**, Representative immunoblot (left) and cumulative quantification (right) of intact and cleaved histone H3 in IL-1β-treated tubular epithelial cells with or without cpFBi. Hsp70 serves as a loading control. The cleaved-to-total H3 ratio is shown relative to vehicle-treated cells; n=3 independent experiments. **H**, Reconstituted histone-processing assay containing histone H3 and ssDNA together with C3, FB and Factor D, with or without cpFBi. Glu-C serves as a positive proteolysis control, and the 0-h reaction as a baseline control. Representative immunoblot (left) and quantification of intact histone H3 (right) from n=4 independent experiments are shown. **I**, Strand-aware aggregate MNase-seq profile centered on transcription start sites (TSSs) in IL-1β-treated tubular epithelial cells after DMSO or cpFBi treatment, showing the TSS and +1, +2 and +3 nucleosome positions. **J**, Heatmaps of H3K27Ac CUT&RUN signal across 1,299 FB-associated active promoters after DMSO or cpFBi treatment, with corresponding isotype controls. **K**, Aggregate normalized MNase-seq signal across the same 1,299 FB-associated active promoters after DMSO or cpFBi treatment. **L**, Pre-ranked gene-set enrichment analysis of genes linked to the 1,299 FB-associated active promoters within the transcriptomic response to cpFBi versus DMSO. Gene ranks extend from higher expression with cpFBi (left) to lower expression with cpFBi (right). Marked in blue are leading-edge genes. **M**, Heatmap showing RNA expression of leading-edge genes contributing to the enrichment in **L** in DMSO- and cpFBi-treated cells, together with the percentage change in MNase signal at their corresponding FB-associated promoters following cpFBi treatment relative to DMSO. Selected genes associated with extracellular-matrix remodeling, inflammatory signaling, metabolism and cellular stress are annotated. Bars show mean ± s.e.m. where indicated. *p<0.05; **p<0.01.

Confocal microscopy confirmed nuclear localization of FB in IL-1β-stimulated tubular epithelial cells, including in Ki67-negative cells (**Figure S13A; Video S1**). Reanalysis of two independent published mouse cardiac nuclear-proteomics datasets similarly identified FB in nuclear-enriched fractions (**Figures S13B-C**), consistent with presence of FB in the nuclear compartment. To determine whether nuclear FB directly engages chromatin, purified FB was tested for DNA binding by EMSA. FB produced a mobility shift of fragmented genomic DNA from renal tubular epithelial cells, whereas BSA did not, with c-MYC serving as a positive DNA-binding control (**Figure 5B**), suggesting that FB can bind DNA directly. To test whether FB associates with chromatin in intact cells, we performed FB CUT&Tag in tubular epithelial cells. This analysis identified 13,214 FB-enriched genomic regions relative to isotype control (**Figure 5C**), with 55% mapping to promoter or intragenic regions (**Figure S13D**). To quantify FB-DNA binding, we used an oligonucleotide corresponding to an enriched sequence motif identified within FB-bound regions (**Methods**). Quantitative EMSA demonstrated concentration-dependent binding of FB to DNA, with a mean apparent *K*_D_ of 0.26 μM (**Figure S13E**). Together, these findings demonstrate that FB binds DNA directly *in vitro* and occupies chromatin broadly in intact tubular epithelial cells.

To define the structural basis of the FB-DNA interaction, we used GraphSite together with AlphaFold2^33,34^ to predict potential DNA-contacting residues in mouse and human FB (**Figures 5D-E** and **S14A-B**). Six and 11 residues were predicted in mouse and human FB, respectively. In both species, these residues clustered mostly within the C-terminal peptidase domain, including positions within and surrounding the serine-protease catalytic triad (**Figures 5D** and **S14A**). Structural modelling indicated a candidate DNA-binding interface incorporating these residues in both species (**Figures 5E** and **S14B; Video S2**). We tested this interface using recombinant human FB carrying amino acid substitutions at predicted DNA-binding sites. A control substitution outside the predicted interface (E232K) had little effect on DNA binding, whereas substitutions at predicted DNA-interacting residues within the C-terminal interface produced progressively weakened binding, culminating in a >6-fold increase in apparent *K*_D_ for an 11-residue alanine-substitution mutant (**Figures 5F** and **S14C-E**). Structural modeling predicted preservation of the global FB fold in the 11-residue mutant (**Figures S14C-D**). Together, these data identify an unexpected DNA binding domain within FB and support a conserved C-terminal interface that contributes to its activity.

The proximity of the DNA-binding interface to the FB serine-protease catalytic machinery raised the possibility that chromatin-associated proteins might be substrates for nuclear FB-dependent proteolysis. Given the known proteolytic processing of histone H3 and the DNA-binding and protease activities of FB, we focused on histone H3, a core component of the nucleosome, as a potential substrate^35^. In protein extracts from IL-1β-stimulated renal tubular epithelial cells, a lower-molecular-weight H3 species consistent with proteolytic processing was readily detected, whereas treatment with the cpFBi^14,32^ markedly reduced the cleaved-to-total H3 ratio (**Figure 5G**). We next examined H3 proteolytic processing in a reconstituted biochemical system containing histone H3 and ssDNA together with FB, with C3 and Factor D included to permit FB activation. Incubation of these components resulted in loss of intact H3, which was prevented by FB inhibition (**Figure 5H**). We suspect that the more extensive loss of H3 in this reconstituted system likely reflects increased substrate accessibility in the absence of nucleosomal context, allowing more complete proteolysis. By contrast, as shown in **Figure 5G**, nucleosome-bound H3 is primarily cleaved at the exposed N-terminal tail, yielding a shorter proteolytic fragment rather than complete degradation. These data support FB-dependent histone H3 proteolysis and suggest a role for the DNA-binding and protease activities of FB in chromatin regulation.

Because H3 processing increases chromatin accessibility, we next examined the consequences of FB activity for chromatin organization. FB and H3K27Ac CUT&RUN were combined with MNase-seq in IL-1β-stimulated tubular epithelial cells with or without FB inhibition, thereby profiling FB occupancy and active regulatory chromatin together with nucleosome-protected DNA^36^. These analyses revealed a marked bias toward increased H3K27Ac following FB inhibition, with substantially more H3K27Ac-positive regions gaining than losing signal (**Figure S15A**). FB occupancy was enriched at transcription start site (TSS)-associated H3K27Ac-positive regions relative to distal active regulatory regions (**Figure S15B**), directing our subsequent analyses toward promoter chromatin. MNase-seq analysis showed the expected phased nucleosome organization around TSSs. FB inhibition reduced normalized MNase signal across the TSS and +1 to +3 nucleosome positions while largely preserving nucleosome periodicity (**Figures 5I and S15C-D**), indicating reduced nucleosome-associated DNA protection.

Intersection of FB occupancy, H3K27Ac enrichment and TSS association identified 1,299 FB-associated active promoters (**Figure 5J**). Across these regions, FB inhibition resulted in a broad increase in H3K27Ac (**Figures 5J**) together with reduced normalized MNase signal at 98% of these promoters (1,271 of 1,299) (**Figure 5K**). The *Pax8* locus, encoding a canonical renal epithelial lineage transcription factor, provides a representative example (**Figure S15E**). Together with the biochemical evidence for FB-dependent H3 processing, these findings suggest that promoter-bound FB promotes local H3 processing and altered promoter chromatin organization.

To connect these chromatin changes with transcriptional output, genes linked to the 1,299 FB-associated active promoters were tested against the transcriptomic response to cpFBi using pre-ranked gene-set enrichment analysis^37^. FB-associated promoter genes were significantly enriched among genes repressed following FB inhibition (**Figure 5L**). Leading-edge genes included regulators of extracellular-matrix remodeling, inflammatory signaling, metabolism and cellular stress, including *Edn1*, *Bnip3*, *P4ha1*, *Il11ra1*, *Slc16a3*, *Pcolce*, *Srebf2*, *Krt7*, *Socs3* and *Fzd2* (**Figure 5M**), several of which have been associated with kidney injury, metabolic reprogramming and fibrotic remodeling^38–41^. Their decreased transcriptional output was accompanied by reduced MNase signal at the corresponding FB-associated promoters (**Figure 5K**). Thus, FB inhibition produced a chromatin state associated with reduced MNase-protected nucleosome occupancy at FB-bound promoters and preferential repression of their linked genes.

Promoters most sensitive to FB inhibition were also examined for shared regulatory features that might identify the transcriptional contexts for the nuclear FB function. Among FB-associated active promoters showing ≥2-fold increased H3K27Ac after FB inhibition, *de novo* motif analysis against the remaining FB-associated active promoters as background identified KLF-like and TEAD-like motifs as the two most enriched sequence features (**Figure S15F**), with AP-1/bZIP-family motifs also overrepresented (**Figure S15G**). Genes linked to the 500 most responsive promoters were enriched for proliferative, metabolic, hypoxic and inflammatory programs (**Figure S15H**). Notably, these and TEAD/YAP, AP-1/JNK and KLF6 regulatory pathways have independently been implicated in proximal tubular injury, metabolic reprogramming and fibrotic repair^42–44^. Together, these findings support a model in which nuclear FB occupies selected promoters and couples FB-dependent histone H3 processing to promoter chromatin organization and transcriptional programs relevant to maladaptive proximal-tubule repair.

### Tubular C3 promotes maladaptive repair and kidney fibrosis with minimal effect on acute injury severity

To determine whether the tubular complement response contributes causally to maladaptive repair *in vivo*, we initially sought to perturb FB. However, the available *Cfb*-deficient strain was not suitable for testing cell-intrinsic FB function because this mouse retains intracellular FB despite loss of circulating protein^45^, analogous to our previous finding that conventionally *C3*-deficient mice lacking serum C3 retain intracellular C3^46^. The cell-permeable FB inhibitor used above was also unsuitable for *in vivo* animal studies because of suspected liver toxicity. CRISPR-Cas9 targeting of *Cfb* in primary tubular epithelial cells markedly reduced viable-cell recovery, precluding a complementary genetic loss-of-function approach (**Figure S16A**). We therefore turned to tubular C3, which could be efficiently and selectively deleted from renal tubular epithelium in *C3*^fl/fl^*Pax8*^Cre+/–^ mice and for which our preceding experiments had established a functional link to mitochondrial and metabolic changes noted during kidney injury, including altered metabolic programs *in vivo* and impaired respiratory reserve in C3-deficient proximal tubular cells (**Figures 4F-K** and **6A**). We used this mouse model to ask whether loss of tubular *C3* altered the transition from acute injury to maladaptive repair *in vivo*.

Folic acid induced a comparable increase in blood urea nitrogen in *C3*^fl/fl^ and *C3*^fl/fl^*Pax8*^Cre+/–^ mice at day 2 (**Figure 6B**), indicating that tubular *C3* deletion did not substantially alter the onset or severity of the initial acute kidney injury. To examine the intervening tissue response at day 7, we performed kidney snRNA-seq together with CD45^+^ cell-enriched scRNA-seq and flow cytometry (**Figure 6A**). Major renal and immune-cell populations were represented in both genotypes, with macrophages comprising the largest immune-cell population after injury (**Figures S16B-E**). Tubular *C3* deletion was associated with reduced macrophage representation, which was independently confirmed by flow cytometry (**Figures 6C**). This was consistent with enrichment of *C3ar1* in monocyte/macrophage populations associated with C3-expressing reactive tubules (**Figure 1J**), as well as with established C3a-C3aR-dependent macrophage chemotaxis and reduced renal macrophage accumulation in *C3ar1*-deficient inflammatory kidney disease^47–50^. Together, these findings support a model in which tubular C3 promotes macrophage accumulation after injury.

**Figure 6.**
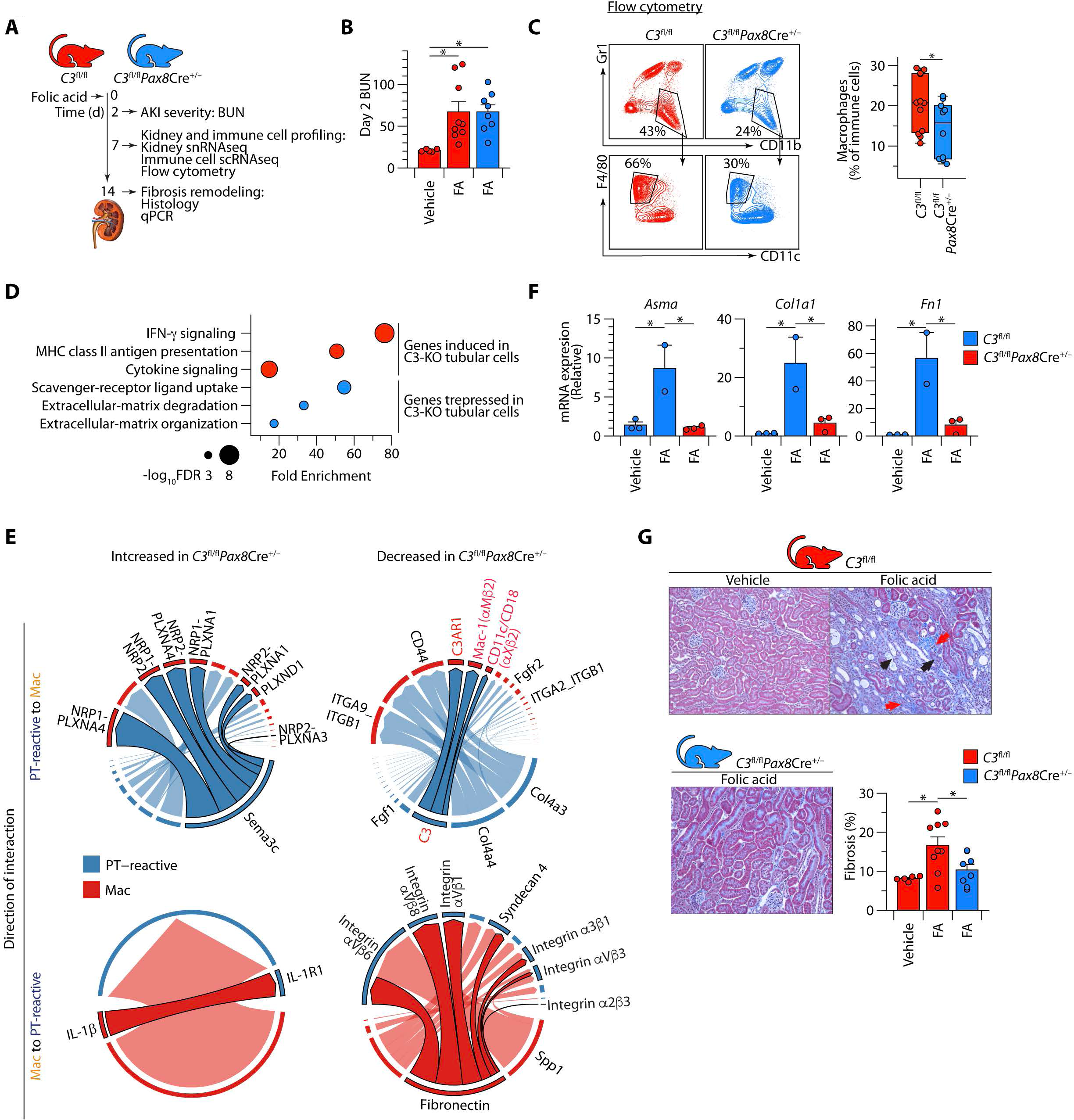
Tubular C3 promotes maladaptive repair and kidney fibrosis with minimal effect on acute injury severity. **A**, Experimental design for folic acid nephropathy in *C3*^fl/fl^ and *C3*^fl/fl^*Pax8Cre*^+/−^ mice. Acute kidney injury severity was assessed by blood urea nitrogen (BUN) at day 2; kidney and immune-cell responses by single-cell profiling and flow cytometry at day 7; and fibrotic remodeling by histology and qPCR at day 14. **B**, BUN concentrations at day 2 after vehicle or folic acid administration in the indicated genotypes. Each point represents an individual mouse. **C**, Representative flow-cytometry plots from folic-acid-injured kidneys of the indicated genotypes (left) and quantification of macrophages as a proportion of renal immune cells (right). Each point represents an individual mouse. Data are from n=2 experiments. **D**, Selected Reactome pathways over-represented among genes expressed at higher levels in macrophages from *C3*^fl/fl^*Pax8Cre*^+/−^ kidneys (red) or *C3*^fl/fl^ kidneys (blue). Bubble position indicates fold enrichment and bubble size indicates −log10 FDR. Data are from n = 2 mice per genotype. **E**, Chord diagrams showing selected ligand-receptor interactions between PT-reactive cells and macrophages that were increased (left) or decreased (right) following tubular *C3* deletion. Arc color denotes the signaling cell population and ribbon direction indicates the predicted direction of communication. Selected ligand-receptor pairs are indicated. **F**, Renal expression of the fibrosis-associated genes *Acta2* (encoding αSMA), *Col1a1* and *Fn1* at day 14. Each point represents an individual mouse. **G**, Representative Masson’s trichrome-stained stained kidney sections from vehicle-treated *C3*^fl/fl^ mice and folic acid-treated *C3*^fl/fl^ and *C3*^fl/fl^*Pax8Cre*^+/−^ mice (left), with quantification of interstitial fibrosis (right). Arrowheads indicate representative areas of fibrosis (red) and tubular injury (black). Each point represents an individual mouse. Data in **B**, **G** and **F** are from n=3 independent experiments. Bars show mean ± s.e.m. *p<0.05.

Given the established role of macrophages in extracellular-matrix remodeling during the transition from acute kidney injury to fibrosis^51^, we also examined whether tubular C3 deletion altered the transcriptional state of the remaining macrophages. Extracellular-matrix and scavenger-receptor programs were reduced, whereas interferon, cytokine-signaling and MHC class II antigen-presentation programs were enriched (**Figure 6D**). This included reduced expression of several remodeling-associated genes, including *Spp1, Fn1, Arg1, Chil3* and *Thbs1* (**Figure S16F; Table S9**). Thus, tubular C3 deletion shifted macrophages away from a remodeling-associated toward a more inflammatory, antigen-presenting state, consistent with prior reports linking C3a-C3aR signaling to repair-associated macrophage polarization^52,53^.

Inferred cell-cell communication analysis further suggested selective remodeling of signaling between PT-reactive cells and macrophages. Despite broadly similar total interaction number and mean interaction strength (**Figure S16G**), differential signaling analysis identified changes in extracellular-matrix and adhesion pathways, including collagen-, laminin-, fibronectin-and integrin-associated signaling (**Figure S16H**). Selected PT-reactive-macrophage ligand-receptor interactions altered by tubular C3 deletion are shown in **Figure 6E**. These analyses suggested that loss of tubular C3 was associated not only with fewer macrophages and altered macrophage state, but also with selective remodeling of the predicted matrix-and adhesion-related interactions linking macrophages to reactive tubules within the injury-repair niche.

Finally, fibrosis was attenuated in mice lacking tubular C3. At day 14, expression of the fibrosis-associated genes *Acta2*, *Col1a1* and *Fn1* was significantly reduced in *C3*^fl/fl^*Pax8*^Cre+/–^ kidneys (**Figure 6F**), accompanied by reduced interstitial fibrosis histologically (**Figure 6G**). Together, these findings suggest that tubular C3 was dispensable for the severity of the initial injury but shapes the intervening immune response and promotes subsequent fibrotic remodeling.

### The C3-FB reactive tubular program is conserved in human kidney disease and associates with kidney dysfunction

To determine whether the injury-associated tubular complement program identified in mice was conserved in humans, we analyzed single-cell transcriptomes from the Kidney Precision Medicine Project (KPMP) (**Figure 7A; Figure S17A**). Transcriptome-wide comparison across 47 subjects using one-to-one orthologous genes showed that mouse PT-reactive populations most closely corresponded to injured/dedifferentiated human proximal tubular states, whereas differentiated mouse proximal tubular populations mapped to their corresponding differentiated human states (**Figure 7B**). Consistent with this cross-species mapping, human injured/dedifferentiated proximal tubular cells showed reduced expression of differentiated tubular markers and increased expression of injury-associated genes, including *HAVCR1*, *VCAM1*, *CD44*, *CCL2* and *C3*, together with prominent *CFB* expression (**Figure 7C**).

**Figure 7.**
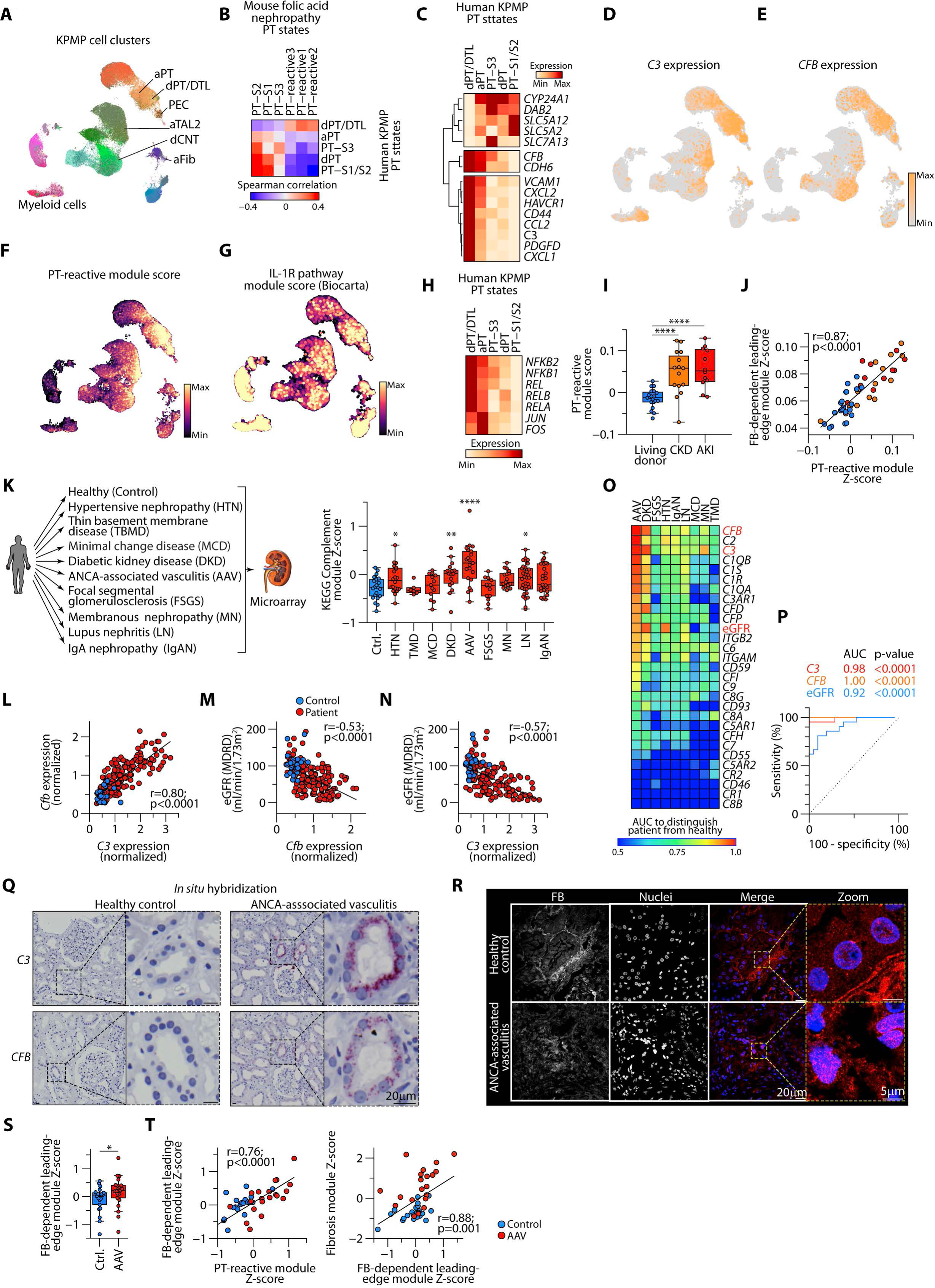
The mouse tubular C3-FB program is conserved in human kidney disease and associates with kidney dysfunction. **A**, UMAP visualization of kidney cell populations in the Kidney Precision Medicine Project (KPMP) single-cell dataset (n=110,346 cells), highlighting proximal tubular states and myeloid populations. aPT, adaptive/maladaptive proximal tubular epithelial cell; dPT/DTL, degenerative proximal tubular epithelial cell/descending thin limb cell; PEC, parietal epithelial cell; aTAL2, adaptive/maladaptive thick ascending limb cell cluster 2; dCNT, degenerative connecting tubule cell; aFib, adaptive/maladaptive fibroblast. **B**, Spearman correlations between gene-expression profiles of mouse proximal tubular populations from folic acid nephropathy and human KPMP proximal tubular states. **C**, Heatmap showing expression of selected homeostatic and injury-associated genes across human proximal tubular states. **D-E**, UMAP feature plots showing *C3* (**D**) and *CFB* (**e**) expression in the human KPMP dataset. **F**, PT-reactive module score across kidney cells. **G**, BIOCARTA IL-1R pathway module score across kidney cells. **H**, Expression of AP-1 and NF-κB family transcription factors across human proximal tubular states. **I**, PT-reactive module scores in proximal tubular cells from healthy (living kidney transplant donors) and patients with chronic kidney disease (CKD) or acute kidney injury (AKI). **J**, Relationship between the FB-dependent leading-edge module score and PT-reactive module score in proximal tubular cells across KPMP samples. Points indicate individual samples from living donors, CKD and AKI. Spearman correlation coefficient and p-value are shown. **K**, Schematic of bulk renal transcriptomic datasets from healthy controls and patients with hypertensive nephropathy (HTN), thin basement membrane disease (TBMD), minimal change disease (MCD), diabetic kidney disease (DKD), ANCA-associated vasculitis (AAV), focal segmental glomerulosclerosis (FSGS), membranous nephropathy (MN), lupus nephritis (LN) or IgA nephropathy (IgAN), with bar plot showing KEGG complement pathway Z-scores in each disease. **L**, Relationship between renal *C3* and *CFB* expression across control and disease samples. **M-N**, Relationship between estimated glomerular filtration rate (eGFR) and renal *CFB* (**M**) or *C3* (**N**) expression. Control and patient samples are indicated. Pearson correlation coefficients and p-values are shown. **O**, Area under the receiver-operating-characteristic curve (AUC) for individual complement genes distinguishing each renal disease from healthy control tissue. *C3*, *CFB* and eGFR (estimated glomerular filtration rate) are highlighted. **P**, Receiver-operating-characteristic curves for *C3*, *CFB* and eGFR distinguishing ANCA-associated vasculitis (n=21) from healthy control samples (n=21). AUC and p-values are shown. **Q**, Representative RNAscope images showing *C3* and *CFB* transcripts in renal tissue from a healthy control and a patient with ANCA-associated vasculitis. Boxed regions are shown enlarged at right. Scale bar, 20μm. Shown are representative images from n=5 patients and n=3 controls. **R**, Representative 40× confocal images of healthy-control and ANCA-associated vasculitis kidney biopsies showing FB staining (grayscale), DAPI nuclear staining (grayscale), and merged images with FB shown in red and nuclei in blue. Enlarged views show selected 25 × 25μm regions from the corresponding merged images. Overview images represent 200 × 200μm fields. Scale bars, 20μm in merged overview images and 5μm in zoomed images. Shown is one representative example from n=3 independent samples per group. **S**, FB-dependent leading-edge module scores in healthy-control and AAV kidney samples. Each point represents an individual sample. **T**, Relationship between the FB-dependent leading-edge module score and the PT-reactive module score (left) or fibrosis-associated module score (right) in healthy-control and AAV samples. Spearman correlation coefficients and p-values are shown.

Human reactive tubular states also recapitulated key features of the regulatory architecture underlying the murine C3-FB response. Both *C3* and *CFB* were expressed within injury-associated proximal tubular populations (**Figures 7D-E**), which also showed high PT-reactive and IL-1R pathway module scores (**Figures 7F-G**). Components of the NF-κB and AP-1 transcriptional machinery implicated in IL-1β-dependent C3-FB induction in mice, including *NFKB1*, *NFKB2*, *REL*, *RELA*, *RELB*, *JUN* and *FOS*, were similarly enriched in injured/dedifferentiated proximal tubular populations (**Figure 7H**). Disease-stratified analysis showed higher tubular *C3* and *CFB* expression in chronic kidney disease (CKD) and acute kidney injury (AKI), alongside the injury marker *VCAM1* (**Figure S17C**). Importantly, the mouse-derived PT-reactive module score was increased in both CKD and AKI relative to healthy living-donor kidneys, with the highest scores in AKI (**Figure 7I**), consistent with the acute injury-associated state captured by the FAN model.

To connect these conserved reactive states to the nuclear FB program defined mechanistically above, we examined the leading-edge genes responsible for enrichment of FB-associated promoter genes among transcripts repressed by FB inhibition (**Figure 5M**). The FB-dependent leading-edge module derived from these genes correlated significantly with the PT-reactive module across KPMP samples (r = 0.79, p < 0.0001; **Figure 7J**). These analyses suggested that the FB-dependent transcriptional program defined experimentally was significantly associated with the human reactive proximal tubular state.

We then extended the analysis across a broader spectrum of human kidney diseases using biopsy transcriptomic data from the European Renal cDNA Bank^54,55^, spanning nine diagnoses and healthy controls (**Figure 7K**, left). Complement-pathway expression varied substantially among diseases, with highest enrichment in ANCA-associated vasculitis (AAV), a severe inflammatory kidney disease frequently presenting with AKI (**Figure 7K**, right). Broad profiling of complement components and receptors similarly demonstrated disease-specific patterns of renal complement expression (**Figure S17D**). Across the cohort, *C3* and *CFB* expression were strongly correlated (r = 0.80, p < 0.0001; **Figure 7L**), consistent with coordinated regulation of these components across human kidney disease.

Renal expression of both genes was also associated with kidney dysfunction. *CFB* expression correlated inversely with estimated glomerular filtration rate (eGFR; r = −0.53, p < 0.0001; **Figure 7M**), as did *C3* (r = −0.57, p < 0.0001; **Figure 7N**). Importantly, these associations remained significant after adjustment for disease type in multivariable linear models (*C3*: β = −25.68, p = 1.16 × 10^−10^; *CFB*: β = −36.87, p = 4.58 × 10^−8^), indicating that the relationship between renal C3-FB expression and kidney dysfunction was not solely attributable to differences among disease categories.

We next asked whether *C3* and *CFB* were associated with disease status. To this end, we performed receiver-operating-characteristic (ROC) analysis across all complement genes and disease groups to assess their ability to distinguish diseased from healthy-control kidneys. Discriminatory performance varied across genes and diagnoses, with *C3* and *CFB* showing particularly strong discrimination in AAV (**Figure 7O**). Specifically, in AAV, the area under the ROC curve (AUC) was 0.98 for *C3*, 1.00 for *CFB* and 0.92 for eGFR (**Figure 7P**). *CFB* discrimination was significantly greater than that by eGFR (p<0.05 by DeLong comparison), whereas the difference between *CFB* and *C3* was not significant; the difference between C3 and eGFR also did not reach significance. Across kidney diseases collectively, *C3*, *CFB* and eGFR showed similar discrimination between diseased and healthy-control kidneys (AUCs 0.86, 0.85 and 0.86, respectively; **Figure S17E**).

Finally, given the particularly strong C3 and CFB disease-associated signal in AAV, we used AAV as a human disease context to seek orthogonal evidence for local tubular C3 and FB expression and FB subcellular localization. RNA *in situ* hybridization showed little tubular *C3* and only a little *CFB* signal in healthy control kidneys, whereas both transcripts were readily detected in renal tubules in AAV (**Figure 7Q**). Notably, *C3* and *CFB* signals overlapped within the same tubular epithelial cells, supporting co-expression of both components of the pathway *in situ*. Immunofluorescence similarly demonstrated abundant tubular FB in AAV kidneys, including a distinct intranuclear pool (**Figure 7R**). Moreover, the FB-dependent leading-edge module score was significantly higher in AAV than in healthy controls (**Figure 7S**) and correlated strongly with the PT-reactive module within AAV (*r* = 0.82, p < 0.0001; **Figure 7T**). FB-dependent leading-edge scores also correlated with a fibrosis-associated transcriptional module (*r* = 0.88, p = 0.001; **Figure 7T**).

Collectively, these findings indicate conservation of the injury-associated C3-FB tubular response across species, link renal C3 and FB expression to kidney dysfunction, and suggest that the experimentally defined FB-dependent transcriptional program tracks with reactive tubular and fibrotic states in human disease. The detection of nuclear FB in diseased human tubules further places the non-canonical FB functions identified mechanistically in experimental systems within a clinically relevant human tissue context.

## Discussion

Here, we identify an injury-induced, cell-intrinsic C3-FB complement program that links inflammatory sensing in proximal tubular epithelial cells to metabolic adaptation, chromatin regulation and tissue repair. IL-1βR signaling recruits NF-κB and AP-1 to *C3* and *Cfb* regulatory elements, inducing local expression of both proteins. Once induced, C3 and FB acquire distinct functions across subcellular compartments that differ from their canonical extracellular activities. FB promotes processing of intracellularly synthesized C3, which, in part, localizes to mitochondria and supports mitochondrial respiratory adaptation, whereas FB additionally enters the nucleus, binds chromatin through a previously unrecognized DNA-binding interface and supports histone H3 processing and promoter chromatin remodeling. *In vivo*, tubular C3 has little effect on acute injury severity but promotes macrophage accumulation and subsequent fibrosis. This response and its upstream regulatory circuitry are conserved in injury-associated proximal tubular states in human kidney disease. Together, these findings extend intracellular complement beyond cell-autonomous activation and show that canonical complement proteins can be repurposed within subcellular compartments as regulators of tissue-cell physiology.

The kidney provides a particularly relevant setting for these findings. AKI can drive incomplete tubular recovery, persistent inflammation and interstitial fibrosis, increasing the risk of CKD and kidney failure^56–59^. Complement deposition occurs across glomerular and tubulointerstitial kidney disorders^23,60–62^ and kidney cells have been known to synthesize complement locally since the 1980s^23,63,64^. Yet how locally produced complement is integrated into injury-associated renal cell states, functions within cells, and influences the transition from acute injury to maladaptive fibrotic repair remain unresolved. This is because bulk tissue measurements cannot readily distinguish complement synthesized by defined renal cell populations from circulating or deposited proteins, while conventional systemic genetic models cannot separate extracellular complement activity from cell-intrinsic functions within kidney cells. By combining cell-resolved transcriptomics, spatial localization, conditional genetics and subcellular functional analyses, our study distinguishes these compartments and identifies locally produced tubular complement as an active component of the injury-response program.

Our findings add to growing evidence that non-canonical, cell-intrinsic functions of complement proteins represent a recurring feature of cell biology rather than isolated exceptions. Studies of intracellular complement have identified roles in metabolism, autophagy, survival and differentiation, with much of the mechanistic framework initially defined in immune cells, particularly lymphocytes^8,16,23^. More recent work suggests that this biology is not restricted to immune cells, but also operates in structural and parenchymal cells. Intracellular FH protects neurons during inflammatory stress independently of secretion and C3^9^; cytosolic C3 regulates autophagy in pancreatic β-cells through ATG16L1; while intracellular CD59 participates directly in the exocytotic machinery controlling insulin secretion^65,66^; and intracellular FH regulates NF-κB signaling and cellular behavior in renal endothelial and malignant kidney cells^67,68^. Our findings identify a further organizing principle of intracellular complement biology: within the same injured epithelial cell, locally produced complement proteins can be partitioned across subcellular compartments to acquire specialized functions in metabolism and chromatin regulation.

This concept may have deep evolutionary roots. C3 and FB are among the most ancient of complement components, with homologs in early metazoans, and predating the classical and lectin pathways^69,70^. Their repurposing for fundamental aspects of cell physiology may therefore reflect functions that preceded their incorporation into the extracellular proteolytic cascade. This possibility is particularly relevant to metabolism. Intracellular complement regulates metabolic programs in other settings: in human CD4^+^ T cells, intracellular C3 activation supports mTOR activity, nutrient uptake, glycolysis and oxidative phosphorylation during effector differentiation^11,46^, whereas mitochondrial C3a negatively regulates respiratory metabolism in stressed retinal epithelial cells^71^. Proximal renal tubular cells have high energetic requirements and depend heavily on mitochondrial oxidative phosphorylation and fatty acid oxidation^72,73^. Our finding that injury-induced tubular C3 localizes in part to mitochondria and is required for full respiratory reserve therefore places intracellular complement within a physiological program central to proximal tubular function and repair. C3-dependent metabolic adaptation may help injured epithelial cells withstand inflammatory stress, whereas persistence of the same program could sustain incompletely repaired states. This provides one explanation for the apparent paradox that intracellular C3 supports metabolic adaptation at the cellular level yet promotes fibrosis at the tissue level. It also suggests that complement may be particularly important in determining the fate of surviving epithelial cells after injury^12,13^.

The nuclear activity of FB extends its function into another fundamental cellular process: chromatin regulation. Nearly five decades ago, two major DNA-binding proteins purified from human serum were identified as FB and β1H, now known as Factor H^74–76^. No clear cellular function or intracellular counterpart was identified, and the observations received little subsequent attention. Our findings provide such a context: nuclear FB occupies chromatin genome-wide, and structural modeling and mutational analysis support a DNA-binding surface within its C-terminal peptidase domain. That two ancient alternative-pathway proteins bind nucleic acids suggests an ancestral and overlooked property of complement proteins. Other non-canonical complement activities are consistent with this broader concept. Microglia-derived C1q can be internalized by neurons and incorporated into RNA-dependent ribonucleoprotein and polysome complexes, altering protein synthesis and proteostasis^77^. Intracellular FH regulates NF-κB nuclear translocation in kidney endothelial and renal carcinoma cells^67,68^; internalized C3 reaches the nucleus of human B cells, where C3 and C3a bind DNA and C3/C3b interact with histones^78^; and tumor-derived C3b acts as a nuclear SIN3A-HDAC1/2 cofactor in chemotherapy-resistant lung cancer, promoting histone deacetylation and transcriptional repression^79^. These observations indicate several routes through which complement can influence transcription and chromatin, but the FB mechanism identified here is distinct: an intrinsic DNA-binding interface linked to histone H3 processing and promoter chromatin remodeling.

Histone proteolysis provides a potential mechanism linking this FB activity to transcriptional regulation. Relatively few histone proteases have been characterized: cathepsin L processes H3 during embryonic stem-cell differentiation, whereas promoter-restricted H3 clipping in yeast precedes histone eviction and is required for normal transcriptional induction^35,80^. Unlike reversible histone modifications, proteolysis removes histone sequences together with their associated regulatory marks and can therefore impose substantial changes in chromatin state. The proximity of the FB DNA-binding interface to its catalytic machinery raises the possibility that DNA binding positions FB-dependent proteolytic activity at chromatin. This could allow chromatin-associated FB to couple DNA binding to local histone processing, encoding transient inflammatory signals into more persistent epithelial-cell states. The marked loss of viability after *Cfb* silencing in primary tubular cells further raises the possibility that stressed tubular cells depend on one or more intracellular FB functions.

Important aspects of intracellular complement biology in kidney tubular cells nevertheless remain unresolved and require dedicated biochemical and spatial investigation. In particular, the precise subcellular sites at which newly synthesized C3 is processed, the nature and relative abundance of the resulting C3-derived fragments, and the compartments in which those fragments signal remain incompletely defined. Our data support FB-dependent post-transcriptional C3 processing, yet *Cfd*, encoding the canonical FB-activating protease Factor D^81,82^, was not detectably expressed in our tubular epithelial datasets. We encountered the same paradox in respiratory epithelial cells, where intracellular C3 activation was FB-dependent despite an unresolved mechanism of FB activation^14^. Defining the biochemical machinery responsible for intracellular FB and C3 processing and mapping the resulting fragments and their activities to specific subcellular compartments, will therefore require further mechanistic investigation.

Recognition of these cell-intrinsic and subcellular functions has important implications for complement therapeutics. C3 and FB are validated renal therapeutic targets: the FB inhibitor iptacopan is approved for C3 glomerulopathy and IgA nephropathy, and the C3/C3b inhibitor pegcetacoplan for C3 glomerulopathy and primary immune-complex membranoproliferative glomerulonephritis^19,20,83^. These successes validate proximal complement inhibition in kidney disease, but existing agents have been developed and pharmacologically characterized principally against circulating or extracellular complement. Whether they achieve meaningful exposure or target engagement within cytoplasmic, mitochondrial or nuclear complement pools is unknown. Effective targeting of complement may therefore need to account not only for the relevant pathway component, but also for its cellular source and subcellular compartment. Intracellular FB may represent one such distinct therapeutic compartment, raising the possibility of approaches that disrupt its DNA-binding interface or intracellular protease activity without necessarily reproducing conventional extracellular pathway inhibition.

Direct pharmacological access to intracellular, and particularly nuclear, complement pools may, however, be challenging. An alternative strategy is to target the signals that induce cell-intrinsic complement in the first place. Our study identifies IL-1β as such a signal in injured proximal tubular cells: macrophage-derived IL-1β engages tubular IL-1R1 and recruits NF-κB and AP-1 to *C3* and *Cfb* regulatory elements. This finding fits a broader principle emerging across cell types in which cell-intrinsic complement is regulated by the environmental signals characteristic of a particular tissue or inflammatory context. In immune cells, we identified LFA-1 engagement during transendothelial migration and cellular interactions to induce intrinsic *C3* transcription^10^. In respiratory epithelial cells infected with SARS-CoV-2, we found that complement induction is instead driven predominantly by type I interferon-JAK1/2-STAT1 signaling, together with NF-κB activity^14^. In the injured kidney, our findings identify IL-1β−IL-1R1-NF-κB/AP-1 signaling as the corresponding regulatory axis. Thus, although intracellular complement functions may be broadly conserved, the signals that induce them can be cell-and context-specific. Identifying these upstream regulatory pathways may provide a therapeutically tractable route to modulating intracellular complement without requiring direct access to every disease-relevant subcellular pool.

This possibility is particularly relevant because IL-1β signaling is already pharmacologically accessible. IL-1R blockade ameliorates experimental renal ischemia-reperfusion injury, and anakinra reduces inflammatory signaling during porcine kidney normothermic machine perfusion^84,85^. Whether IL-1R1 blockade during an appropriate post-injury window can prevent maladaptive repair remains untested, but tubular *C3* deletion attenuated subsequent fibrosis while leaving acute injury severity largely unchanged. In the injured kidney, this regulatory axis also forms part of an epithelial-myeloid feedback circuit: macrophage-derived IL-1β induces tubular *C3* and *Cfb*, whereas tubular C3 feeds back on the myeloid compartment to influence macrophage accumulation and state. C3a-C3aR signaling is one plausible route, given *C3ar1* expression in monocyte/macrophage populations and evidence that C3aR promotes renal macrophage recruitment^47,51^. iC3b signaling through CR3, and potentially CR4^86^, provides an alternative, supported by evidence that locally produced stromal C3 can regulate myeloid infiltration through iC3b-CR3 in tumors^87^. Spatial and single-cell studies likewise implicate monocyte-derived, extracellular-matrix-remodeling macrophages in AKI-to-CKD progression^47,51^. Consistent with such reciprocal interactions, tubular C3 deletion reduced macrophage accumulation and altered macrophage transcriptional programs and macrophage-tubular communication, including extracellular-matrix-associated signaling. Locally produced complement may therefore shape repair both cell intrinsically, through epithelial metabolism and chromatin, and cell extrinsically, through the surrounding immune niche.

The conservation of coordinated C3 and FB expression in human injury-associated proximal tubular states, together with their association with kidney dysfunction, supports the relevance of this biology to human disease. Renal C3 and CFB should therefore be evaluated prospectively as biomarkers of pathway activity, disease trajectory and treatment response, including for patient stratification and response assessment in complement-directed trials. More broadly, our findings suggest that the intracellular redeployment of evolutionarily ancient immune proteins provides a mechanism through which conserved inflammatory pathways acquire cell-and organ-specific functions. Understanding both the subcellular activities of these proteins and the tissue-specific signals that induce them may be necessary to translate this expanded view of complement biology into appropriately targeted therapies.

## Limitations of the study

Several limitations determine important next steps. Inefficient deletion with the available *Cfb* knockout prevented direct genetic testing of tubular FB *in vivo*, while loss of viability following *Cfb* silencing restricted complete loss-of-function studies in primary cells. The proposed DNA-binding interface is supported by computational modeling and mutational analysis but will ultimately require a dedicated study for detailed direct structural visualization. We have also not established how FB enters the nucleus, is activated intracellularly in the apparent absence of FD, whether in the nucleus or cytoplasm, or the precise mechanism and subcellular sites of FB-dependent C3 processing, including the identity of the C3-derived fragments generated. We have also not fully defined the biochemical sequence linking FB activation to DNA binding and the mechanism of histone processing. Finally, the human analyses performed here are cross-sectional and require prospective studies to confirm causality and prognostic utility.

## Supplementary Figure Legends

**Figure S1. The healthy kidney has a distinct complement transcriptional profile at steady state. A,** Heatmap showing expression of complement genes across individual samples from 13 perfused mouse tissues in E-MTAB-6081. Scores for the combined complement-pathway gene set and KEGG complement pathway are shown above. Liver and kidney samples are highlighted in red. **B**, Hierarchically clustered matrix of pairwise Pearson correlations between tissue samples based on complement-gene expression profiles. Liver and kidney samples show distinct complement-expression profiles relative to other tissues. Color bars denote tissue identity.

**Figure S2. Complement-pathway responses across experimental models of kidney injury.** Meta-analysis of publicly available mouse kidney injury transcriptomes. Bubble plots show enrichment of the indicated complement pathways (left) and expression of individual complement genes (right) for each injury model and time point. Bubble size denotes statistical significance. Color indicates normalized enrichment score (NES) for pathways and log2 fold change for individual genes. GEO accession numbers are shown on the right. K, KEGG; R, Reactome; WP, WikiPathways; H, Hallmark.

**Figure S3. Annotation and complement expression in the FAN kidney single-nucleus atlas. A,** Dot plot showing expression of selected lineage and cell-identity markers used to annotate kidney populations identified by snRNA-seq. Dot size indicates the percentage of nuclei expressing each gene, and color indicates scaled average expression. **B**, Density maps showing the distribution of kidney nuclei within the shared UMAP embedding at days 0, 2, 7 and 14 after folic acid administration. Color indicates normalized density; proximal tubular (PT) and immune-cell (Imm) regions are indicated. **C**, UMAP feature plots showing expression of *C3*, *Cfb* and *C3ar1*. **D**, Pairwise Pearson correlation matrix of complement-component gene expression across the major kidney cell populations (left) and relationship between mean C3 and Cfb expression across the same cell populations (right). PT-reactive1, PT-reactive2 and PT-reactive3 populations are labeled and shown in red; other major kidney cell populations are shown in blue. snRNAseq data in **Figure S3** are from n=1 vehicle and n=2 folic acid treated mice per timepoint.

**Figure S4. Differential protein abundance in tubular-cell culture supernatants following FA treatment.** Volcano plot showing differential protein abundance in in cell-culture supernatants from primary renal tubular epithelial cells following FA versus vehicle treatment. Proteins meeting the predefined differential-abundance thresholds are shown in red (increased with FA) or blue (decreased with FA), with C3 and FB indicated. Dashed lines denote the thresholds used to define differentially abundant proteins. Data are from n=2 independent experiments.

**Figure S5. Molecular and temporal characterization of reactive proximal tubular states after kidney injury. A**, UMAP feature plots showing expression of proximal tubular identity genes (*Lrp2*, *Slc5a12*, *Slc22a6* and *Slc7a13*) and injury-associated genes (*C3*, *Havcr1*, *Lcn2* and *Vcam1*) across reclustered proximal tubular populations. **B**, Heatmap showing expression of curated genes associated with injured, maladaptive and differentiated proximal tubular states across PT-reactive and differentiated proximal tubular populations, as well as *C3* and *Cfb*. **C**, Representative kidney section from a vehicle-treated *C3*-tdTomato reporter mouse showing *C3*-tdTomato and VCAM1 expression. Scale bar, 250μm. Image is representative of n=3 independent experiments. **D**, Relative abundance of differentiated and reactive proximal tubular populations at days 0, 2, 7 and 14 after folic acid administration. **E**, RNA-velocity stream plots showing inferred local cell-state trajectories across proximal tubular populations at days 2, 7 and 14 after folic acid administration. Streamlines and arrowheads indicate the local inferred direction of transcriptional state change within the shared proximal tubular embedding. snRNAseq data in **Figure S5** are from n=1 vehicle and n=2 folic acid treated mice per timepoint.

**Figure S6. Reactive proximal tubular cells are prominent participants in inferred intercellular communication after kidney injury. A-B**, CellChat-derived incoming and outgoing interaction strengths for the indicated kidney cell populations considering all inferred ligand-receptor interactions (**A**); and incoming and outgoing interaction strengths restricted to complement-mediated interactions (**B**). Point size in **A-B** indicates the number of inferred ligand-receptor interactions. Imm, immune cells; IC, intercalated cells; Fib, fibroblasts; EC, endothelial cells; CD, collecting duct cells; Ren, renin-lineage cells; PT, proximal tubular cells; PT-reactive, reactive proximal tubular cells; Pod, podocytes; PEC, parietal epithelial cells; LH, loop of Henle cells. **C**, Representative kidney section from a vehicle-treated *C3*-tdTomato reporter mouse showing *C3*-tdTomato reporter signal and C3aR1 staining. Scale bar, 50μm. Image is a representative example from n=3 experiments performed. snRNAseq data in **Figure S6** are from n=5 folic-acid treated mice.

**Figure S7. IL-1R pathway is enriched in reactive proximal tubular cells and contributes to C3-FB induction. A**, Gene-set enrichment analysis comparing reactive with differentiated proximal tubular populations. Pathways enriched differentiated and reactive proximal tubular cells are shown in blue and red, respectively; cytokine signaling is highlighted. **B**, UMAP feature plot showing the BIOCARTA IL-1R pathway module score across proximal tubular populations. **C**, Heatmap showing expression of IL-1R pathway genes in proximal tubular cells at days 0, 2, 7 and 14 after folic acid injury. Bars at right indicate the maximal expression of each genes actoss the time course. **D**, Relative *Il1r1*, *C3* and *Cfb* mRNA expression in unstimulated cells and IL-1β-stimulated proximal tubular epithelial cells carrying control or *Il1r1*-targeting gRNA. Bars show mean ± s.e.m. *p<0.05; **p<0.01; ****p<0.0001. snRNAseq data in **Figure S7** are from n n=1 vehicle and n=2 folic acid treated mice per timepoint.

**Figure S8. AP-1 and NF-κB regulon activity is enriched in reactive proximal tubular states. A**, Combined AP-1 and NF-κB regulon activity across differentiated and reactive tubular populations. **B**, UMAP feature plots showing SCENIC-inferred activity of JUN, JUNB, REL, RELA and RELB regulons (top) and expression of the corresponding AP-1/NF-κB family genes together with *C3* and *Cfb* (bottom). snRNAseq data in **Figure S8** are from n=1 vehicle and n=2 folic acid treated mice per timepoint.

**Figure S9. IL-1β remodels c-JUN and RELA occupancy at injury-responsive genes. A**, Aggregate c-JUN and RELA CUT&Tag signal centered on called peaks in untreated and IL-1β-treated tubular epithelial cells, together with matched IgG controls. Numbers below indicate the number of called peaks. **B**, Aggregate plots and heatmaps showing H3K27Ac regions repressed or induced following IL-1β stimulation. Numbers below indicate the number of called peaks. **C-D**, Aggregate plots and heatmaps showing c-JUN (**C**) and RelA (**D**) peak signal repressed or induced by IL-1β at H3K27Ac peaks. Numbers below indicate the number of regions in each group. **E**, Network representation of IL-1β-regulated genes associated specifically with RELA, with both RELA and c-JUN, or specifically with c-JUN. Red and blue denote IL-1β-induced and repressed genes, respectively; selected genes including *C3* and *Cfb* are highlighted in yellow. **F**, T7 endonuclease assay confirming CRISPR-Cas9 editing of *Jun*, *Rela* and *Nfkb1*. Shown is a representative example from n=3 experiments. **G**, Relative *C3* and *Cfb* mRNA expression 48h following CRISPR-Cas9 targeting of *Jun*, *Rela* or *Nfkb1*, relative to control gRNA, in IL-1β-stimulated tubular cells. Data are from n=3 experiments. Bars show mean ± s.e.m. *p<0.05; **p<0.01.

**Figure S10. IL-1β induces C3 and FB protein and reveals intracellular complement pools in tubular epithelial cells. A,** Native immunoblot showing C3 in cell lysates from primary tubular epithelial cells with or without IL-1β stimulation for the indicated times. Recombinant human C3 is shown as a control. Because proteins were resolved under native, non-denaturing conditions, standard molecular-weight markers are not applicable and are therefore not shown. β-actin serves as a loading control. **B,** Representative immunoblots showing C3 (top) and FB (bottom) in cell-culture supernatants from vehicle-or IL-1β-treated primary tubular epithelial cells. Recombinant human C3 and FB are shown as controls. Shown are examples from n=3 experiments. **C,** Representative confocal images of IL-1β-stimulated tubular epithelial cells showing C3 together with calnexin, GPP130, EEA1, RAB7, LAMP2, LC3 or TOM20. Phalloidin and DAPI mark F-actin and nuclei, respectively. Scale bars, 10μm. Images are representative examples from n=2 experiments. **D**, Representative confocal image showing C3 and FB in IL-1β -stimulated tubular epithelial cells. Phalloidin and DAPI mark F-actin and nuclei, respectively. Scale bar, 10μm. Shown is a representative example from n=2 experiments. **E**, Total-protein staining of cell-culture supernatants from tubular epithelial cells treated with IL-1β in the presence of the indicated concentrations of cpFBi. The samples correspond to those analyzed for C3a in **Fig. 4F**; a cropped region of this blot is shown beneath the C3a immunoblot in **Fig. 4F**.

**Figure S11. Canonical *C3* and *Cfb* transcripts are expressed in proximal tubular cells. A**, Sashimi plots showing *C3* splice-junction usage in proximal tubular epithelial cells with or without IL-1β stimulation, together with annotated mouse *C3* transcript isoforms. **B**, Expression of the indicated *C3* transcripts in proximal tubular epithelial cells with or without IL-1β stimulation (left) and in mouse liver (right). **C**, Sashimi plots showing *Cfb* splice-junction usage in proximal tubular epithelial cells with or without IL-1β treatment, together with annotated mouse *Cfb* transcript isoforms. **D**, Expression of the indicated *Cfb* transcript isoforms in proximal tubular epithelial cells with or without IL-1β stimulation (left) and in mouse liver (right). **E**, Relative contribution of the predominant *C3* (left) and *Cfb* (right) transcript isoforms to total transcript abundance in untreated and IL-1β-stimulated proximal tubular cells and liver. **F**, C3 mRNA expression in IL-1β-stimulated TKPTS cells treated with cpFBi or DMSO vehicle, shown as log2 RPKM. Bars show mean ± s.e.m.; individual points represent independent samples. **p<0.01; ****p<0.0001

**Figure S12. Tubular *C3* deletion impairs metabolic adaptation following IL-1β stimulation. A,** Representative immunoblot (left) and quantification (right) of C3 in cell-culture supernatants from primary tubular epithelial cells isolated from *C3*^fl/fl^ and *C3*^fl/fl^*Pax8Cre*^+/−^ mice, with or without IL-1β stimulation. Recombinant human C3 is shown as a control. Shown is one representative example from n=2 experiments. **B**, Basal OCR in IL-1β-stimulated primary proximal tubular epithelial cells from *C3*^fl/fl^ and *C3*^fl/fl^*Pax8Cre*^+/−^ mice. **E**, ECAR during mitochondrial stress testing with sequential addition of oligomycin, FCCP and rotenone in IL-1β-stimulated primary proximal tubular epithelial cells from *C3*^fl/fl^ and *C3*^fl/fl^*Pax8Cre*^+/−^ tubular epithelial cells. Data in **B-C** are from n=2 independent experiments. **E**, Basal ECAR, maximal ECAR and glycolytic reserve derived from **C**. Bars show mean ± s.e.m. **p<0.01.

**Figure S13. Nuclear localization and DNA binding of FB. A**, Representative confocal microscopy showing nuclear FB in both Ki67-positive and Ki67-negative IL-1β-treated renal tubular epithelial cells. Ki67 is shown in green, FB in red, phalloidin in magenta and DAPI in blue. Scale bar, 10μm. **B-C**, FB abundance in cytoplasmic and nuclear-enriched fractions from two independently published mouse cardiac nuclear-proteomics datasets (Fang *et al.*, 2025 (**B**) and Eslami *et al.*, 2026 (**C**)). Values are shown using the abundance measures reported in the respective datasets. **D**, Genomic distribution of FB-enriched CUT&Tag regions among promoter, intragenic and intergenic sequences. **E**, EMSA showing concentration-dependent binding of recombinant FB to a double-stranded oligonucleotide containing a sequence motif enriched within FB-bound genomic regions by *de novo* motif analysis (**Methods**). DNA and protein are visualized by SYBR Green and SYPRO Ruby staining, respectively; c-MYC serves as a positive DNA-binding control. Shown is a representative experiment from n=2 independent experiments. Mean apparent *K*_D_, 0.26 μM.

**Figure S14. A conserved C-terminal surface contributes to FB DNA binding. A,** GraphSite-predicted DNA-contacting residues across AlphaFold2-predicted human wild-type FB and the human FB^11xAlaMut^ variant, shown together with AlphaFold2 model confidence, protein domains and residue conservation. The dotted line denotes the GraphSite prediction threshold used to classify candidate DNA-contacting residues. Predicted residues homologous to DNA-contacting residues identified in mouse FB are indicated in bold, and asterisks denote residues belonging to the serine-protease catalytic triad. Please note that GraphSite analysis of FB^11xAlaMut^ additionally identified R175 as a candidate DNA-contacting residue that was not predicted in wild-type FB, suggesting a potential alternative DNA-contacting site that could contribute to residual binding after disruption of the principal predicted interface. **B**, AlphaFold2 structural model of human FB showing the predicted C-terminal DNA-binding interface and selected candidate DNA-contacting residues. **C**, AlphaFold2-predicted structures of wild-type FB and FB^11xAlaMut^ shown in two orientations (top), with enlarged views of the predicted C-terminal DNA-binding interface (bottom). Residues constituting the predicted interface in wild-type FB and the corresponding alanine substitutions in FB^11xAlaMut^ are highlighted (red and orange, respectively). **D**, Structural superposition of wild-type FB and FB^11xAlaMut^ colored according to Cα root-mean-square deviation (RMSD). **E**, EMSAs showing concentration-dependent DNA binding by the indicated recombinant FB variants. DNA and protein are visualized by SYBR Green and SYPRO Ruby staining respectively. Shown are representative examples from n=2 experiments. Apparent *K*_D_ values are shown below each assay.

**Figure S15. FB inhibition alters promoter H3K27 acetylation and reduces nucleosome-associated DNA protection. A,** Aggregate H3K27Ac CUT&RUN signal at regions showing decreased (left; n=5,753) or increased (right; n=23,227) H3K27Ac following FB inhibition, with corresponding isotype-control signal. **B**, Aggregate FB and isotype-control CUT&RUN signal at H3K27Ac-positive regions associated with TSSs (n=8,789) or distal regulatory regions (n=31,993). **C**, Aggregate MNase-seq signal centered on TSSs in IL-1β-treated tubular epithelial cells, comparing IL-1β alone, IL-1β plus DMSO and IL-1β plus cpFBi. **D**, Autocorrelation of MNase-seq signal after DMSO or cpFBi treatment, showing largely preserved nucleosome phasing. **E**, Genome-browser tracks showing FB and H3K27Ac CUT&RUN signal at the *Pax8* locus after DMSO or cpFBi treatment. **F**, KLF-like and TEAD-like sequence motifs enriched among FB-associated active promoters showing ≥2-fold increased H3K27Ac following FB inhibition, using the remaining FB-associated active promoters as background. Motif frequencies and enrichment p-values are shown. **G**, Frequency of an AP-1/bZIP-family motif among FB-associated promoters showing ≥2-fold increased H3K27Ac following FB inhibition compared with the remaining FB-associated active promoters. Fold enrichment is indicated. **H**, Pathway enrichment analysis of genes associated with the 500 FB-associated active promoters showing the greatest increase in H3K27Ac after FB inhibition. The x-axis indicates statistical significance, bubble size indicates fold enrichment and color indicates the number of genes in the overlap.

**Figure S16. Tubular C3 deletion reduces renal macrophage accumulation and alters macrophage transcriptional state and intercellular signaling after kidney injury. A**, T7 endonuclease assay confirming CRISPR-Cas9 editing of *Cfb* in tubular epithelial cells following electroporation with *Cfb*-targeting or scrambled-control gRNA (left), and quantification of viable-cell recovery following *Cfb* targeting relative to scrambled control (right). Primer sets 1 and 2 amplify regions encompassing *Cfb* exons 1 and 2, respectively. Bars show mean ± s.e.m.; individual values are shown. ***p<0.001. **B**, UMAP visualization and annotation of immune-cell populations identified by snRNA-seq. **C**, UMAP visualization and annotation of immune-cell populations identified by CD45^+^ cell-enriched scRNA-seq. **D**, Relative contribution of *C3*^fl/fl^ and *C3*^fl/fl^*Pax8Cre*^+/−^ nuclei to the indicated renal cell populations in the snRNA-seq dataset (left), with cluster size shown at right. **E**, Relative contribution of the indicated genotypes to immune-cell populations in the CD45^+^ cell-enriched scRNA-seq (left), with cluster size shown at right. Macrophages showed reduced representation in *C3*^fl/fl^*Pax8Cre*^+/−^ kidneys. Data in **B** and **D** are from n=3 mice per genotype; data in **C** and **E** are from n = 2 mice per genotype. **F,** Volcano plot showing differential gene expression in macrophages from *C3*^fl/fl^ and *C3*^fl/fl^*Pax8Cre*^+/−^ kidneys. Differentially expressed genes were defined by absolute log2-fold change ≥0.2 and adjusted p< 0.05. Genes expressed at higher levels in tubular C3-KO macrophages are shown in red and those expressed at higher levels in control macrophages in blue; selected genes are indicated. Data are from n=2 mice per genotype. **G**, Total number (left) and mean strength (right) of CellChat-inferred cell-cell interactions in *C3*^fl/fl^ and *C3*^fl/fl^*Pax8Cre*^+/−^ kidneys. **H**, Differential incoming and outgoing CellChat-inferred signaling involving macrophages (left) and PT-reactive cells (right) between *C3*^fl/fl^ and *C3*^fl/fl^*Pax8Cre*^+/−^ kidneys. Selected signaling pathways are indicated.

**Figure S17. Human kidney C3 and FB associate with reactive proximal tubular states and renal dysfunction. A**, UMAP visualization and annotation of all cell populations in the human KPMP single-cell transcriptomic dataset. aFib, adaptive/maladaptive fibroblast; aPT, adaptive/maladaptive proximal tubular epithelial cell; aTAL1 and aTAL2, adaptive/maladaptive thick ascending limb cell clusters 1 and 2; B, B cell; C-TAL, cortical thick ascending limb cell; cDC, classical dendritic cell; CNT, connecting tubule cell; CNT-IC-A, connecting tubule intercalated cell type A; CNT-PC, connecting tubule principal cell; cycEC, cycling endothelial cell; cycEPI, cycling epithelial cell; cycMNP, cycling mononuclear phagocyte; cycT, cycling T cell; dC-TAL, degenerative cortical thick ascending limb cell; dCNT, degenerative connecting tubule cell; dCNT-PC, degenerative connecting tubule principal cell; DCT1, distal convoluted tubule cell type 1; dDCT, degenerative distal convoluted tubule cell; dEC-PTC, degenerative peritubular capillary endothelial cell; dIC-A, degenerative intercalated cell type A; dPC, degenerative principal cell; dPT, degenerative proximal tubular epithelial cell; dPT/DTL, degenerative proximal tubular epithelial cell/descending thin limb cell; DTL1, descending thin limb cell type 1; dVSMC, degenerative vascular smooth muscle cell; EC-AEA, afferent/efferent arteriole endothelial cell; EC-GC, glomerular capillary endothelial cell; EC-LYM, lymphatic endothelial cell; EC-PTC, peritubular capillary endothelial cell; Fib, fibroblast; IC-A and IC-B, intercalated cells types A and B; Mac-M2, M2 macrophage; Mast, mast cell; MC, mesangial cell; MDC, monocyte-derived cell; Mon, monocyte; M-TAL, medullary thick ascending limb cell; MyoF, myofibroblast; ncMon, non-classical monocyte; NK1 and NK2, natural killer cell clusters 1 and 2; NKT, natural killer T cell; PC, principal cell; pDC, plasmacytoid dendritic cell; PEC, parietal epithelial cell; PL, plasma cell; Pod, podocyte; PT-S1/S2, proximal tubular epithelial cells segments 1 and 2; PT-S3, proximal tubular epithelial cell segment 3; Ren, renin-positive juxtaglomerular granular cell; T, T cell; T-Cyt, cytotoxic T cell; T-reg, regulatory T cell; tPC-IC, transitional principal-intercalated cell; VSMC/P, vascular smooth muscle cell/pericyte. **B**, Heatmap showing the mouse-derived PT-reactive module score across KPMP cell populations in living donor, AKI and CKD kidneys. **C**, UMAP feature plots showing *C3*, *CFB*, *VCAM1* and *C3AR1* expression in living-donor, CKD and AKI samples. **D**, Heatmaps showing expression of complement components (top) and receptors/regulators (bottom) across the indicated human kidney diseases and healthy controls. AAV, ANCA-associated vasculitis; DKD, diabetic kidney disease; FSGS, focal segmental glomerulosclerosis; HTN, hypertensive nephropathy; IgAN, IgA nephropathy; LN, lupus nephritis; MCD, minimal change disease; MN, membranous nephropathy; TMD, thin basement membrane disease. **E**, Receiver-operating-characteristic (ROC) curves for renal *C3* and *CFB* expression and eGFR distinguishing kidney disease samples (n=165) from healthy controls (n=21). AUC and p-values are shown.

## Supplementary Tables

**Table S1. Public Datasets used in this study.** Publicly available transcriptomic and proteomic datasets analyzed in this study.

Table S2. CRISPR guides and their genomic coordinates used in this study.

Table S3. Curated gene sets used in this study.

**Table S4. Gene expression in wild-type proximal tubular epithelial cells after IL-1β treatment.** Gene expression values (RPKM) are shown for *C3*^fl/fl^ mouse proximal tubular epithelial cells treated with carrier or IL-1β, together with log2 fold change (log2FC) and statistical comparisons. Differentially expressed genes (DEGs) were defined as those with an absolute log2FC ≥ 1 and FDR < 0.05. DEGs are denoted as 1 for IL-β-induced genes and −1 for IL-1β-repressed genes.

**Table S5. Gene-expression and pathway analyses of C3- and FB-dependent transcriptional programs in renal tubular epithelial cells. A,** Gene expression values (RPKM) in IL-1β-treated *C3*^fl/fl^ and *C3*^fl/fl^*Pax8Cre* proximal tubular epithelial cells. **B,** Gene expression values (RPKM) in IL-1β-treated TKPTS cells cultured with cell-permeable FB inhibitor (cpFBi) or carrier (DMSO). **C,** Biological pathway enrichment analyses of MSigDB Hallmark and Canonical gene sets, with significantly enriched pathways defined as an absolute normalized enrichment score (NES) ≥ 1.5 and *p* < 0.05. Shown are comparisons between *C3*^fl/fl^ and *C3*^fl/fl^*Pax8Cre* renal tubular epithelial cells from the folic acid nephropathy model (top), IL-1β-treated *C3*^fl/fl^ and *C3*^fl/fl^*Pax8Cre* proximal tubular epithelial cells *in vitro* (middle), and IL-1β-treated TKPTS cells cultured with cpFBi or DMSO carrier (bottom).

**Table S6. Tubular proteomics. A-B,** Proteins identified by mass spectrometry (**A**) and differentially abundant proteins (**B**) in cell-culture supernatants from vehicle-or folic-acid primary proximal tubular epithelial cells. DAP, differentially abundant protein; +1 indicates increased abundance and −1 decreased abundance following folic acid treatment relative to vehicle.

**Table S7. IL-1β-induced AP-1/NF-κB chromatin occupancy and transcriptional regulation. A-B**, differential c-JUN (**A**) and RELA (**B**) CUT&Tag peaks comparing control and IL-1β-treated cells, with genomic coordinates, annotation and assigned gene. **C**, differential H3K27Ac regions, comparing control-treated and IL-1β-treated cells, with genomic coordinates, annotation and assigned gene. Association with IL-1β-regulated gene expression (DEG) is indicated as +1, −1 or 0 for genes that are upregulated, downregulated or not differentially expressed, respectively. **D**, complete list of 5983 genomic regions showing increased H3K27Ac after IL-1β.

**Table S8. Factor B-associated active promoter regions.** The table lists 1,299 Factor B-bound, H3K27Ac-positive transcription start-site (TSS) promoters. For each region, the change in MNase signal (ΔMNase) is reported as percentage change.

**Table S9. Differential gene expression analysis of macrophages from tubular C3-KO and control mice with folic acid nephropathy.** Log2 fold change (log2FC) and statistical comparisons are shown for detected genes from scRNA-seq analysis of macrophages from *C3*^fl/fl^*Pax8Cre* tubular C3-KO mice and *C3*^fl/fl^ control mice. Log2FC is calculated as tubular C3-KO versus control. Differentially expressed genes (DEGs) were defined as those with an absolute log2FC ≥ 0.2 and adjusted p-value < 0.05. In the DEG-direction column, 1 indicates higher expression in tubular C3-KO macrophages and −1 indicates lower expression in tubular C3-KO macrophages (higher expression in control macrophages).

## Supplementary Videos

**Video S1. Nuclear localization of FB in IL-1β-stimulated tubular epithelial cells, related to Figures 5A and S14A.** Three-dimensional rendering of a confocal z-stack of IL-1β-stimulated tubular epithelial cells, demonstrating prominent localization of FB within DAPI-defined nuclei. FB is shown in red and DAPI in blue.

**Video S2. Structural models of the predicted FB DNA-binding interface, related to Figures 5D-E and S15A-B. A-B,** Structural visualization of AlphaFold2-predicted mouse (**A**) and human (**B**) FB models highlighting GraphSite-predicted DNA-contacting residues clustered within the C-terminal peptidase domain. The predicted residues define a candidate DNA-binding groove (shown in red) in proximity to and including some of the residues of the serine-protease catalytic triad.

## Acknowledgments

We acknowledge the computational resources of the NIH High-Performance Computing (HPC) Biowulf cluster (https://hpc.nih.gov) and the Hummel-2 HPC cluster at the University of Hamburg, which was funded by the Deutsche Forschungsgemeinschaft (DFG, German Research Foundation; grant 498394658). Molecular graphics and analyses were performed with UCSF ChimeraX, developed by the Resource for Biocomputing, Visualization, and Informatics at the University of California, San Francisco, with support from National Institutes of Health grant R01-GM129325 and the Office of Cyber Infrastructure and Computational Biology, National Institute of Allergy and Infectious Diseases. This research was supported by the National Institute of General Medical Sciences of the NIH (grant R35GM138283 to M.K.), the National Institute of Diabetes and Digestive and Kidney Diseases (grant R01DK12262401A1 to D.P.), the Deutsche Forschungsgemeinschaft (fellowship FR 3851/2-1 to T.F. and SFB 1192 project B6 II to T.F.). J.J. was supported by the Else Kroener-Fresenius Stiftung Promotionskolleg iPRIME, Hamburg (iPRIME_2021_EKPK.10). This research was also supported by the Intramural Research Programs of the National Institute of Diabetes and Digestive and Kidney Diseases (ZIA/DK075149 to B.A.) and the National Heart, Lung, and Blood Institute (ZIA/HL006223 to C.K.) of the NIH. The project underlying this report was also funded by the German Federal Ministry of Research, Technology and Space (BMFTR; grant 01EO2106); responsibility for the content of this publication lies with the authors. The contributions of the NIH author(s) are considered Works of the United States Government. The findings and conclusions presented in this paper are those of the author(s) and do not necessarily reflect the views of the NIH or the U.S. Department of Health and Human Services.

## Notes

### Competing Interest Statement

The authors have declared no competing interest.

