## Supplementary Figures for "Intracellular complement Factor B couples inflammatory sensing to metabolism, chromatin regulation and maladaptive tissue repair"

### Figure S1

**A**

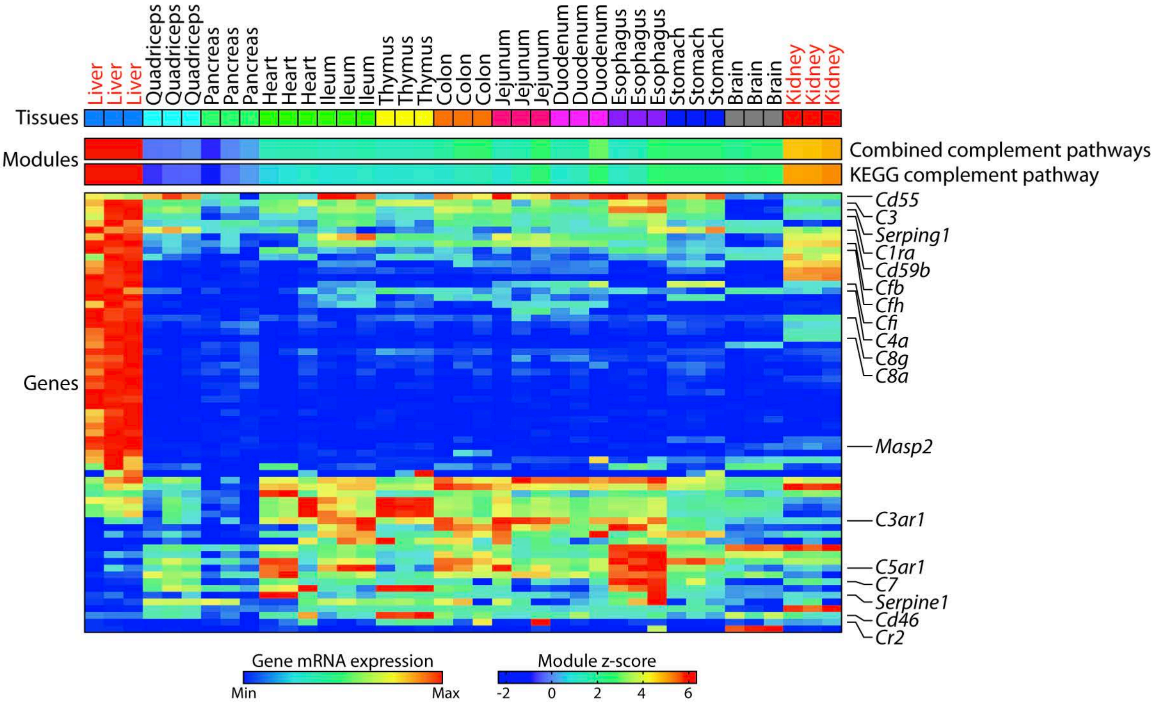

Tissues
Modules
Genes
Combined complement pathways
KEGG complement pathway
Cd55
C3
Serping1
C1ra
Cd59b
Cfb
Cfh
Cfi
C4a
C8g
C8a
Masp2
C3ar1
C5ar1
C7
Serpine1
Cd46
Cr2
Gene mRNA expression
Min
Max
Module z-score
-2
0
2
4
6


**B**

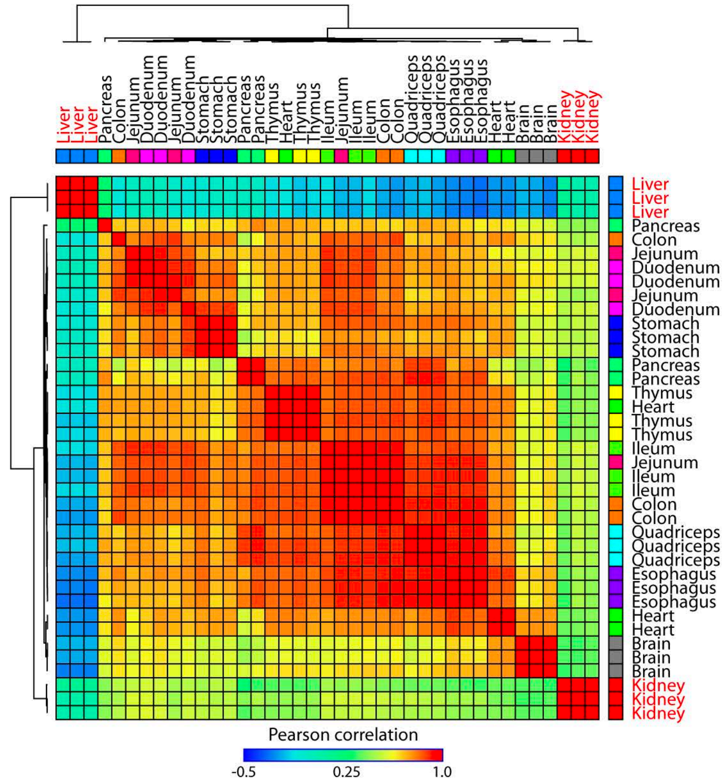

Pearson correlation
-0.5
0.25
1.0

**Figure S2**

**A**

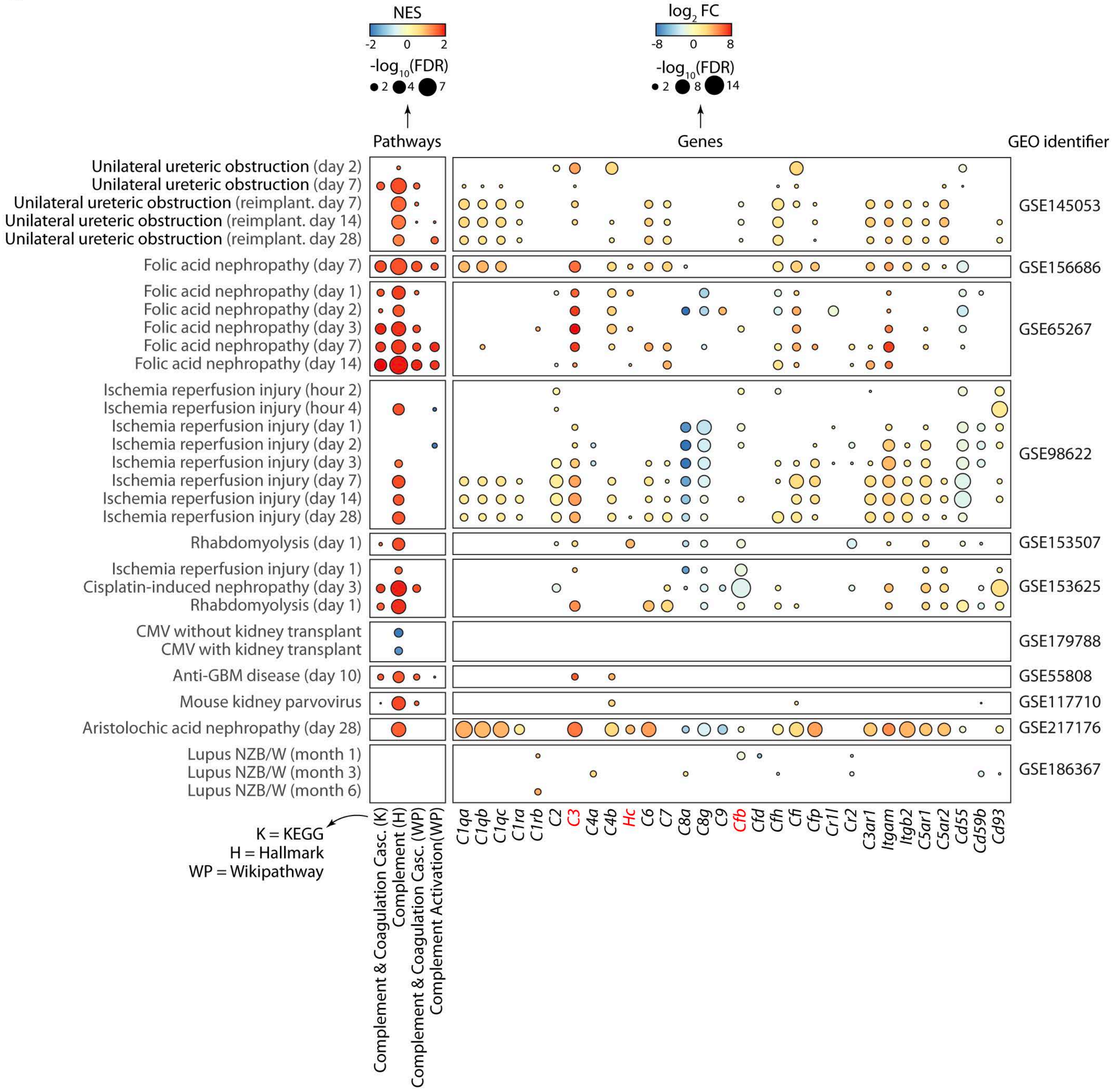
NES
-2
0
2
-log10(FDR)
2
4
7
Pathways
log2 FC
-8
0
8
-log10(FDR)
2
8
14
Genes
GEO identifier
Unilateral ureteric obstruction (day 2)
Unilateral ureteric obstruction (day 7)
Unilateral ureteric obstruction (reimplant. day 7)
Unilateral ureteric obstruction (reimplant. day 14)
Unilateral ureteric obstruction (reimplant. day 28)
GSE145053
Folic acid nephropathy (day 7)
GSE156686
Folic acid nephropathy (day 1)
Folic acid nephropathy (day 2)
Folic acid nephropathy (day 3)
Folic acid nephropathy (day 7)
Folic acid nephropathy (day 14)
GSE65267
Ischemia reperfusion injury (hour 2)
Ischemia reperfusion injury (hour 4)
Ischemia reperfusion injury (day 1)
Ischemia reperfusion injury (day 2)
Ischemia reperfusion injury (day 3)
Ischemia reperfusion injury (day 7)
Ischemia reperfusion injury (day 14)
Ischemia reperfusion injury (day 28)
GSE98622
Rhabdomyolysis (day 1)
GSE153507
Ischemia reperfusion injury (day 1)
Cisplatin-induced nephropathy (day 3)
Rhabdomyolysis (day 1)
GSE153625
CMV without kidney transplant
CMV with kidney transplant
GSE179788
Anti-GBM disease (day 10)
GSE55808
Mouse kidney parvovirus
GSE117710
Aristolochic acid nephropathy (day 28)
GSE217176
Lupus NZB/W (month 1)
Lupus NZB/W (month 3)
Lupus NZB/W (month 6)
GSE186367
K = KEGG
H = Hallmark
WP = Wikipathway
Complement & Coagulation Casc. (K)
Complement (H)
Complement & Coagulation Casc. (WP)
Complement Activation(WP)
C1qa
C1qb
C1qc
C1ra
C1rb
C2
C3
C4a
C4b
Hc
C6
C7
C8a
C8g
C9
Cfb
Cfd
Cfh
Cfi
Cfp
Cr1l
Cr2
C3ar1
Itgam
Itgb2
C5ar1
C5ar2
Cd55
Cd59b
Cd93

**Figure S3**

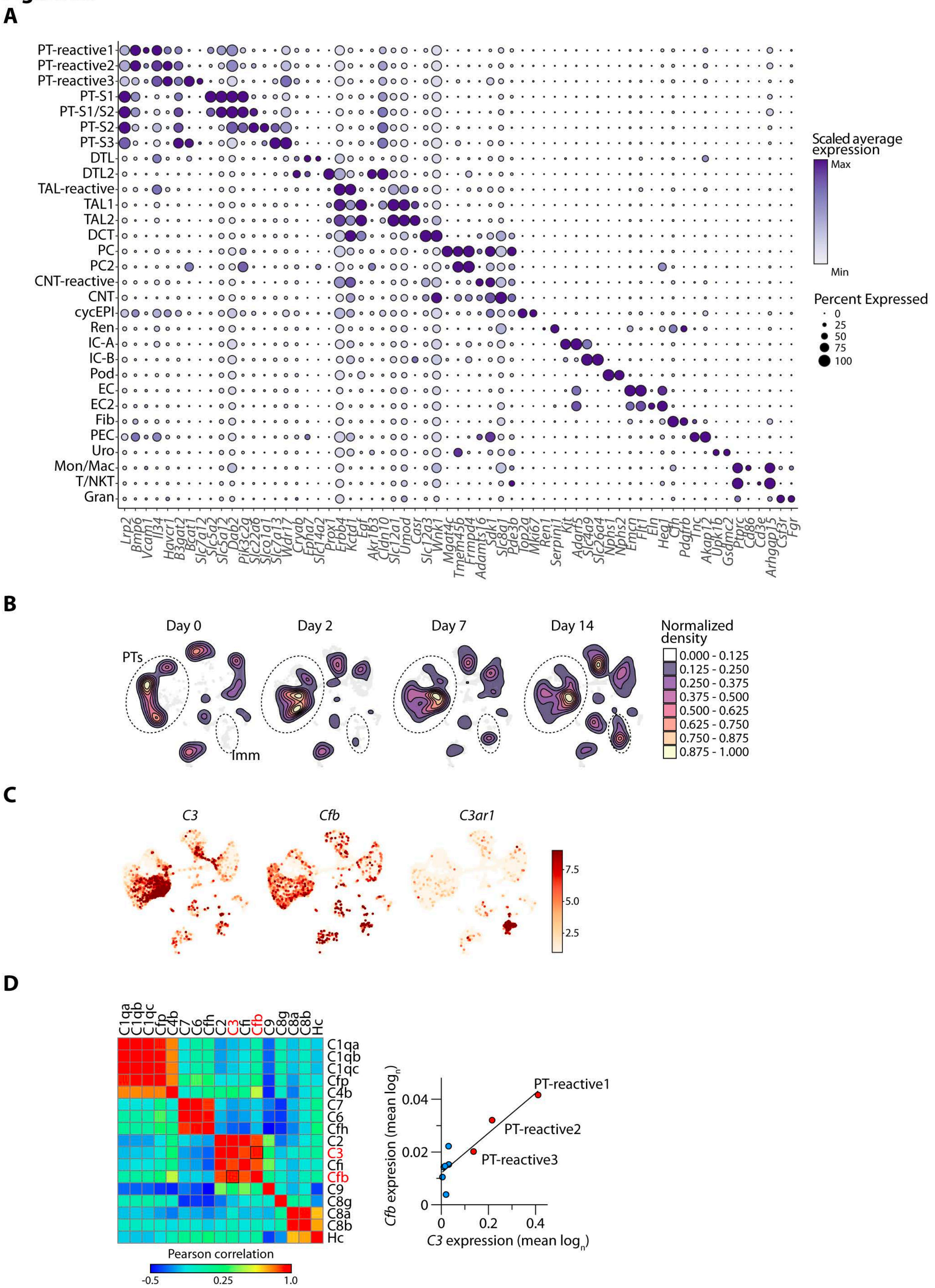
A
PT-reactive1
PT-reactive2
PT-reactive3
PT-S1
PT-S1/S2
PT-S2
PT-S3
DTL
DTL2
TAL-reactive
TAL1
TAL2
DCT
PC
PC2
CNT-reactive
CNT
cycEPI
Ren
IC-A
IC-B
Pod
EC
EC2
Fib
PEC
Uro
Mon/Mac
T/NKT
Gran
Lrp2 Bmp6 Vcam1 Il34 Havcr1 B3gat2 Bcat1 Slc7a12 Slc5a2 Slc5a12 Dab2 Pik3c2g Slc22a6 Slco1a1 Slc7a13 Wdr17 Cryab Epha7 Slc14a2 Prox1 Erbb4 Kctd1 Egf Akr1b3 Cldn10 Slc12a1 Umod Casr Slc12a3 Wnk1 Mgat4c Tmem45b Frmpd4 Adamts16 Sdk1 Slc8a1 Pde3b Top2a Mki67 Ren1 Serpini1 Kit Adgrf5 Slc4a9 Slc26a4 Nphs1 Nphs2 Emcn Flt1 Eln Heg1 Cfh Pdgfrb Tnc Akap12 Upk1b Gsdmc2 Ptprc Cd86 Cd3e Arhgap15 Csf3r Fgr
Scaled average expression
Max
Min
Percent Expressed
0
25
50
75
100
B
Day 0
Day 2
Day 7
Day 14
PTs
Imm
Normalized density
0.000 - 0.125
0.125 - 0.250
0.250 - 0.375
0.375 - 0.500
0.500 - 0.625
0.625 - 0.750
0.750 - 0.875
0.875 - 1.000
C
C3
Cfb
C3ar1
7.5
5.0
2.5
D
C1qa C1qb C1qc Cfp C4b C7 C6 Cfh C2 C3 Cfi Cfb C9 C8g C8a C8b Hc
C1qa
C1qb
C1qc
Cfp
C4b
C7
C6
Cfh
C2
C3
Cfi
Cfb
C9
C8g
C8a
C8b
Hc
Pearson correlation
-0.5
0.25
1.0
Cfb expresion (mean log$_n$)
0.04
0.02
0
PT-reactive1
PT-reactive2
PT-reactive3
0
0.2
0.4
C3 expression (mean log$_n$)

**Figure S4**

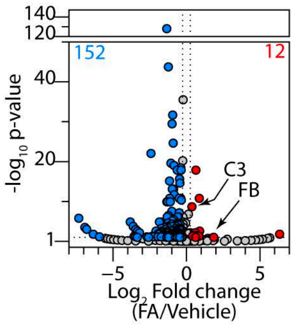

140
120
152
12
40
20
1
-log10 p-value
C3
FB
−5
0
5
Log2 Fold change
(FA/Vehicle)

### Figure S5

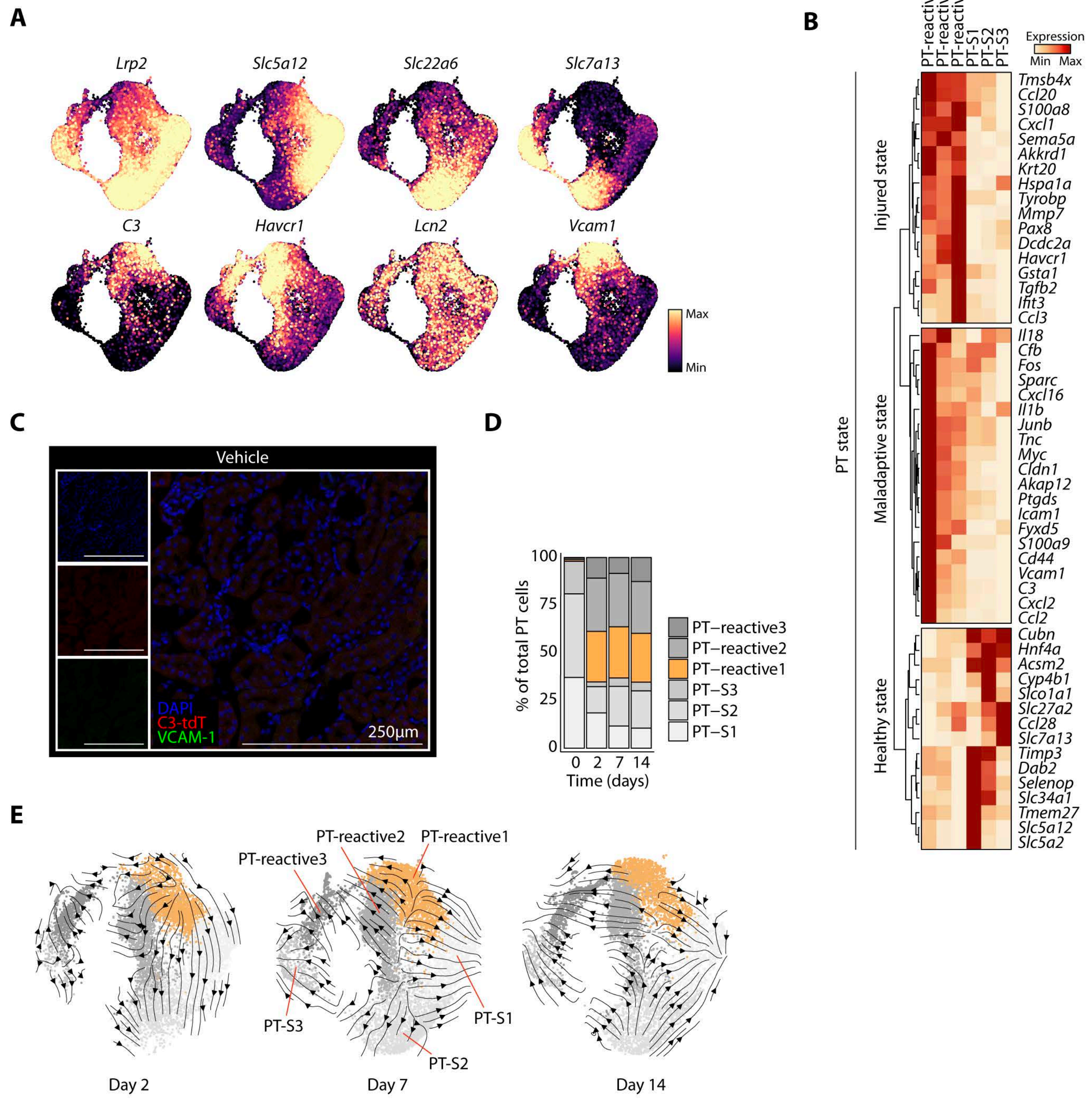
A
Lrp2
Slc5a12
Slc22a6
Slc7a13
C3
Havcr1
Lcn2
Vcam1
Max
Min
B
PT-reactive1
PT-reactive2
PT-reactive3
PT-S1
PT-S2
PT-S3
Expression
Min Max
PT state
Injured state
Maladaptive state
Healthy state
Tmsb4x
Ccl20
S100a8
Cxcl1
Sema5a
Akkrd1
Krt20
Hspa1a
Tyrobp
Mmp7
Pax8
Dcdc2a
Havcr1
Gsta1
Tgfb2
Ifit3
Ccl3
Il18
Cfb
Fos
Sparc
Cxcl16
Il1b
Junb
Tnc
Myc
Cldn1
Akap12
Ptgds
Icam1
Fyxd5
S100a9
Cd44
Vcam1
C3
Cxcl2
Ccl2
Cubn
Hnf4a
Acsm2
Cyp4b1
Slco1a1
Slc27a2
Ccl28
Slc7a13
Timp3
Dab2
Selenop
Slc34a1
Tmem27
Slc5a12
Slc5a2
C
Vehicle
DAPI
C3-tdT
VCAM-1
250µm
D
% of total PT cells
100
75
50
25
0
0 2 7 14
Time (days)
PT–reactive3
PT–reactive2
PT–reactive1
PT–S3
PT–S2
PT–S1
E
PT-reactive3
PT-reactive2
PT-reactive1
PT-S3
PT-S1
PT-S2
Day 2
Day 7
Day 14

Figure S6

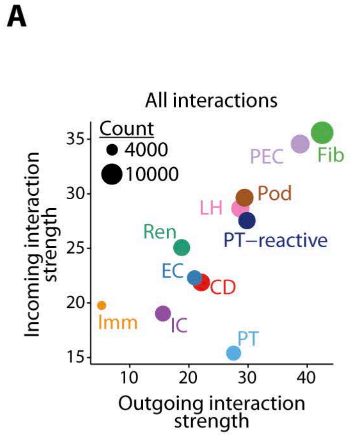

A
All interactions
Count
4000
10000
Incoming interaction strength
Outgoing interaction strength
PEC
Fib
Pod
LH
PT−reactive
Ren
EC
CD
Imm
IC
PT
35
30
25
20
15
10
20
30
40


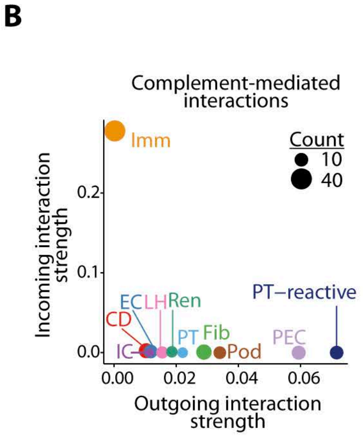

B
Complement-mediated interactions
Count
10
40
Imm
Incoming interaction strength
Outgoing interaction strength
EC
LH
Ren
CD
PT−reactive
IC
PT
Fib
Pod
PEC
0.2
0.1
0.0
0.00
0.02
0.04
0.06


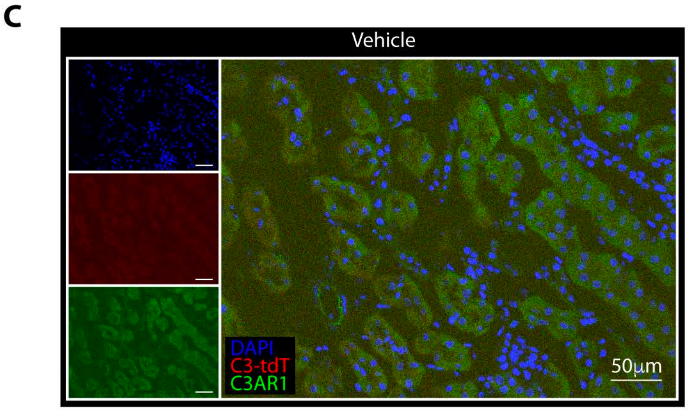

C
Vehicle
DAPI
C3-tdT
C3AR1
50µm

### Figure S7

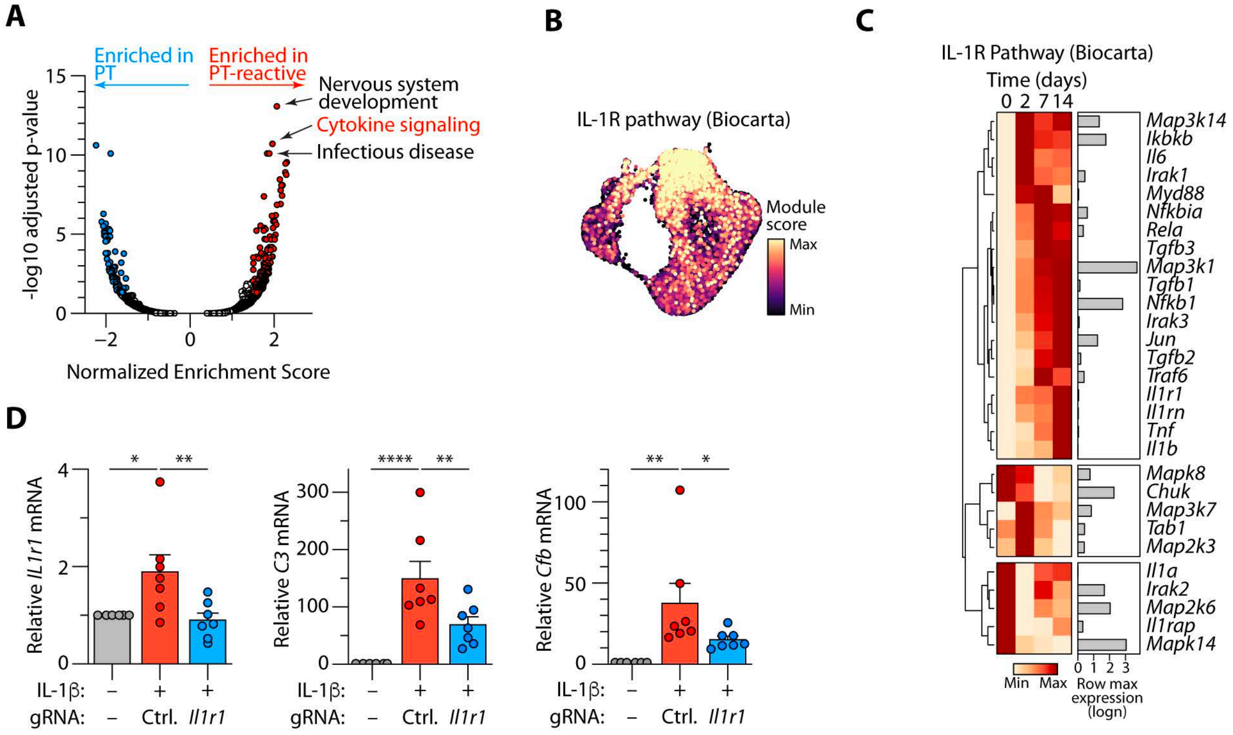
A
Enriched in PT
Enriched in PT-reactive
Nervous system development
Cytokine signaling
Infectious disease
-log10 adjusted p-value
Normalized Enrichment Score
B
IL-1R pathway (Biocarta)
Module score
Max
Min
C
IL-1R Pathway (Biocarta)
Time (days)
0 2 7 14
Map3k14
Ikbkb
Il6
Irak1
Myd88
Nfkbia
Rela
Tgfb3
Map3k1
Tgfb1
Nfkb1
Irak3
Jun
Tgfb2
Traf6
Il1r1
Il1rn
Tnf
Il1b
Mapk8
Chuk
Map3k7
Tab1
Map2k3
Il1a
Irak2
Map2k6
Il1rap
Mapk14
Min Max
Row max expression (logn)
D
Relative Il1r1 mRNA
Relative C3 mRNA
Relative Cfb mRNA
IL-1β:
gRNA:
Ctrl.
Il1r1

### Figure S8

**A**

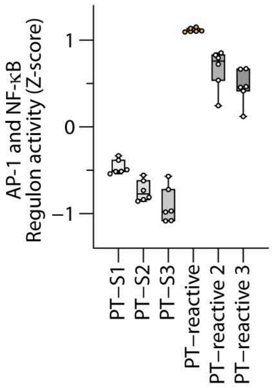

AP-1 and NF-κB
Regulon activity (Z-score)
1
0
−1
PT−S1
PT−S2
PT−S3
PT−reactive
PT−reactive 2
PT−reactive 3


**B**

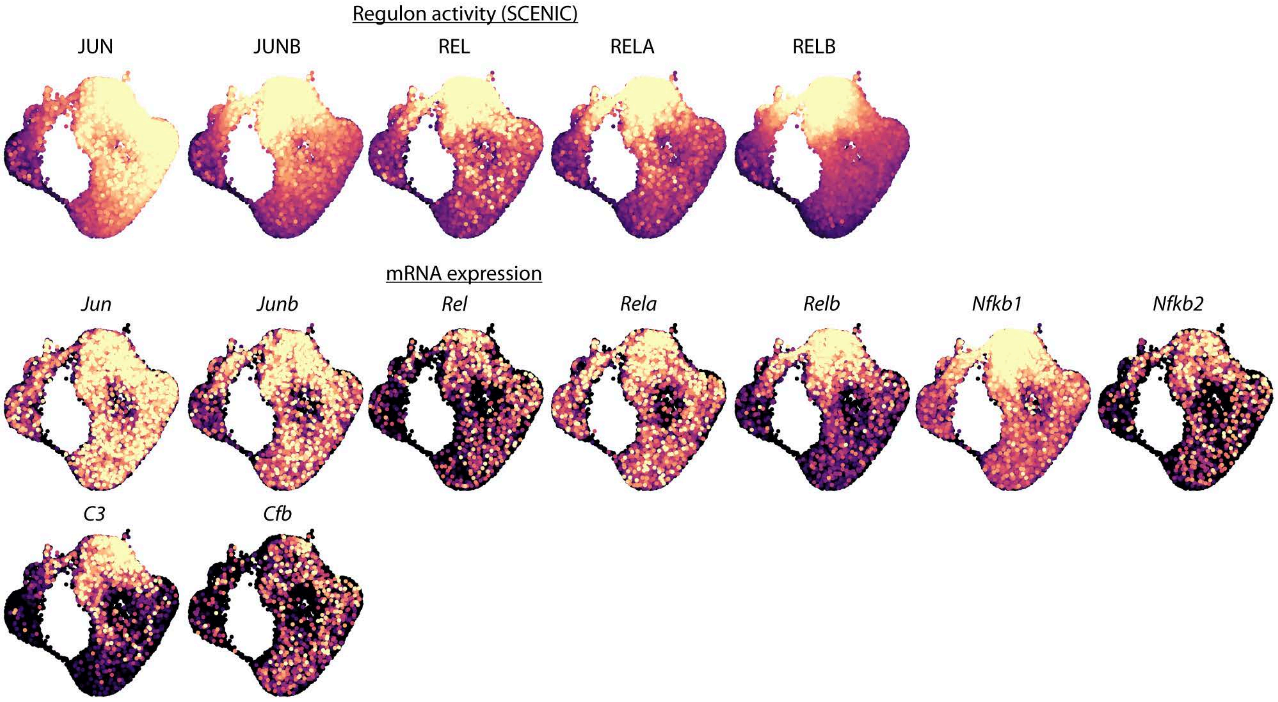

Regulon activity (SCENIC)
JUN
JUNB
REL
RELA
RELB
mRNA expression
Jun
Junb
Rel
Rela
Relb
Nfkb1
Nfkb2
C3
Cfb

### Figure S9

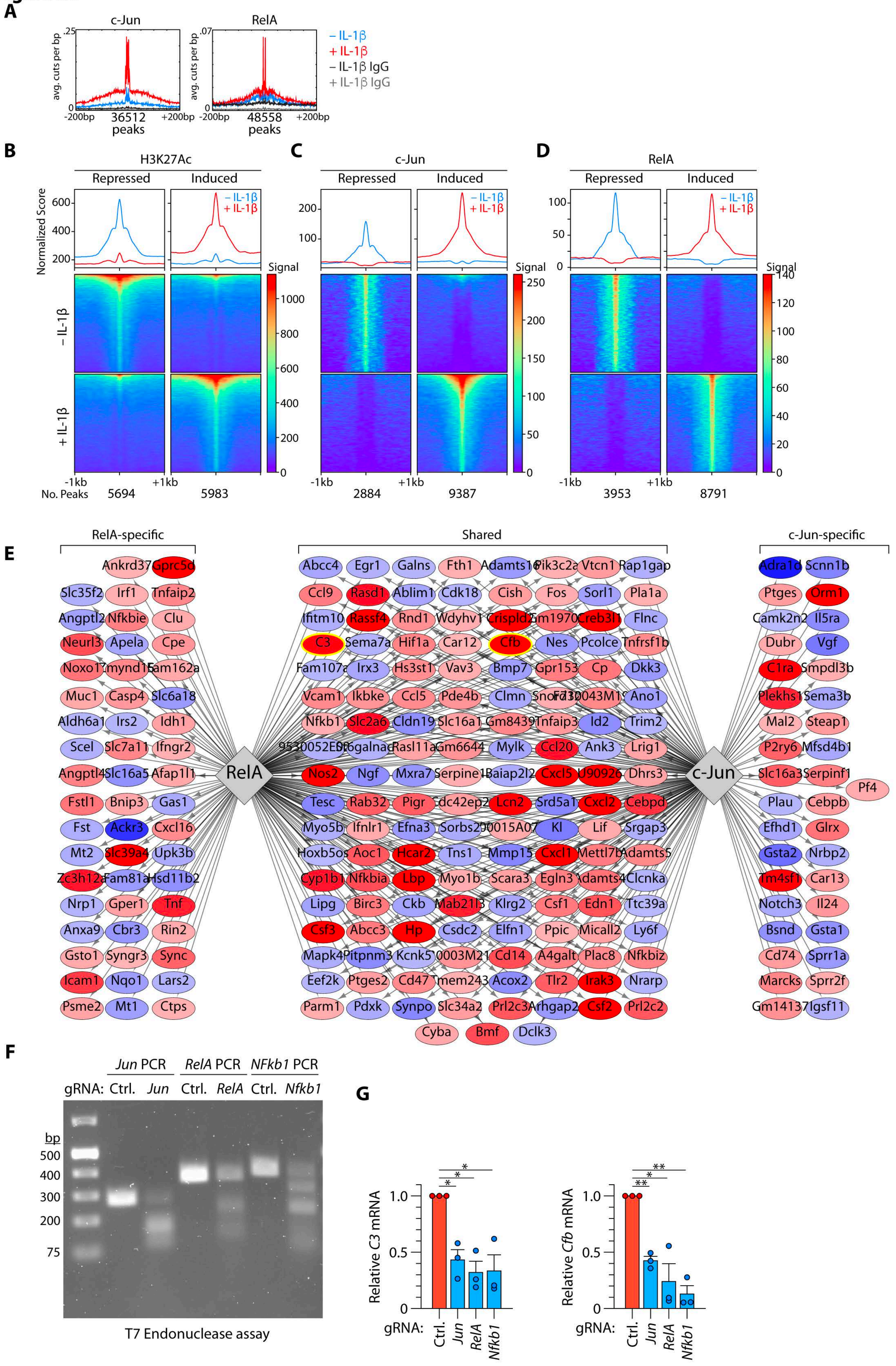
A
c-Jun
RelA
– IL-1β
+ IL-1β
– IL-1β IgG
+ IL-1β IgG
avg. cuts per bp
36512 peaks
48558 peaks
-200bp
+200bp
B
H3K27Ac
C
c-Jun
D
RelA
Repressed
Induced
Normalized Score
Signal
– IL-1β
+ IL-1β
-1kb
+1kb
No. Peaks
5694
5983
2884
9387
3953
8791
E
RelA-specific
Shared
c-Jun-specific
RelA
c-Jun
F
Jun PCR
RelA PCR
NFkb1 PCR
gRNA: Ctrl. Jun Ctrl. RelA Ctrl. Nfkb1
bp
500
400
300
200
75
T7 Endonuclease assay
G
Relative C3 mRNA
Relative Cfb mRNA
gRNA:
Ctrl.
Jun
RelA
Nfkb1

### Figure S10

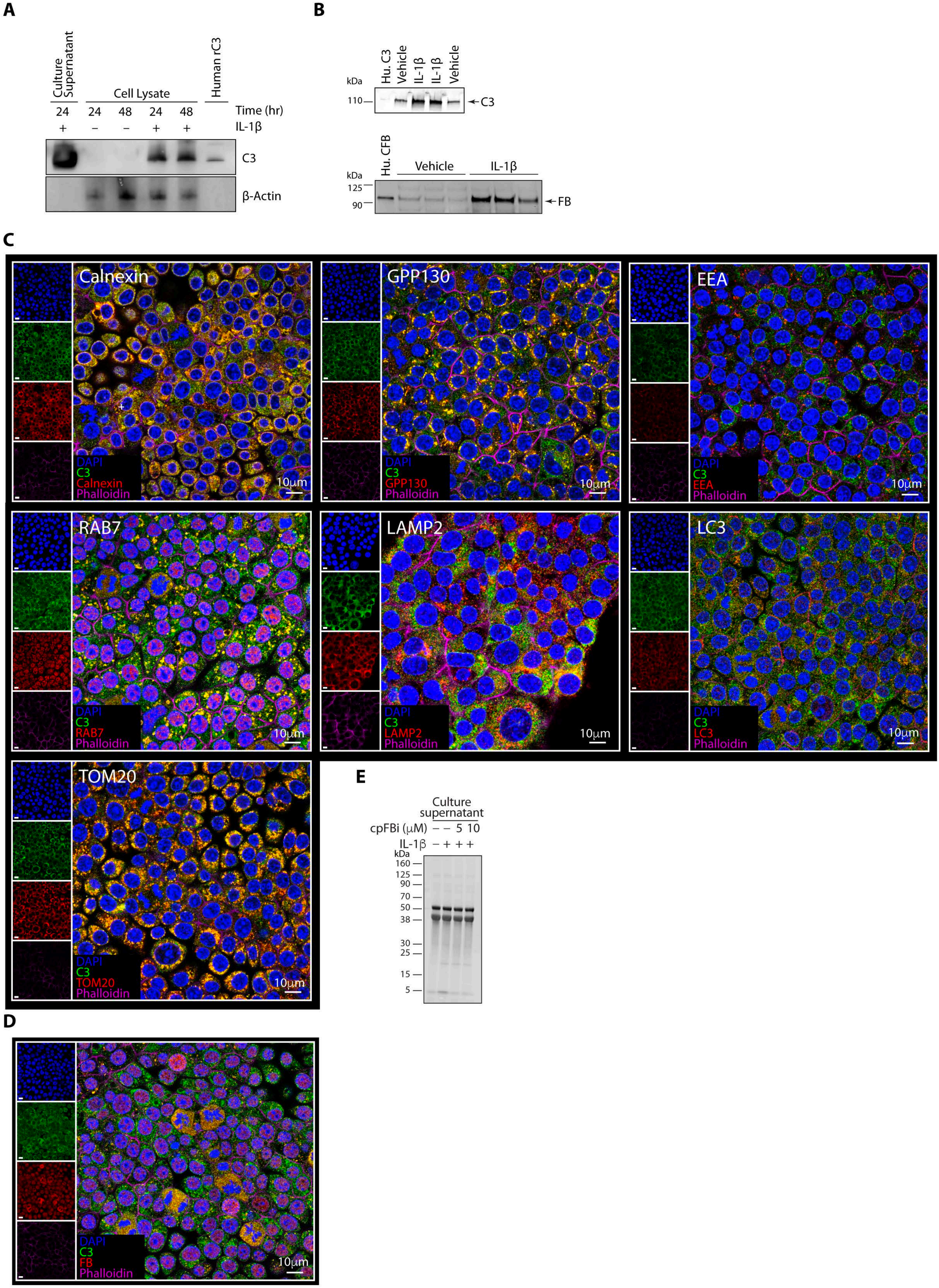
A
Culture Supernatant
Cell Lysate
Human rC3
24 24 48 24 48 Time (hr)
+ – – + + IL-1β
C3
β-Actin
B
kDa
Hu. C3
Vehicle
IL-1β
IL-1β
Vehicle
110
← C3
Hu. CFB
Vehicle
IL-1β
125
90
← FB
C
Calnexin
GPP130
EEA
RAB7
LAMP2
LC3
TOM20
DAPI
C3
Phalloidin
10μm
D
DAPI
C3
FB
Phalloidin
10μm
E
Culture supernatant
cpFBi (μM) – – 5 10
IL-1β – + + +
kDa
160
125
90
70
50
38
30
25
15
5

#### Figure S11

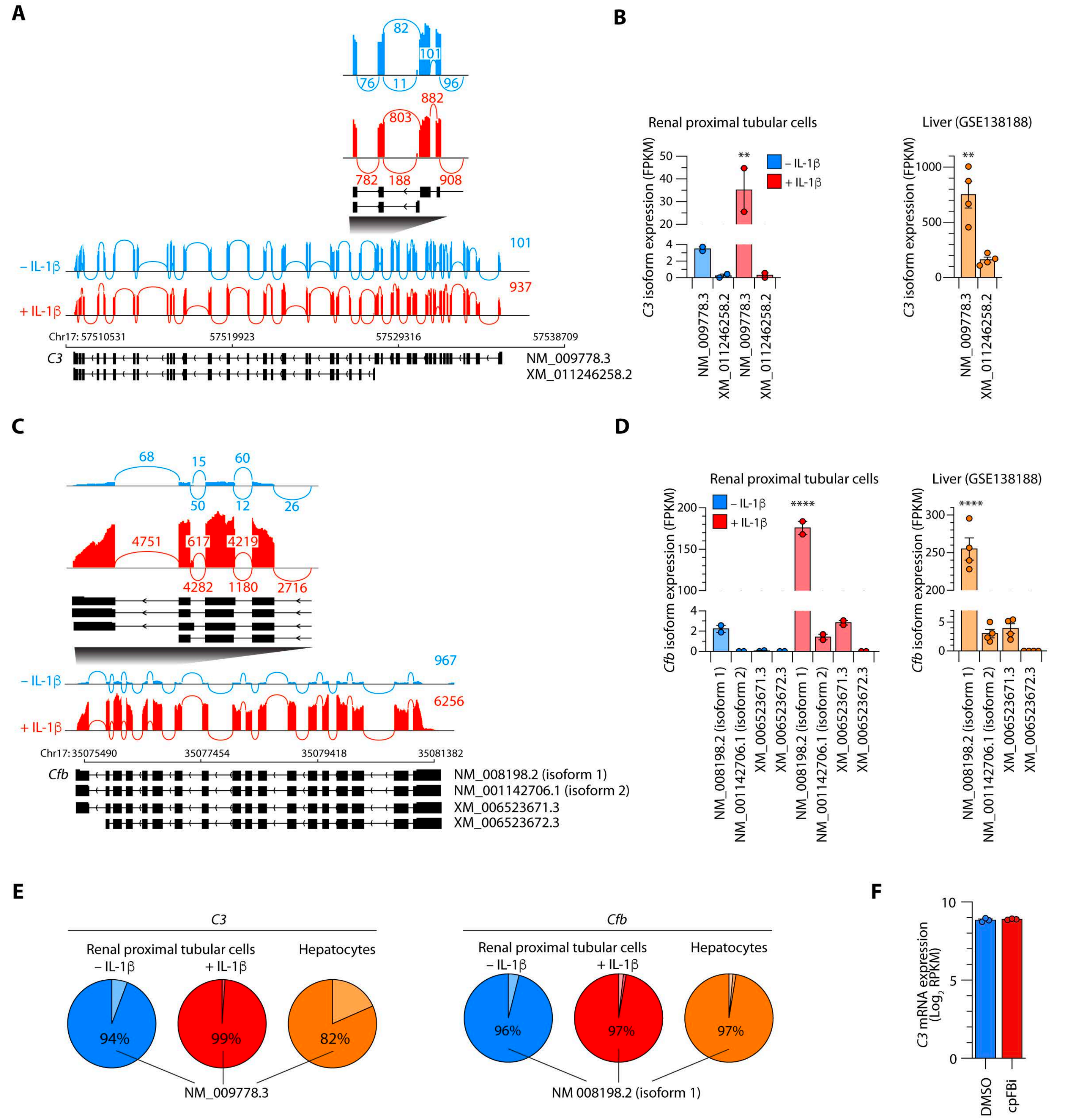

A
82
101
76
11
96
882
803
782
188
908
101
937
– IL-1β
+ IL-1β
Chr17: 57510531
57519923
57529316
57538709
C3
NM_009778.3
XM_011246258.2
B
Renal proximal tubular cells
Liver (GSE138188)
C3 isoform expression (FPKM)
– IL-1β
+ IL-1β
**
NM_009778.3
XM_011246258.2
C
68
15
60
50
12
26
4751
617
4219
4282
1180
2716
967
6256
Chr17: 35075490
35077454
35079418
35081382
Cfb
NM_008198.2 (isoform 1)
NM_001142706.1 (isoform 2)
XM_006523671.3
XM_006523672.3
D
Renal proximal tubular cells
Liver (GSE138188)
Cfb isoform expression (FPKM)
****
E
C3
Cfb
Renal proximal tubular cells
Hepatocytes
94%
99%
82%
96%
97%
97%
NM_009778.3
NM 008198.2 (isoform 1)
F
C3 mRNA expression (Log2 RPKM)
DMSO
cpFBi

### Figure S12

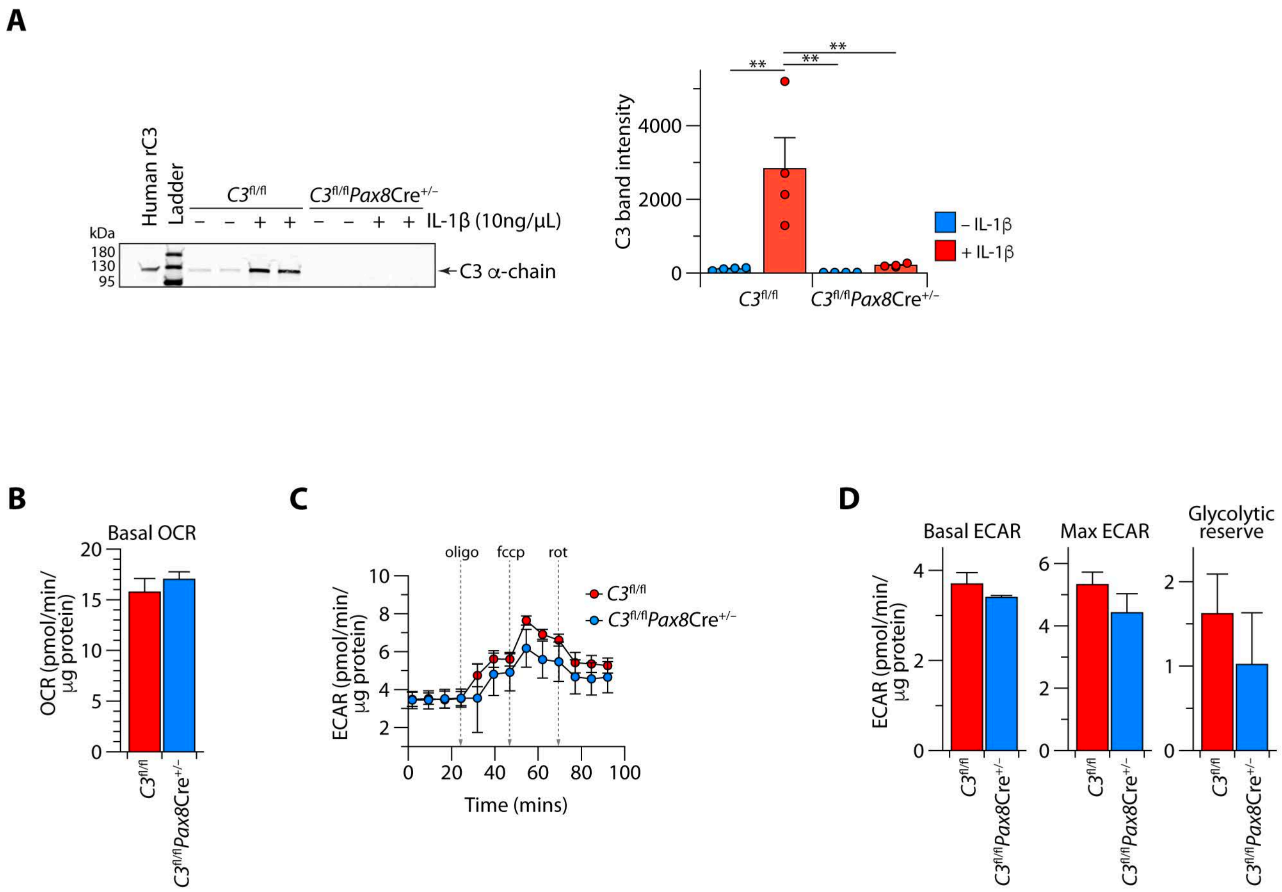
A
Human rC3
Ladder
C3fl/fl
C3fl/flPax8Cre+/−
− − + + − − + + IL-1β (10ng/μL)
kDa
180
130
95
C3 α-chain
C3 band intensity
4000
2000
0
**
**
**
– IL-1β
+ IL-1β
C3fl/fl
C3fl/flPax8Cre+/−
B
Basal OCR
OCR (pmol/min/ μg protein)
20
15
10
5
0
C3fl/fl
C3fl/flPax8Cre+/−
C
oligo
fccp
rot
ECAR (pmol/min/ μg protein)
10
8
6
4
2
0
20
40
60
80
100
Time (mins)
C3fl/fl
C3fl/flPax8Cre+/−
D
Basal ECAR
Max ECAR
Glycolytic reserve
ECAR (pmol/min/ μg protein)
4
2
0
6
4
2
0
2
1
0
C3fl/fl
C3fl/flPax8Cre+/−

**Figure S13**

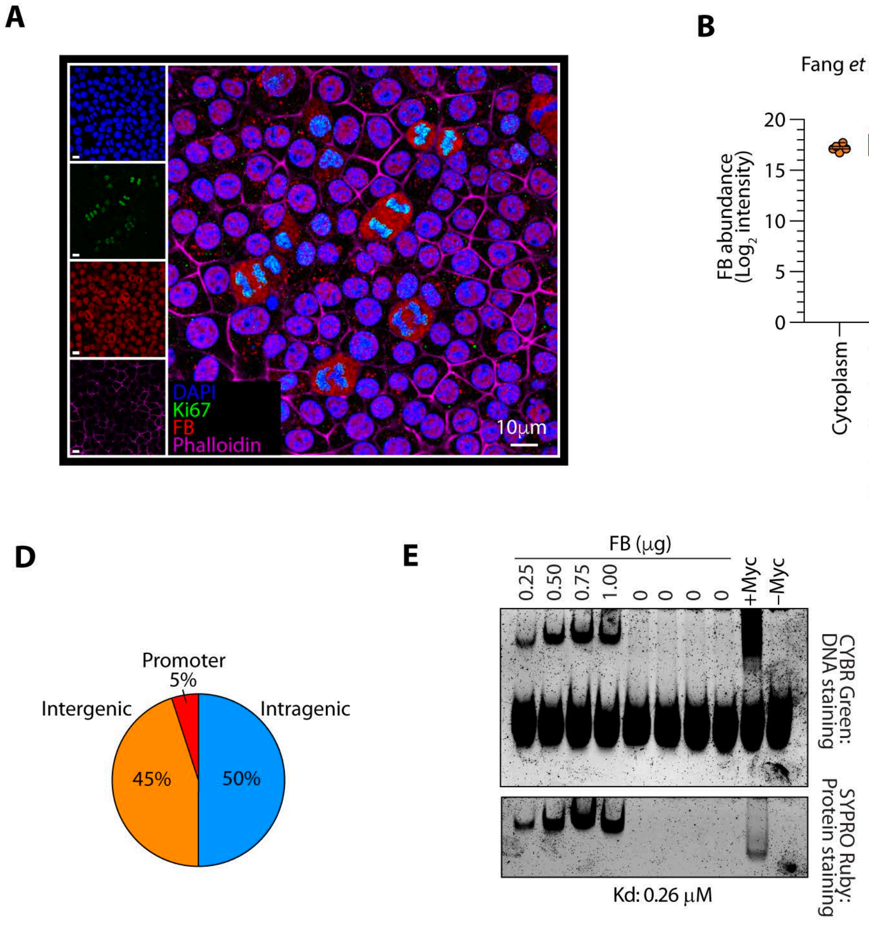
A
DAPI
Ki67
FB
Phalloidin
10μm
B
Fang et al., 2025
FB abundance (Log2 intensity)
20
15
10
5
0
Cytoplasm
Nuclear fraction A
Nuclear fraction B
C
Eslami et al., 2026
FB abundance (VSN-normalized intensity)
20
10
0
Cytoplasm
Nucleus
D
Promoter
5%
Intergenic
Intragenic
45%
50%
E
FB (μg)
0.25
0.50
0.75
1.00
0
0
0
0
+Myc
−Myc
CYBR Green: DNA staining
SYPRO Ruby: Protein staining
Kd: 0.26 μM

### Figure S14

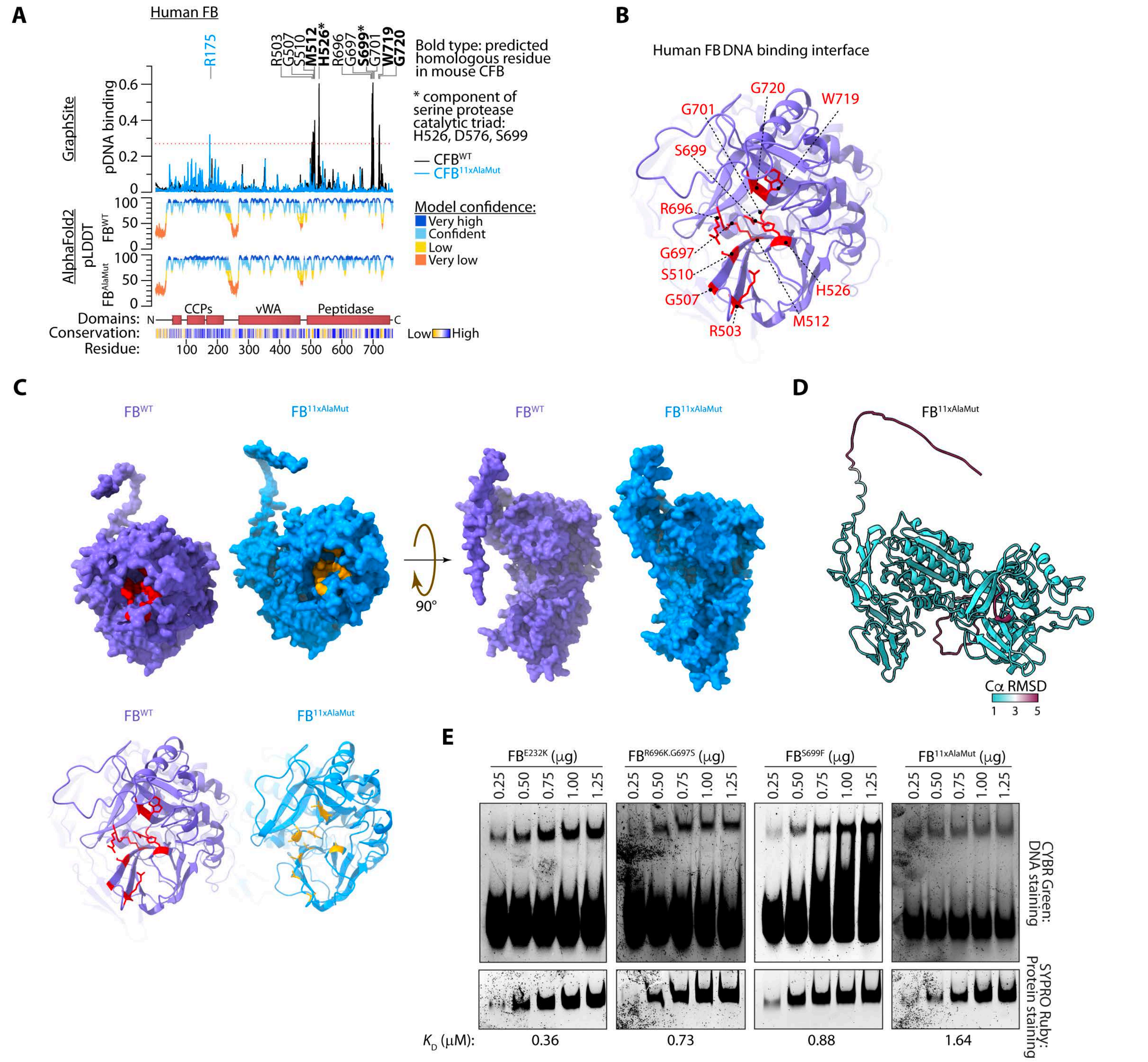
A
Human FB
R175
R503
G507
S510
M512
H526*
R696
G697
S699*
G701
W719
G720
Bold type: predicted homologous residue in mouse CFB
* component of serine protease catalytic triad: H526, D576, S699
— CFB^WT
— CFB^11xAlaMut
GraphSite
pDNA binding
Model confidence:
Very high
Confident
Low
Very low
AlphaFold2
pLDDT
FB^WT
FB^AlaMut
Domains:
CCPs
vWA
Peptidase
Conservation:
Low
High
Residue:
B
Human FB DNA binding interface
C
FB^WT
FB^11xAlaMut
90°
D
FB^11xAlaMut
Cα RMSD
E
FB^E232K (μg)
FB^R696K.G697S (μg)
FB^S699F (μg)
FB^11xAlaMut (μg)
0.25
0.50
0.75
1.00
1.25
CYBR Green: DNA staining
SYPRO Ruby: Protein staining
K_D (μM):
0.36
0.73
0.88
1.64

### Figure S15

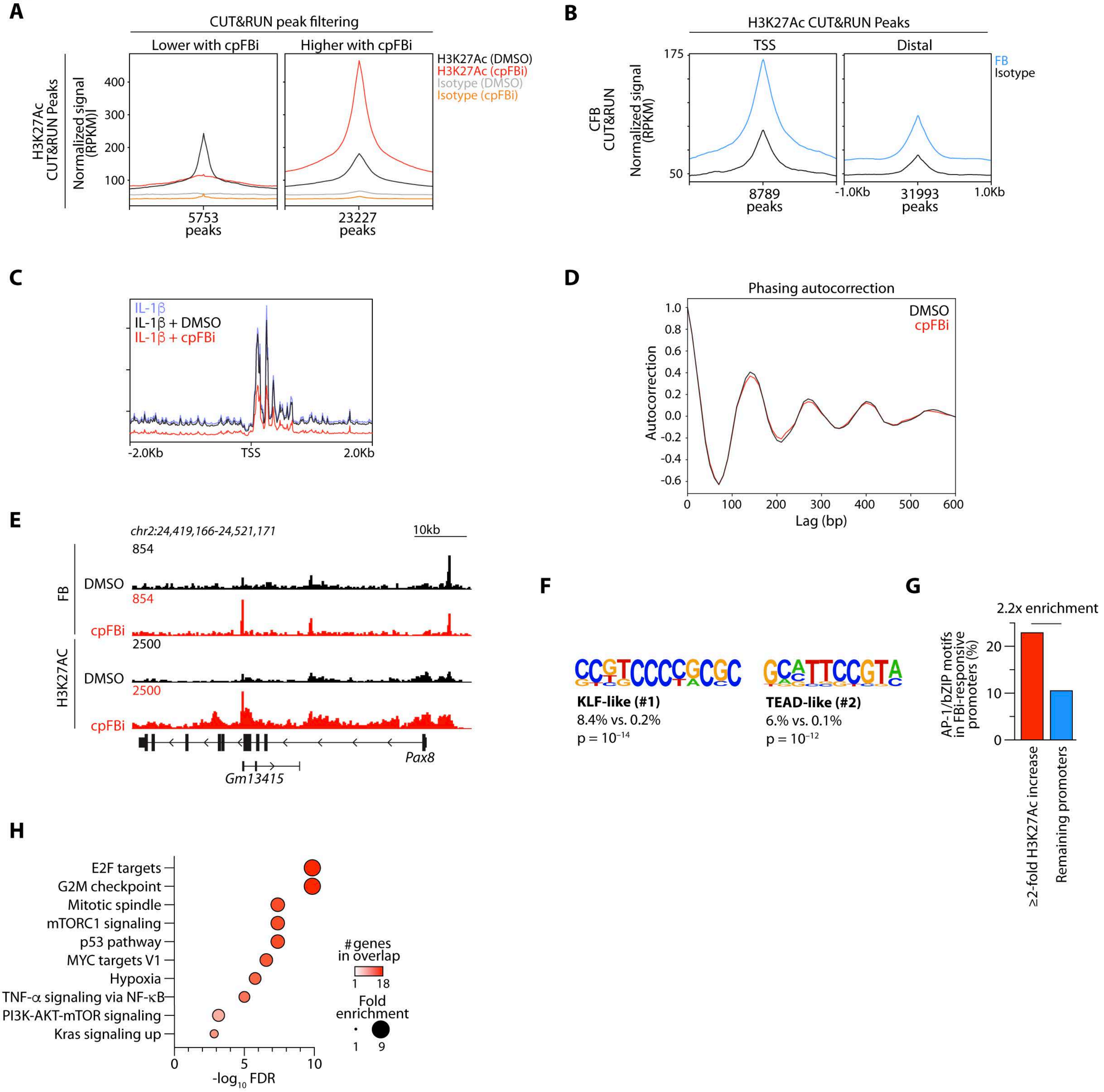
A
CUT&RUN peak filtering
Lower with cpFBi
Higher with cpFBi
H3K27Ac (DMSO)
H3K27Ac (cpFBi)
Isotype (DMSO)
Isotype (cpFBi)
H3K27Ac CUT&RUN Peaks
Normalized signal (RPKM)]
400
300
200
100
5753 peaks
23227 peaks
B
H3K27Ac CUT&RUN Peaks
TSS
Distal
FB
Isotype
CFB CUT&RUN
Normalized signal (RPKM)
175
50
8789 peaks
-1.0Kb
31993 peaks
1.0Kb
C
IL-1β
IL-1β + DMSO
IL-1β + cpFBi
-2.0Kb
TSS
2.0Kb
D
Phasing autocorrection
DMSO
cpFBi
Autocorrection
1.0
0.8
0.6
0.4
0.2
0.0
-0.2
-0.4
-0.6
0
100
200
300
400
500
600
Lag (bp)
E
chr2:24,419,166-24,521,171
10kb
FB
DMSO
854
cpFBi
854
H3K27AC
DMSO
2500
cpFBi
2500
Pax8
Gm13415
F
KLF-like (#1)
8.4% vs. 0.2%
p = 10^-14
TEAD-like (#2)
6.% vs. 0.1%
p = 10^-12
G
2.2x enrichment
AP-1/bZIP motifs in FBi-responsive promoters (%)
20
10
0
≥2-fold H3K27Ac increase
Remaining promoters
H
E2F targets
G2M checkpoint
Mitotic spindle
mTORC1 signaling
p53 pathway
MYC targets V1
Hypoxia
TNF-α signaling via NF-κB
PI3K-AKT-mTOR signaling
Kras signaling up
#genes in overlap
1
18
Fold enrichment
1
9
0
5
10
-log10 FDR

### Figure S16

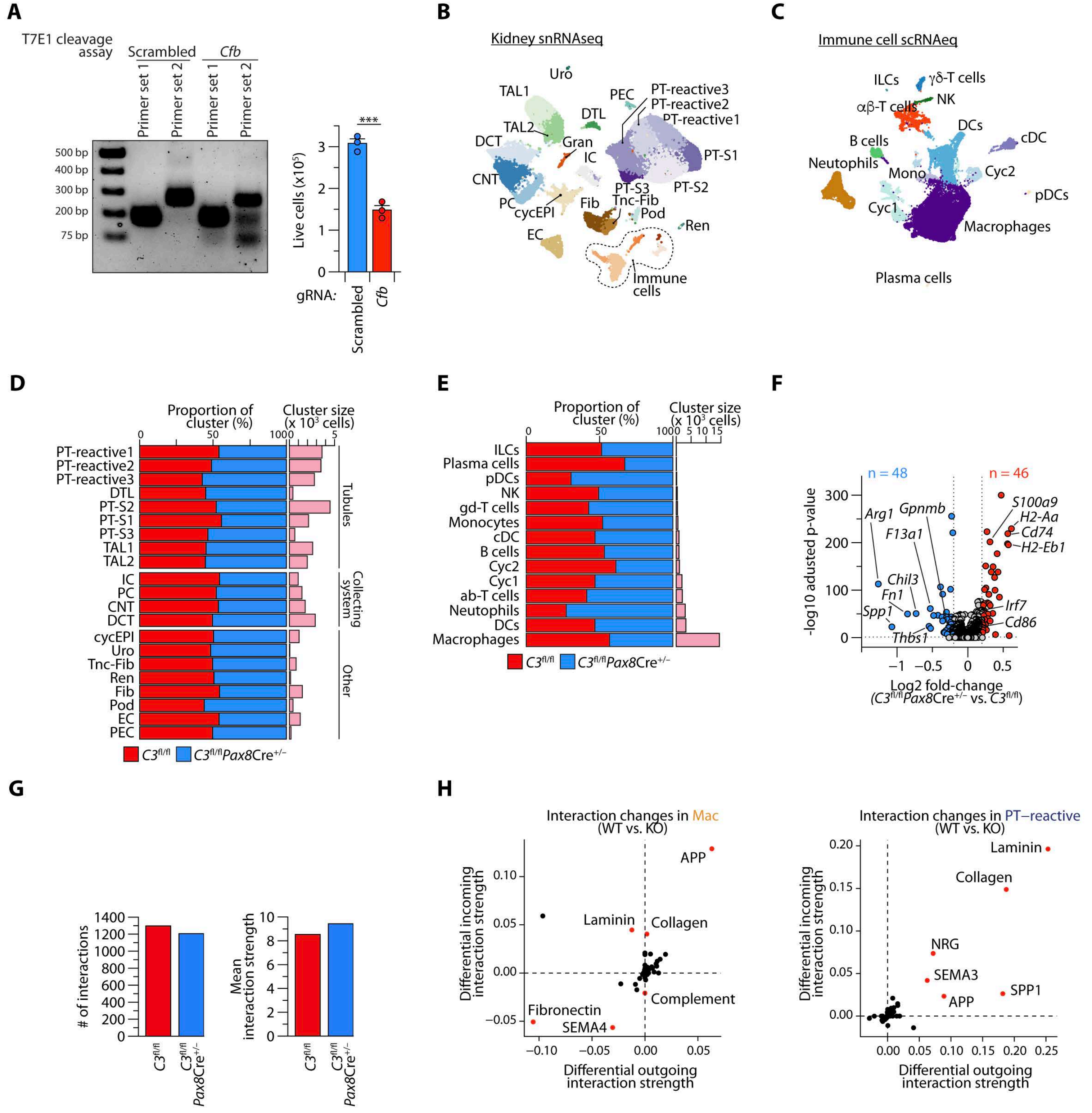
A
T7E1 cleavage assay
Scrambled
Cfb
Primer set 1
Primer set 2
Primer set 1
Primer set 2
500 bp
400 bp
300 bp
200 bp
75 bp
***
Live cells (x10^5)
gRNA:
Scrambled
Cfb
B
Kidney snRNAseq
Uro
TAL1
PEC
PT-reactive3
PT-reactive2
PT-reactive1
DTL
TAL2
Gran
DCT
IC
PT-S1
CNT
PT-S3
PT-S2
PC
cycEPI
Fib
Tnc-Fib
Pod
Ren
EC
Immune cells
C
Immune cell scRNAeq
ILCs
γδ-T cells
NK
αβ-T cells
DCs
B cells
cDC
Neutrophils
Mono
Cyc2
pDCs
Cyc1
Macrophages
Plasma cells
D
Proportion of cluster (%)
Cluster size (x 10^3 cells)
PT-reactive1
PT-reactive2
PT-reactive3
DTL
PT-S2
PT-S1
PT-S3
TAL1
TAL2
IC
PC
CNT
DCT
cycEPI
Uro
Tnc-Fib
Ren
Fib
Pod
EC
PEC
Tubules
Collecting system
Other
C3fl/fl
C3fl/fl Pax8Cre+/−
E
Proportion of cluster (%)
Cluster size (x 10^3 cells)
ILCs
Plasma cells
pDCs
NK
gd-T cells
Monocytes
cDC
B cells
Cyc2
Cyc1
ab-T cells
Neutrophils
DCs
Macrophages
C3fl/fl
C3fl/fl Pax8Cre+/−
F
n = 48
n = 46
-log10 adusted p-value
Arg1
Gpnmb
F13a1
Chil3
Fn1
Spp1
Thbs1
S100a9
H2-Aa
Cd74
H2-Eb1
Irf7
Cd86
Log2 fold-change
(C3fl/fl Pax8Cre+/− vs. C3fl/fl)
G
### of interactions
Mean interaction strength
C3fl/fl
C3fl/fl Pax8Cre+/−
H
Interaction changes in Mac
(WT vs. KO)
APP
Laminin
Collagen
Complement
Fibronectin
SEMA4
Interaction changes in PT−reactive
(WT vs. KO)
Laminin
Collagen
NRG
SEMA3
APP
SPP1
Differential incoming interaction strength
Differential outgoing interaction strength

### Figure S17

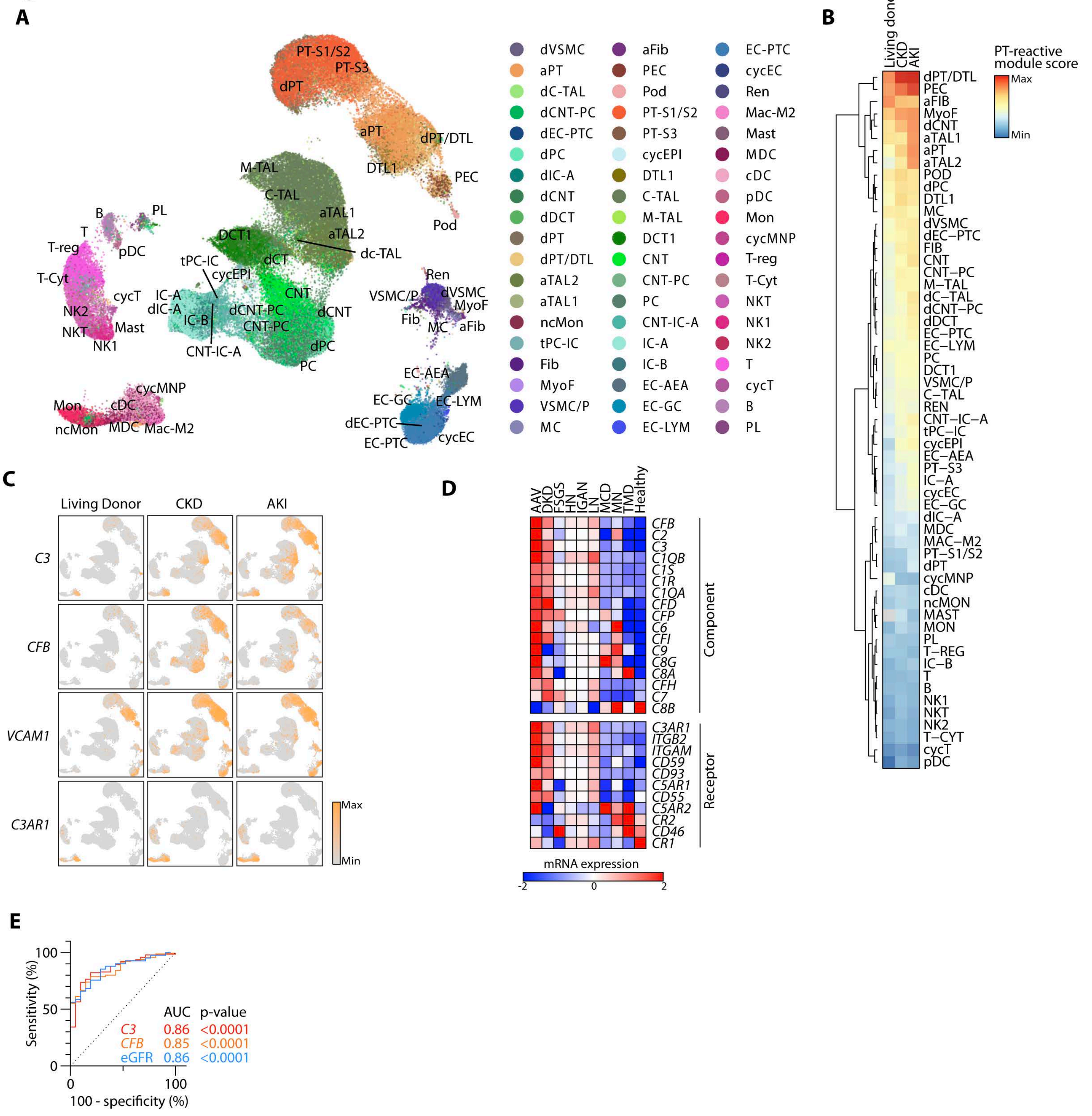
A
B
Living donor
CKD
AKI
PT-reactive module score
Max
Min
C
Living Donor
CKD
AKI
C3
CFB
VCAM1
C3AR1
D
AAV
DKD
FSGS
HN
IGAN
LN
MCD
MN
TMD
Healthy
Component
Receptor
mRNA expression
E
Sensitivity (%)
100 - specificity (%)
AUC p-value
C3 0.86 <0.0001
CFB 0.85 <0.0001
eGFR 0.86 <0.0001
